# EASYstrata: An All-in-One Workflow for Genome Annotation and Genomic Divergence Analysis

**DOI:** 10.1101/2025.01.06.631483

**Authors:** Quentin Rougemont, Elise Lucotte, Loreleï Boyer, Alexandra Jalaber de Dinechin, Alodie Snirc, Tatiana Giraud, Ricardo C. Rodríguez de la Vega

**Affiliations:** Ecologie Systématique et Evolution, CNRS, Universite Paris-Saclay, AgroParisTech, 91 198 Gif-sur-Yvette, France

## Abstract

New reference genomes and transcriptomes are increasingly available across the tree of life, opening new avenues to tackle exciting questions. However, there are still challenges associated with annotating genomes and inferring evolutionary processes and with a lack of methodological standardisation. Here, we propose a new workflow designed for evolutionary analyses to overcome these challenges, facilitating the detection of recombination suppression and its consequences in terms of rearrangements and transposable element accumulation. To do so, we assemble multiple bioinformatic steps in a single easy-to-use workflow. We combine state-of-the-art tools to detect transposable elements, annotate genomes, infer gene orthology relationships, compute divergence between sequences, infer evolutionary strata (i.e., footprints of stepwise extension of recombination suppression) and their structural rearrangements, and visualise the results. This workflow, called EASYstrata, was applied to reannotate 42 published genomes from *Microbotryum* fungi. We show in further case examples from a plant and an animal that we recover the same strata as previously described. While this tool was developed with the goal to infer divergence between sex or mating-type chromosomes, it can be applied to any pair of haplotypes whose patteern of divergence is of interest. This workflow will facilitate the study of non-model species for which newly sequenced phased diploid genomes are becoming available.

## Introduction

The past ten years have seen an increasing number of available high-quality genomes across a wide range of organisms, including many non-model species [1]. This opens up the possibility to better understand the evolution of genomes across the tree of life. For instance, it is now possible to study the patterns of recombination suppression along sex chromosomes [2] and to empirically test the theoretical hypotheses developed to explain these patteerns [3]. In particular, recombination suppression often progressively extends along sex chromosomes and other supergenes [4]. Such stepwise extension of recombination suppression can be inferred in genomes by detecting patterns of evolutionary strata, i.e., discontinuous stair-like patterns of synonymous divergence between sex chromosomes or mating-type chromosomes [5,6], Fig 1). Indeed, synonymous substitutions regularly accumulate with time between alleles on alternative sex chromosomes from the onset of recombination suppression, making synonymous divergence a proxy for the time since recombination suppression. For detecting stepwise extension of recombination suppression, synonymous divergence needs to be plotted against the ancestral gene order, because rearrangements following recombination suppression reshuffle gene order, therefore erasing footprints of evolutionary strata. The proxy for ancestral gene order is typically the sex chromosome from the homogametic sex (X or Z) in XY and ZW systems. In UV-like systems, in which neither of the two sex or mating-type chromosomes recombine, both accumulating rearrangements, an outgroup with recombining sex-related chromosomes can be used [2,6]. The evolutionary and proximal hypotheses for explaining stepwise extension of recombination suppression can be tested by investigating the degrees of rearrangements between sex chromosomes and the distribution of deleterious mutations [7].

**Figure 1.**
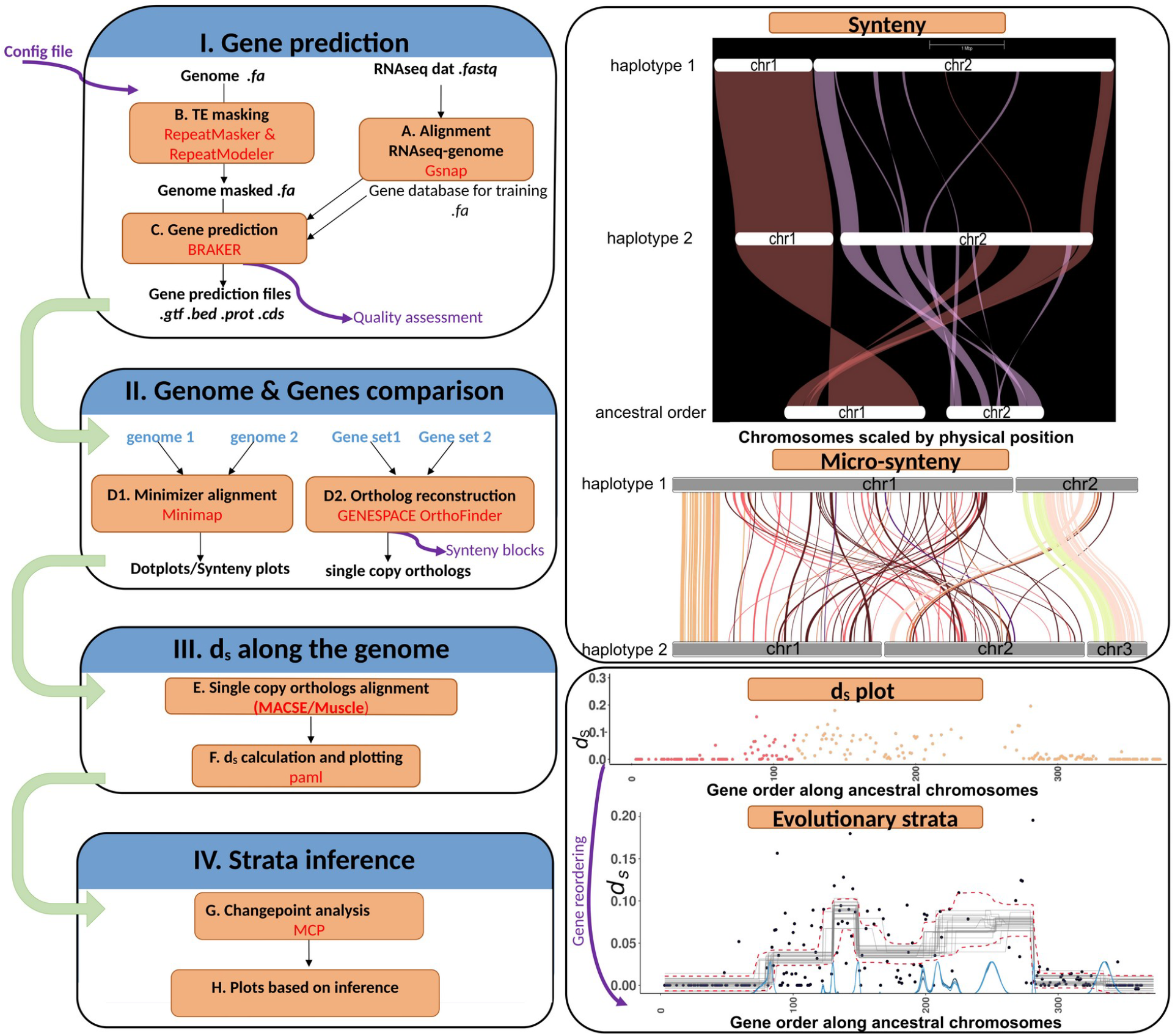
Overview of the workflow used for haplotype alignment and visualisation, transposable element (TE) detection, gene prediction, synteny analysis, per-gene synonymous divergence (d _S_) and changepoint analysis. A list of all third-party tools is provided in Table S1. Details of how to install the workflow are provided in the github INSTALL.md file.

However, the large amount of genomic data generates multiple challenges for analyses. While standardised procedures are now becoming available for genome assemblies [8], this is not yet the case for gene prediction. Indeed, analysing genomes annotated with different gene calling methods may introduce biases in downstream inference [9], for example if a method systematically predicts more genes than another method. In addition, the computation of synonymous divergence may be difficult given that the dedicated tools possess a steep learning curve [10]. Identifying and objectively delimiting evolutionary strata is often not straightforward either, given the stochastic noise in synonymous divergence values. For all these reasons, inference of evolutionary strata in sex and mating-type chromosomes can be a considerable challenge, while being crucial to our understanding of the evolution of sex chromosomes and other supergenes.

Easy-to-use, reproducible and efficient workflows are still lacking in this research area, precluding routine analyses of evolutionary strata and limiting the reanalysis of existing data. There is an increasing need for reproducible and standardised workflows to enable comparison across multiple organisms. Here, we provide a reproducible workflow for i) transposable element (TE) and gene calling, ii) gene filtration, quality assessment and subsequent inference of orthology relationships, iii) synteny pattern inferences based on both whole-genome and gene-based comparisons, and iv) synonymous divergence visualisation and evolutionary stratum inferences, as well as assessment of their association with chromosomal rearrangements. We called the workflow “EASYstrata” for Evolutionary analysis with Ancestral SYnteny for strata identification. EASYstrata is fully available at https://github.com/QueentinRougemont/EASYstrata, and can be deployed on any cluster. Below, we describe its difflerent modules, including options depending on the availability of ancestral gene rank and of gene annotations. The pipeline has been used previously for studying the giant sex chromosomes of *Silene latifolia* [11], and we describe here its features and its application to a set of 42 previously published genomes in *Microbotryum* anther-smut fungi, focusing on two study cases among them. In this group of fungi, there were ancestrally two mating-type chromosomes, bearing each a mating-type locus (HD and PR loci, for homeodomain and pheromone receptor genes, respectively), and with two alleles at the PR mating-type locus (called a _1_ and a_2_). The species *M. lagerheimii* has been shown to be a good proxy with the ancestral gene order of these two mating-type chromosomes [6]. We generated here a new reference genome for this species (Supp Methods and Results). Across the *Microbotryum* genus, there have been multiple events of mating-type chromosome rearrangements bringing PR and HD loci on the same chromosome, and recombination suppression linking these two loci. Recombination suppression has then extended away from the mating-type loci in successive steps independently in several species [12]. By using EASYstrata, we show in case examples that we recover the same strata as previously described. We also recovered the strata previously inferred in the threespine stickleback (*Gasterosteus aculeatus*; [13]). With this tool, we hope to stimulate research on the inference of evolutionary strata along sex chromosomes, mating-type chromosomes and other supergenes across a broad spectrum of organisms.

## Methods

### Input data and file configuration

Three data sets are typically needed in the workflow (**Fig 1**): first, high-quality genome assemblies in fasta format. These can be in the form of two alternative mating-type chromosomes, a full diploid genome including a pair of sex chromosomes, such as X/Y or Z/W chromosomes, or any autosome bearing supergenes with two differentiated haplotypes. The two differentiated sex chromosomes or haplotypes are hereafter called haplotype1 and haplotype2. Second, the workflow needs a haplotype designated as a proxy for the ancestral gene order of the region of interest, in fasta format. This can be the X chromosome in XY systems [5] or the genome of a closely related species with recombining mating-type chromosomes when neither of the two haplotypes recombine and therefore both accumulate rearrangements, thus diverging from the ancestral gene order [2,6]. In the absence of a chromosome constituting a good proxy for the ancestral gene order, dedicated methods for ancestral gene order reconstruction may be used (e.g., Agora [14]). Even if no proxy for an ancestral state is available, at least one closely related species (outgroup) is recommended to help identify single-copy orthologs between haplotypes. Third, a set of protein sequences from as many closely related species as possible (in fasta format) can be provided, which will improve gene prediction in the genome annotation step with BRAKER [15]. If no such data are provided, the pipeline will use the pre-partitionned OrthoDB clades provided at https://bioinf.uni-greifswald.de/bioinf/partitioned_odb12/ [16]. In this case, a taxon name corresponding to the partition the species belong to should be provided (e.g., vertebrata, fungi, as detailed in the public repository). Optionally, RNAseq data (paired-end or single-end) can be used as input for the genome annotation step, in the form of a .txt file containing the list of files with RNAseq reads in fasta or fastq format.

The workflow takes as input a config file including the path to the files described above as well as additional information and external data. An overview of the input parameters is summarised in **Table 1**. Detailed examples with difflerent uses cases related to *Microbotryum* (examples 1-4,6), the white campion *Silene latifolia* (example 5), and the threespine stickleback *Gasterosteus aculeatus* (example 7) can be found in the examples page of the public repository (https://github.com/QueentinRougemont/EASYstrata/tree/main/example_data). Short names for the two haplotypes will be used as a basename to rename the fasta file of each haplotype genome, and the contig names within genomes, which will in turn serve as a basename ID in the genes name. Because the workflow relies on masking transposable elements prior to gene annotation using NCBI databases, a NCBI species name should be provided (NCBI species names can be accessed at https://www.ncbi.nlm.nih.gov/Taxonomy/taxonomyhome.html/index.cg). The evaluation of gene prediction completeness crucially relies on BUSCO (benchmarking universal single-copy orthologs [17]), i.e., evolutionarily-informed expectations of gene content for near-universal single-copy orthologs. A BUSCO lineage name is therefore also compulsory in the config file. In addition, the path to a .txt file containing the name of the contigs or chromosomes of interest in the haplotype1 (or ancestral chromosome), corresponding to the target sex chromosome, mating-type chromosome, supergene or any focal region of interest should be provided in the config file. Below, we detail each step that will be performed to annotate a given pair of genomes and compute per-gene synonymous divergence (summarised in **Table 1** and **Fig 1**). Our workflow depends on many third-party tools (detailed in Table S1). These tools were chosen based on the following criteria: 1) they are known to be well maintained by the community and to meet the standard for a high level of reproducibility, 2) they display high performance (e.g., ability to deal with complex and or large genome for BRAKER and RepeatMasker) and 3) they provide results of high quality (e.g., High BUSCO score for BRAKER and complete gene, consistent orthogroups for Orthofinder).

**Table 1.** Summary of the main compulsory and optional parameters to provide in the config file. If a single genome is provided, only the annotation step is performed. The orthoDBspecies value should be chosen in the following list: “Metazoa” “Vertebrata” “Viridiplantae” “Arthropoda” “Eukaryota” “Fungi” “Alveolata” and will be used for annotation if no additional data are available. With two genomes, the whole pipeline can be implemented. Alternatively, gtf/gff files may already be available, in which case only the latter stages of the workflow are executed. If no RNAseq data is available or if a bam file of aligned data is already available, the workflow will also be run accordingly. If a genome is available for the ancestral state, plots are drawn along the gene order in this genome. Without a proxy of the ancestral genome, plots are drawn along the gene order in haplotype1. More details available on the public repository.

| parameter | description |
| --- | --- |
| *compulsory parameters* |  |
| genome1 | full path to haplotype1 genome assembly (fasta, compressed or not) |
| haplotype1 | a short name to be used in basename of chromosome contigs (eg: haplo1) |
| *optional parameters* |  |
| genome2 | full path to haplotype2 genome assembly (fasta, compressed or not) |
| haplotype2 | a short name to be used in basename of chromosome contigs (eg: haplo2) |
| ancestral_genome | full path to ancestral genome (fasta, compressed or not) or to an outgroup |
| ancestral_gff | full path to ancestral gff or to an outgroup |
| *Annotation parameters* |  |
| annotate | A string (YES/NO); if NO or empty no annotation will be performed |
| fungus | A string (YES/NO); if NO or empty the fungus option is not provided to Braker |
| orthoDBspecies | A string stating the lineage to be used for orthoDB protein download (see |
|  | legend below), *compulsory* if RelatedProt is not provided |
| RelatedProt | Optional: full path to a set of external protein data (fasta format) |
| gtf1 | Optional: name of gtf1; only used if annotate = NO |
| gtf2 | Optional: name of gtf2; only used if annotate = NO |
| rnaseq | Optional: a string (YES/NO); if YES rnaseq data will be used |
| bam | Optional: full path to a list of bam files if rnaseq data are already aligned to the reference genome RNAseq option should still be set to YES so the bam will be used by BRAKER |
| RNAseqlist | Optional: full path to txt file listing rnaseq data (PE or SE) |
| *Transposable element TE parameters* |  |
| annotateTE | A string YES/NO if YES TE annotation is performed, if NO skip (assume TE already soft-masked from genome) |
| TE database | Full path to a custom TE database (if available) |
| ncbi_species | Compulsory: species name to be used for ncbi based de-novo TE discovery |
| *BUSCO compulsory parameters* |  |
| busco_lineage | BUSCO species name for your target species (can be obtained using Busco --list-datasets) |
| *chromosome/scaffold compulsory info* |  |
| scaffold | full path to a table listing the target genome to be used as reference, the region of interest, and whether chromosomes should be reversed ("R") or not ("N") |
| *Optional files for plotting* |  |
| TEgenome1 | Bed file of TE for genome1 (automatically constructed if annotateTE="YES") |
| TEgenome2 | Bed file of TE for genome1 (automatically constructed if annotateTE="YES") |
| TEancestral | Bed file of TE for the ancestral species |

### Transposable element detection

*De-novo* detection of transposable elements (TEs) is performed for each haplotype separately using RepeatModeler 2.0.2 [18]. Next, RepeatMasker [19] is used to improve TE detection and masking using TE databases. It is then possible to use one or several of the following: *i)* the *de-novo* families of TEs detected by RepeatModeler, and optionally *ii)* in-house TE libraries, and *iii)* RepBase database of TEs. The annotations labelled as “unknown” by RepeatModeler (i.e., without any known annotation) can be removed from the TE set to limit the rate of false positive TE identification that would correspond to genuine genes, or left as is, which may potentially induce an increased rate of false negative genes. If a softe-masked genome is already available, and no TE detection is needed, the option annotateTE = “NO” can be set directly in the config file, and this step will be skipped.

### Genome annotation and filtering

The resulting soft masked genome is then fed to BRAKER 3 [15], a method that combines GeneMark ETP [20] and AUGUSTUS [21]. While our pipeline has been optimised with BRAKER v2.1.6, we obtained similar results with BRAKER 3 and chose to focus on the latest version because it is the currently maintained version, and it does not require the license associated with GeneMark. We propose two use cases for Braker: *i*) with a set of protein sequences from closely related species only, or *ii*) with a set of protein sequences from closely related species and RNAseq data for the focal species.

In use case *i*, BRAKER is run for five successive (independent) rounds, in order to take into account (limited) stochasticity, and the round providing the highest BUSCO score is kept for downstream processing. In use case *ii*, the same steps as case *i* are applied to the protein sequences (BRAKER and selection of the best round). Additionally, raw RNAseq reads (paired end or single end) are trimmed with Trimmomatic [22] and mapped to the reference genome using Gsnap [23], with the resulting alignment being further sorted and filtered with Samtools [24]. A table of read count and mapping quality is produced at this step, together with plots of read depth along each assembled pseudomolecule. Then, BRAKER runs GeneMark-ETP and Augustus on mapped RNAseq data. The best protein sequence round, as evaluated with BUSCO (see below), and the RNAseq run are combined using TSEBRA [25]. While BRAKER 3 enables the use of both RNAseq and external data along with TSEBRA in a single command call, we found that calling BRAKER with each data type separately and combining with TSBERA afteerwards provided higher BUSCO scores (97% vs >98% on average); we therefore chose to stick with this approach. Moreover, this approach enables the users to more easily customize the TSEBRA config file. At this step, a report is automatically generated using BRAKER utilities, enabling the assessment of the number of complete and partial genes, the number of single- and multi-exon genes, the number of introns per gene and the number of genes fully or partially supported by external evidence (i.e., RNAseq and databases). Various histograms are also automatically produced, which may be exploited to further filter genes with aberrant profiles.

The resulting gtf file, which contains both Augustus-and GeneMark-based gene predictions is then modified with a custom perl script (obtained here: https://github.com/Gaius-Augustus/BRAKER/issues/457). The file is parsed in order to insert the haplotype name in each gene ID. This step facilitates the analyses of genes in a phylogenetic context, especially when multiple species are to be studied.

This gtf is then used to extract the CDS and protein sequences from the genome of each haplotype using gfflread [26]. Finally, data are filtered to keep a single transcript per gene, the longest one, of prime importance to accurately infer orthology. This constitutes the final gff, CDS and protein sequence files for which BUSCO scores are eventually computed. The quality of the gene prediction is further assessed automatically through blasts against the reviewed Swiss-Prot database available from uniprot. This enables the user to assess the proportion of predicted genes with at least one match against uniprot. Optionally, InterProScan will be run, combining several databases, such as pfam, panther, NCBIfam, cath-gene3D, in order to obtain prediction of the gene functions, family classification and domain predictions. The resulting hits, which are of interest in themselves for gene function inference, are also used to compute the proportion of genes with a hit against a database among all genes.

### Identifying rearrangements

#### Minimizer-based whole genome alignment

Alignments of the two haplotype sequences against each other and against the ancestral genome proxy are performed with Minimap2 [27] due to its rapidity and ease of use. Dotplots of primary alignments are automatically generated using PafR [28] and, if a list of sex-chromosome regions or supergenes of interest is provided in the config file, synteny plots are automatically drawn, to facilitate visual inspection of rearrangements.

#### Synteny analysis based on orthologous gene sets

Synteny is investigated based on the gene positions in the different haplotypes of orthologous genes or alleles. To that aim, the pipeline first infers gene orthology relationships using OrthoFinder [29]from the annotated gene set obtained from BRAKER. Instead of only extracting orthology information between single-copy genes, we took advantage of the recently developed GeneSpace pipeline [30], which combines Diamond v2.0.8 [31], OrthoFinder v2.5.4 and MCScanX [32], to integrate gene order and orthology relationships and detect gene collinearity (i.e., local synteny) between genomes. GeneSpace is run with default parameters and automatically produces dotplots among all pairs of genomes, as well as riparian plots based on gene order and gene physical position. If a set of target scaffolds (e.g., sex chromosomes or supergene haplotypes) are provided, specific subplots are also constructed. The results can easily be further processed *a posteriori* by the user, for instance by exploiting pangenome gene sets and customising riparian and dotplots (e.g., [11]).

The resulting single-copy orthologs extracted from OrthoFinder output files are further reformatted to serve as input to the Rideogram package [33], enabling the visualisation of gene order from the target pairs of scaffolds (sex chromosomes or supergene haplotypes). A Circos plot is also produced using circlize [34] from the resulting single-copy orthologs (i.e., one-to-one gametologs) highlighting the regions of interest. Optionally, bed files of the genes and transposable elements can be provided and will be plotted as inner tracks.

#### Calculating synonymous divergence between haplotypes

The set of single-copy orthologs identified by OrthoFinder is used as input on which the per-gene synonymous divergence (d_S_) and non-synonymous divergence (d_N_) are computed between haplotypes, as well as their standard errors (SE), using the yn00 program implemented in PAML [35]. A comparison between paml and codeml used in the pairwise mode revealed highly similar results (Fig S30); the pipeline computes both so that the user can compare the resulting estimates. To that aim, the CDS (coding sequences, fasta format) for each single-copy orthologous pair are aligned using MACSE 2.0.5 [36]. Aligning with MUSCLE [37], implemented in TranslatorX [38] yielded very similar results as with MACSE. All the processes involved are encompassed in a simple bash script. The per-gene synonymous divergence (d_S_) values are plotteed against the ancestral gene order with the goal to detect evolutionary strata. In cases where no proxy for the ancestral genome is available, the gene order along the haplotype 1 contigs is used to plot the per-gene synonymous divergence (d_S_) values. Figures are produced by either plotteing the d_S_ values along the genomic position or along the gene order. We defined the gene order as the order of the genes along the genome being taken as a proxy for the ancestral state, or along the haplotype1 (e.g., the X chromosome), genes being numbered from 1 to n, where n is the total number of genes along the X, thus accounting for the fact that many genes on the X will not have any ortholog on the Y. The rank thus does not include the information on the genomic position, which can render plots more readable, and can avoid biases in change-point analyses. In addition, this accounts for the fact that many genes on the haplotype1 will not have any ortholog on the second corresponding haplotype.

#### Visualising synonymous divergence (d_S_) along genomes and inferring evolutionary strata

The resulting per-gene d_S_ values are plotteed along the contigs of the genome representing the ancestral gene arrangement, using the genomic coordinates in base pairs (from the ancestral-like haplotype) and also using the gene order. The plots are performed in R with R packages listed in table S1.

The last step is to objectively infer the existence of evolutionary strata and delimit them, i.e., to detect discontinuous changes in d_S_ mean values along the ancestral gene order. This is performed using the Bayesian analysis implemented in the MCP R package [39]. If recombination suppression is a stepwise process, it is expected to result in difflerent mean d_S_ values (i.e., strata of *d*_*S*_, **Fig 2A**) in the different fragments that have successively stopped recombining (Charlesworth, 2010). Each stratum can be approximated by a linear model. In principle, different linear models for the distinct strata should fit better than a single model for the whole dataset, and changepoint location should correspond roughly to the boundaries of these strata (**Fig 2B**). Therefore, we implemented in the pipeline a series of models testing different events of recombination suppression, having generated evolutionary strata. Importantly, one-to-one gametologs need to be placed according to their order including all genes (i.e., also not one-to-one gametologs), as the relative spacing between genes can influence the change-point analysis. We implemented models with up to eight changepoints, which corresponds to six evolutionary strata if pseudo-autosomal (recombining) regions are present on each side of the non-recombining region. The Markov chain Monte Carlo (MCMC) function is run for 100,000 iterations across five replicates and 15,000 burn-in periods, and the results are automatically plotted for each model, including the intervals around the estimated changepoint (using qfit()). The most likely model is statistically chosen using leave-one-out cross-validation as implemented in the loo() function from the loo R package [40]. The estimated log-predictive density (ELPD) is computed for each model and models are compared using the loo_compare() function. Optionally, Bayesian hypothesis testing can be performed to compare the significance of mean d_S_ difflerences between all pairs of inferred strata, as implemented in the hypothesis() function. Bayes Factor (BF) > 5 is typically considered as good evidence of strong differences in mean d_S_ values. In addition, violin plots implemented in the ggstatsplots package [41] of d_S_ values in the difflerent strata are plotteed to further visualise the extent and significance of their differences. The coordinates of the strata are then used to visualise *a posteriori*, the distribution of d_S_ values along the genome, coloured by stratum, and to automatically redraw the ideogram, also coloured by stratum. For numbers of changepoints from 3 to 8, all the results will be automatically exported in a first pass analysis that returns i) a pdf file displaying the location of the changepoint, ii) a pdf file displaying the convergence of the chain, iii) a pdf file with violin plots of the d_S_ values for each stratum, iv) a pdf file with colored plot of the d_S_ values along the ancestral order v) a pdf file with colored plot of the d_S_ values along physical order, as well as vi) various text files displaying model weights, changepoint location and Bayes Factor testing the support for the difflerent strata.

**Figure 2.**
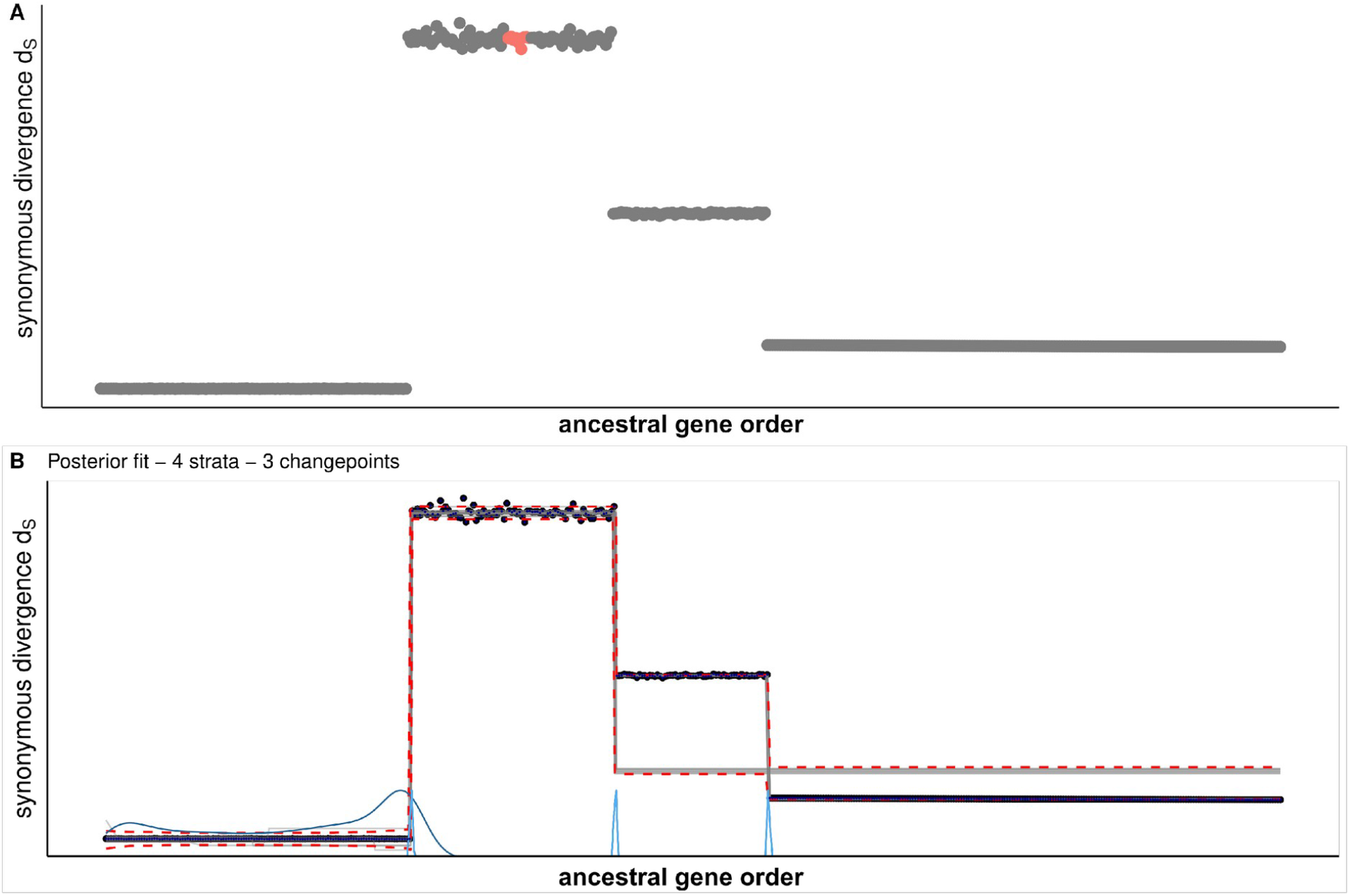
**A)** Expected patteern of d_S_ values of synonymous substitutions (d_S_) between alleles in the two sex chromosomes or haplotypes under a scenario with three successive events of recombination suppression, the first one leading to the formation of the stratum with the highest mean d _S_ value and containing the sex-determining locus (red dot) and the second one leading to a stratum with an intermediate level of synonymous divergence. **B)** Change-point location on these simulated data inferred with the MCP (multiple change point) R package in a Bayesian framework. The blue curves at the botteom of the x axis are the posterior distributions of the change point locations. The dashed red lines are the 2.5 and 97.5% quantiles of the fitteed (expected) values. The gray lines are 25 samples from the posterior values.

#### Running the workflow at difflerent stages of the process

Given the many different use cases of our workflow, we provide different options to run this pipeline (numbered from 1 to 7). While option 1 is used to perform a whole analysis, many users might only be interested in TE and gene prediction, available through option number 6, this can be useful if a single genome is to be annotated without any further analyses being required. Option number 2 will perform the TE, gene prediction, as well as GeneSpace and Synteny analyses. This can be useful, for instance, to help identifying sex chromosomes when no such information is available a priori. Option 3 is used to perform GeneSpace and synteny analyses as well as computing synonymous and non-synonymous divergence and infer evolutionary strata (plotteing d_S_ model comparison, Circos plots and Ideograms), which is useful if annotated genomes (gff and fasta files) are already available. Similarly, option 4 allows to compute d _S_ and infer evolutionary strata in cases where the synteny analysis was already performed. Option 5 is used to perform GeneSpace and synteny analyses only. Finally, option 7 will be the option of choice to perform model comparisons and draw associated graphs including ideograms of a posterior strata. This can be highly useful given that option 1 will perform a first-pass model comparison along the ‘default’ ancestral gene order (i.e., the order of genes along the genomic coordinate of the haplotype 1), which may not allow inferring strata if the ‘default’ gene order is not the ancestral gene order. Several user-crafteed modifications of the contig order may be required, or even deleting some contig fragments, for example if part of the ancestral sex or mating-type chromosome became autosomal in the studied species; typically, inferring strata will require visual inspection and biological interpretation from the GeneSpace riparian plots outputs, Circos plots, ideograms without priors and distribution of d_S_ values. These decision steps cannot be automated as they require interpretation.

## Results

### Application to a set of genomic data

We propose a new workflow that builds upon existing and state-of-the-art tools to detect evolutionary strata. Our approach includes three steps: 1) predicting genes and transposable elements, 2) comparing genes and genomes between haplotypes, and 3) inferring evolutionary strata through changepoint analyses of evolutionary divergence (d_S_). Each module of the workflow can be run separately in order to provide a high level of flexibility for different use cases: i) with all data (2 haplotypes, with or without an ancestral sequence, with or without RNAseq data, all analyses will be performed corresponding to option 1); ii) without information on sex chromosome, option 2 will perform only the repeat and gene annotation step as well as Genespace and synteny analyses; iii) with already annotated haplotype and ancestral-like sequence only the d_s_ computations, synteny and changepoint analyses will be performed, corresponding to option 3; iv) option 4 will perform the same step as option 3 except the Genespace analyses; v) option 5 will only perform the Genespace and synteny analyses, which can be useful for a previously annotated genome without information on sex chromosomes; vi) with only a single haplotype, only gene annotation will be performed, option 6); vii) to only perform a changepoint analysis use option 7, which can be useful afteer reordering the scafflold following a first-pass analysis; viii) to only draw plots afteer the d_S_ computation use the option 8.

This workflow is mainly intended toward people unfamiliar with bioinformatics, in order to facilitate each of these steps. Below, we briefly show a detailed application of the difflerent steps to a set of 42 published genomes of *Microbotryum* fungi. For the annotation part with different datasets, we focused on *Microbotryum lychnidis-dioicae*, one of the few species of the genus for which RNAseq data were available. For the second part (strata inference), we present results for *M. violaceum caroliniana*, a species with well assembled genomes and evolutionary strata of different ages, as well as additional examples from *Microbotryum* species in supplementary results. The inference of evolutionary strata took advantage of a newly assembled genome of *M. lagerheimii* (strain 129.01.A1) whose genome assembly details are presented in supplementary methods and **table S2**. Finally, we show in supplementary materials an example application to the fully automatic recovery of the strata already identified in the threespine stickleback (*Gasterosteus aculeatus*) genome [13] and we point the reader to [11] for an EASYstrata application to the giant sex chromosomes of the white campion *Silene latifolia* (detailed examples 5 and 7 in the public repository).

### Genome annotation

First, we performed on these case studies a classic transposable element (TE) detection that builds upon RepeatModeler and RepeatMasker tools. Optionally, an external database of existing TEs for the studied organisms can be provided. Next, we performed genome annotation with BRAKER V3 with either *i*) RNAseq data only, *ii*) custom external evidence only, *iii*) online external evidence from orthoDB for the target species only or *iv*) all options together. Results for the genome of the a _1_ mating type are presented in **Table 2** (**Table S4** for the a_2_ mating type). We implemented a filtration step to systematically keep the longest transcript, as many alternative transcripts are produced by BRAKER by default (**Table 2, Table S4**). Mapping rates are computed on the fly after running GSNAP and plots are automatically drawn (**Fig S1-S2**, mean read depth was 160 and 166 in the a_1_ and the a_2_ genomes, respectively, **Table S6**). Based on the longest transcript, the BUSCO results from the gene prediction tools all produced similar results. These results were in close agreement with the number of BUSCO genes predicted directly on the genome of the a _1_ mating type (i.e., C: 98.4% [S: 97.7%, D: 0.7%], F: 0.2%, M: 1.4%) (see **Table S3** for the genome description) and were slightly better than previously published annotation scores (*M. lychnidis-dioicae a*_*1*_: C: 96.3% [S: 95.7%, D: 0.6%], F: 1.4%, M: 2.3%; *M. lychnidis-dioicae a*_*2*_: C: 96.2% [S: 95.3%, D: 0.9%], F: 1.5%, M: 2.3%, n: 1764). The least accurate prediction scores were obtained when using only our custom database (built from previous annotations) (**Table 2**). Additional evaluation is automatically implemented in our workflow by running blast against the SwissProt database, enabling the users to further assess the proportion of genes with a blast and the length of the hits (**Table 2**). In addition, further annotation information can be obtained if the option to run InterProScan is set to “yes” in the config file (**Table 2**). While BUSCO enables an accurate assessment of the completeness of the gene prediction, it should be noted that BRAKER may fail to predict fast-evolving genes, even when they are well represented in a database. This is in particular the case of the pheromone receptor (PR) gene, which is of major interest in our focal species, controlling pre-mating fusion, and displaying several million-year old trans-specific polymorphism [42]. In the absence of RNAseq data, this important gene was in general not predicted when applying our pipeline across hundreds of genomes. Another issue pertains to the best way of combining RNAseq and external database (orthoDB11 and custom DB) in the final annotation with BRAKER. While we used TSEBRA following BRAKER’s recommendation, we found that the weight given to each evidence impacted very slightly the final BUSCO score. For instance, in our studied case in **Table 2**, we found that the combination of RNAseq with the two other databases (orthoDB11 and custom DB) actually resulted in a very small decrease in BUSCO accuracy, with six genes being missed in the combined annotation, as compared to results when using RNAseq only. Similarly, only four new genes were classified as fragmented compared to annotations using RNAseq alone. To circumvent this issue, we implemented the parsing of the list of missing genes and added them to the final prediction (which modestly increased the rate of false gene duplications as well: C: 98.9% [S: 96.2%,D: 2.7%], F: 0.5%, M: 0.6%). However, we recommend the user to carefully monitor the BUSCO score at each step and decide whether such genes should be added or not. In the same vein, we noted that the longest transcript among multiple alternative transcripts may not be a BUSCO gene, as this later sometimes corresponded to a transcript of shorter size. Therefore, we also implemented some minor corrections to retrieve those transcripts initially predicted but dropped during the size-selection step. Results obtained after reannotation of all 42 genomes from our lab are provided in **Table S5**.

**Table 2.** Evaluation of genome annotation results in the case *Microbotryum lychnidis-dioicae* 1064. BUSCO (v5.7.1) scores are provided along with a number of basic statistics for the genome of one mating type (*M. lychnidis-dioicae 1064 a*_*1*_). The results for the genome of the second mating type (*M. lychnidis-dioicae 1064 a*_*2*_) are provided in **Table S4**. Only the longest transcript was kept for each gene for the BUSCO evaluation. Results are presented for data either i) with only RNAseq data, ii) only the OrthoDB11 dataset for fungi, or iii) a custom database (CustomDB) built from previous gene annotation work in the lab. Column “all” refers to the combination of the three previous data; %uniprot match and %interproscan matches refer to the number of BRAKER genes with at least one match against either uniprot database or the whole databases used in InterProScan. Finally, the number of single-copy orthologs between the ancestral-like mating-type chromosome of *M. lagerheimii* and the *a*_*1*_/*a*_*2*_ mating-type chromosomes of *M. lychnidis-dioicae* are provided.

|  | RNAseq only | OrthoDB11 | CustomDB | All |
| --- | --- | --- | --- | --- |
| **Number of genes** | 9,850 | 9,778 | 9,363 | 10,568 |
| **Number of transcripts** | 11,945 | 13,629 | 12,670 | 14,091 |
| **Mean CDS length (bp)** | 1,626.15 | 1,633.53 | 1,704.16 | 1,527.57 |
| **Number of introns per gene** | 4.71 | 4.3 | 4.45 | 4.54 |
| **BUSCO** | C:98.9%<br>[S:98.3%,D:0.6%]<br>] F:0.2%,M:0.9% | C:98.6%<br>[S:98.0%,D:0.6%]<br>F:0.3%,M:1.1% | C:97.4%<br>[S:96.8%,D:0.6%]<br>F:0.3%,M:1.2% | C:98.3%<br>[S:97.4%,D:0.6%]<br>] F:0.4%,M:1.6% |
| **Complete BUSCOs (C)** | 1,744 | 1,739 | 1,723 | 1,734 |
| **Complete and single-copy BUSCOs (S)** | 1,734 | 1,728 | 1,712 | 1,718 |
| **Complete and duplicated BUSCOs (D)** | 10 | 11 | 11 | 10 |
| **Fragmented BUSCOs (F)** | 3 | 5 | 10 | 7 |
| **Missing BUSCOs (M)** | 17 | 20 | 31 | 23 |
| **Total BUSCO groups searched** | 1,764 | 1,764 | 1,764 | 1,764 |
| **Uniprot match (%)** | 62.49 | 64.49 | 54.84 | 59.84 |
| **InterproScan match (%)** | 90.18 | 92.39 | 93.57 | 89.13 |
| **Single-copy orthologs present on both mating-type chromosomes** | 620 | 631 | 623 | 634 |

### Genome and Gene Comparisons

Following the gene prediction steps, different operations will be launched if requested. Once the genes are correctly filtered, they are fed to GeneSpace, which internally runs Orthofinder, Blast and MCScanX to produce a rich output, suitable both for whole genome comparison and for focusing on the non-recombining region of interest (ex: **Fig S3, Fig 3A**). The output itself can be further processed by the user. Next, we used the output of the Phylogenetic Hierarchical Orthogroups (N0.tsv) to infer single-copy orthologs whose numbers are provided in the last row of Table 2; highly similar numbers were inferred with the various alternative approaches. These single-copy orthologs are further used to construct an ideogram to display microsynteny (gene-by-gene links between all single-copy orthologs) considering both mating-type chromosomes, or one mating-type chromosome and an ancestral-like genome (**Fig 5**); and to construct a circos plot based on the same data (**Fig 3B**) and highlight genes of interest, if any. The purpose of these ideogram and circos plots is two-fold: i) helping understanding the extent of rearrangements between the two mating-type or sex chromosomes, and relative to the ancestral sequence if such sequence is available, and ii) draw hypotheses regarding the number of evolutionary strata. Moreover, the code will generate both uncolored ideograms and circos plots as well as colored ideograms and circos plots based on the distribution of strata inferred from the MCP analysis (**Fig S4**) as well as ideograms and circos plots with colors based on the quantiles of observed dS values. As a complement to the gene synteny analysis, broad patterns of synteny along sex chromosomes are plotteed automatically using the pafr plot_synteny function following a minimap alignment of the genome between each other (**Fig S5**) and dotplots are generated on the fly (**Fig S6**). While this is not a central analysis for the identification of evolutionary strata, it is a straightforward approach to identify inversions, as well as possible assembly errors.

**Figure 3.**
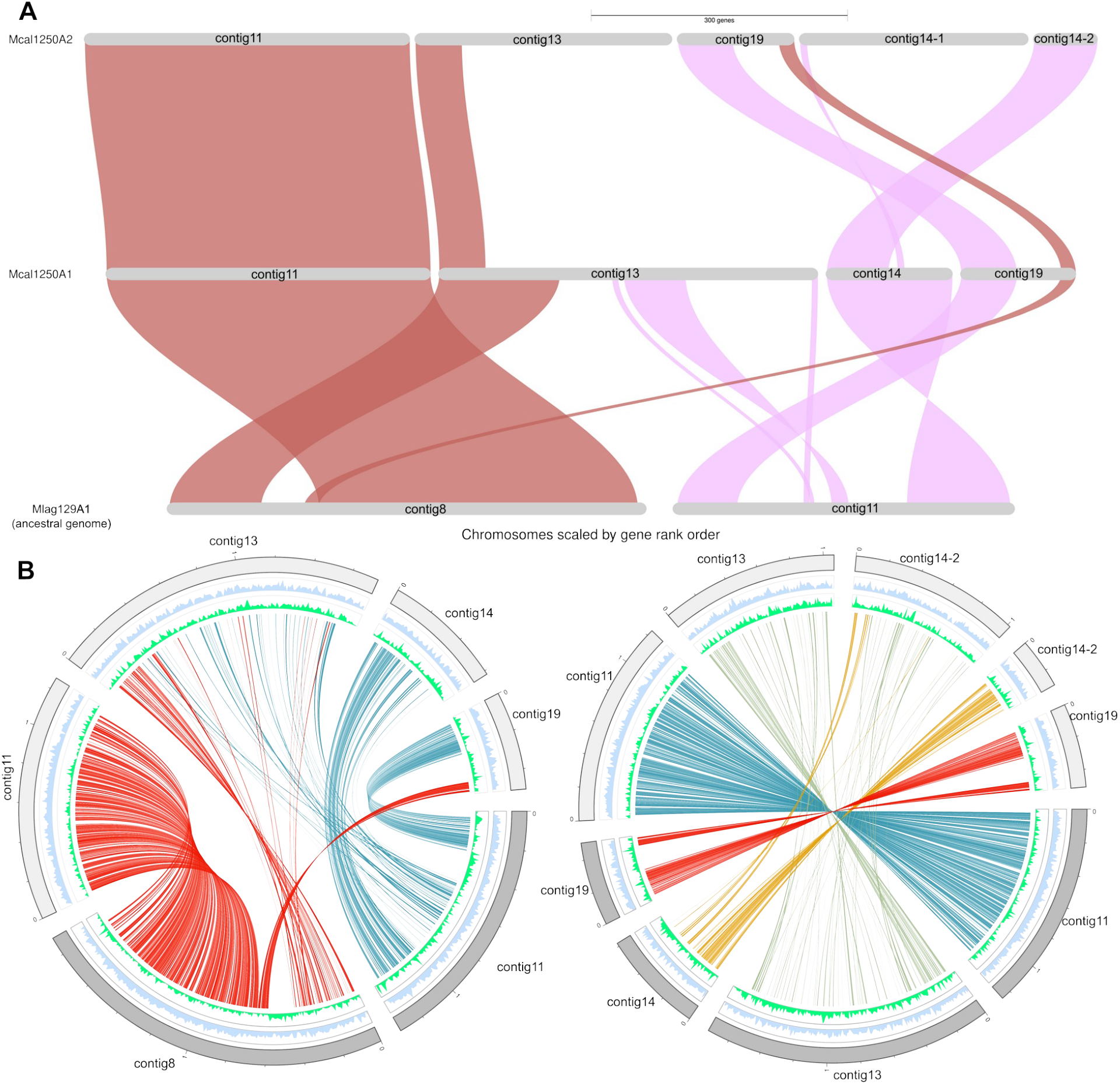
GeneSpace (A) and Circos plots (B) showing synteny and orthology relationships between mating-type chromosomes of *Microbotryum lagerheimii (Mlag129A1)* and *M. v. caroliniana 1250* a_1_ and between the two mating-type chromosomes of *M. v. caroliniana 1250* (a_1_ and a_2_) Contig 8 corresponds to the HD chromosome and contig11 to the PR chromosome. **A)** Plots showing major synteny blocks as inferred by combining orthofinder, blasts and MCScanX results. Synteny groups must include at least five consecutive genes. The large synteny observed between contig8 and the two corresponding contigs in *M. v. caroliniana* a_1_ and a_2_ indicates that a large part of the ancestral HD mating-type chromosome is now an autosome. **B)** Circos plots of the *M. v. caroliniana 1250 a*_*1*_ mating-type chromosome compared to the *M. lagerheimii* HD and PR mating-type chromosomes (lefte) and *M. v. caroliniana 1250 a*_*1*_ versus *a*_*2*_ mating-type chromosomes (right). The inner circle displays gene density in light blue and TE density in spring green respectively. Afteer d_S_ computation, additional circos plots are automatically constructed with inner links colored based on the quantiles of computed d_S_ values instead of being colored by contigs. Afteer the strata inference step, external links are also colored according to the inferred strata (see working examples on github).

### Inference of evolutionary strata

Next, orthofinder single-copy orthologs are reformatteed and corresponding CDS are passed as input to MACSE for alignment. The orthologs are used to compute d_S_ and d_N_ values, as shown in **Fig 4A**. For the changepoint analysis (**Fig S7**), the gene order is crucial to inference. When this information is not known *a priori*, users can run the pipeline ‘as is’ using default values, identify reversed contigs, and rerun the last plotteing steps (options 7 and 8 in the workflow), which typically takes a few minutes at most (**Fig 4B**).

**Figure 4.**
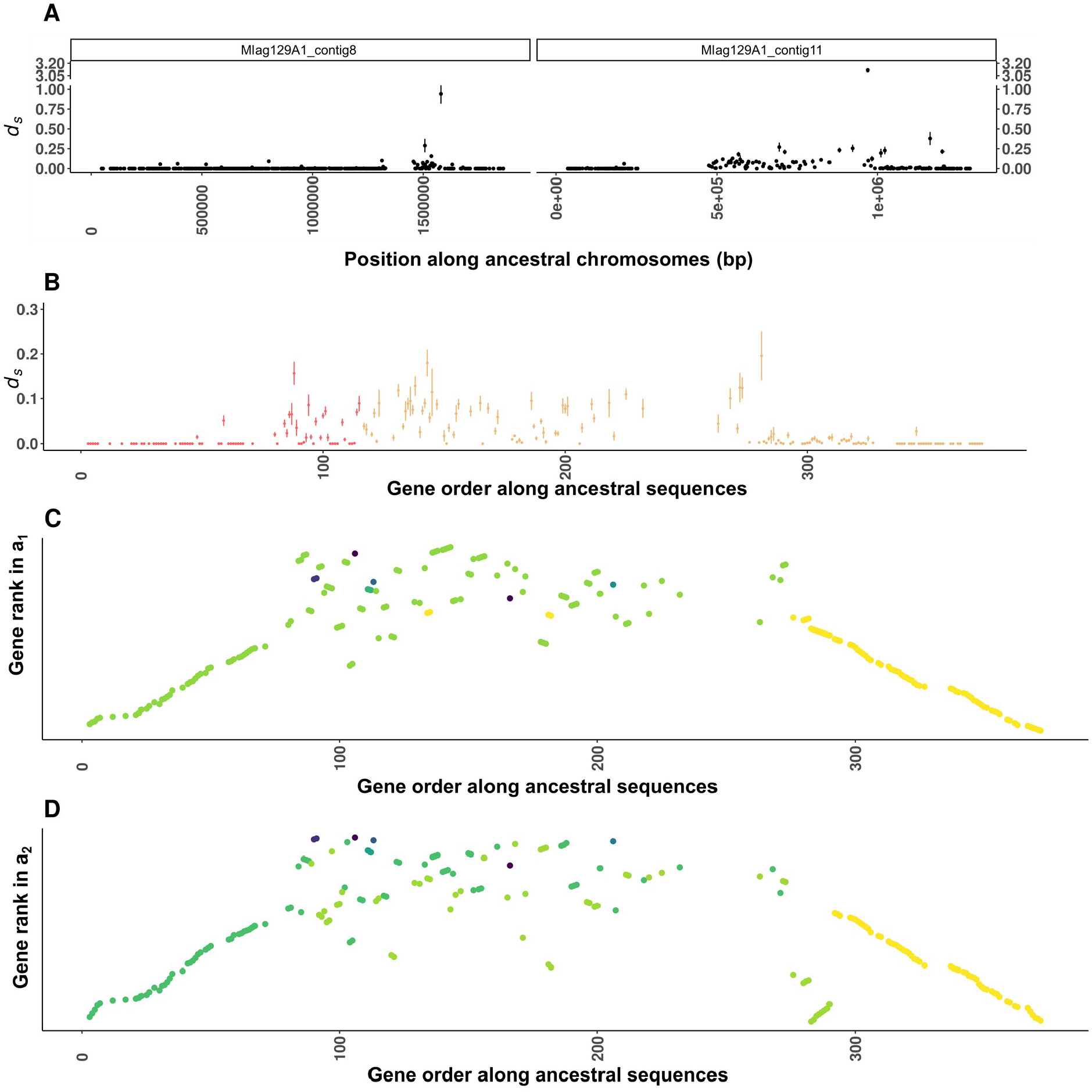
Observed d_S_ values and rearrangements in the *Microbotryum v. caroliniana* case. **A)** d_S_ values between the *M. v. caroliniana* a_1_ and a_2_ mating-type chromosomes along the mating-type chromosomes of *M. lagerheimii*, with the contig 11 corresponding to the PR chromosome and the contig 8 corresponding to the HD chromosome. Points represent the per-gene d_S_ values plotteed according to the genomic positions of the genes along the chromosome (in base pairs). B) Distribution of the d_S_ values along the ancestral gene order (i.e., the gene order on the *M. lagerheimii* mating-type chromosomes) instead of the genomic coordinates (in base pairs). Contigs are reversed to match the order of the chromosomal rearrangement and fusion in *M. v. caroliniana* (i.e., contig 8 (HD) in red and contig 11 (PR) in orange). The large region in contig 8 that became an autosome upstream of the centromere in panel A (>500 kbp) has been removed. The region downstream of the centromere in contig 11 (<460 kbp) is also removed. C) and D) Current rank of the genes in the a _1_ and a_2_ mating-type chromosomes along the ancestral gene order, showing the rearrangements compared to the ancestral state, points are colored according to the contig of origin in M. v. caroliniana a1 or a2 assemblies.

We applied the pipeline to the *M. v. caroliniana* dataset. Afteer a first pass we identified the contigs forming the mating-type chromosomes, and their homology with those of *M. lagerheimii* **(Fig 3)** and plotteed automatically the d_S_ values (**Fig 4A**). The inference of the rearrangements having formed the *M. v. caroliniana* mating-type chromosomes allowed to decide the order of genes to be plotteed on the X axis for the d_S_ plot (**Fig 4B**) and to perform the changepoint analysis. The model comparison from the changepoint analysis indicates that the best models was those with 7 and 5 changepoints based on the leave-one-out weight approach (weight = 0.577 and 0.339, **Table S7**), corresponding to 5 and 3 evolutionary strata along with the 2 PARs (pseudo-autosomal regions) on each side (**Table S8** for example of the MCP output). Analysis of the difflerent changepoint locations supported previous inferences of evolutionary strata (**Fig 5, Fig S7-S10, Table S9** for detailed Bayes-Factor). While the leave-one-out cross validation procedure provides insight into which models perform best, the shape of the posterior fitteed values (**Fig S7**) also need to be carefully inspected. Similarly, the posterior plot and mixing and convergence of the MCMC are central checks that need to be performed, as for any Bayesian analysis (**Fig S11**). While a first analysis is implemented with default setteings in the “automated” approach of our workflow (see e.g., **Fig S12** for MCP without priors), the R environment will be entirely exported and we strongly recommend the users to further explore the results manually and test if the evolutionary strata inferred fit with their working hypothesis and get support from other analyses, such as the identified rearrangements from the synteny analysis, or comparisons with closely related species that may display, for example, difflerent patteerns for young evolutionary strata. Here, for instance, priors were used regarding the location of the changepoints along the gene order based on the visual change in mean d _S_ values. The changepoint analysis supported the existence of the previously reported strata in [43]. There was no significant difflerence in the mean d_S_ between the youngest evolutionary stratum supporting previous work (light blue in [43]) and the PAR (p>0.5, **Fig S7-S9** including all detailed statistical tests); BayesFactor from the MCP analysis also returned only modest evidence for the di fflerence in d_S_ values (BF = 2.99, see table **S9** for all bayes factor).

We provide additional results and examples of evolutionary strata inference from the species *M. v. tatarinowii* (strain 1400) in **Figs S13 to S19** from the species *Microbotryum lychnidis-dioicae* (strain 1064) which was annotated with RNAseq above (**Figs S20 to S24**) and for the threespine stickleback (*Gasterosteus aculeatus*) genome assembly, using published gene predictions as well as a new set of gene predictions **(Figs S25 to S30)**.

**Figure 5.**
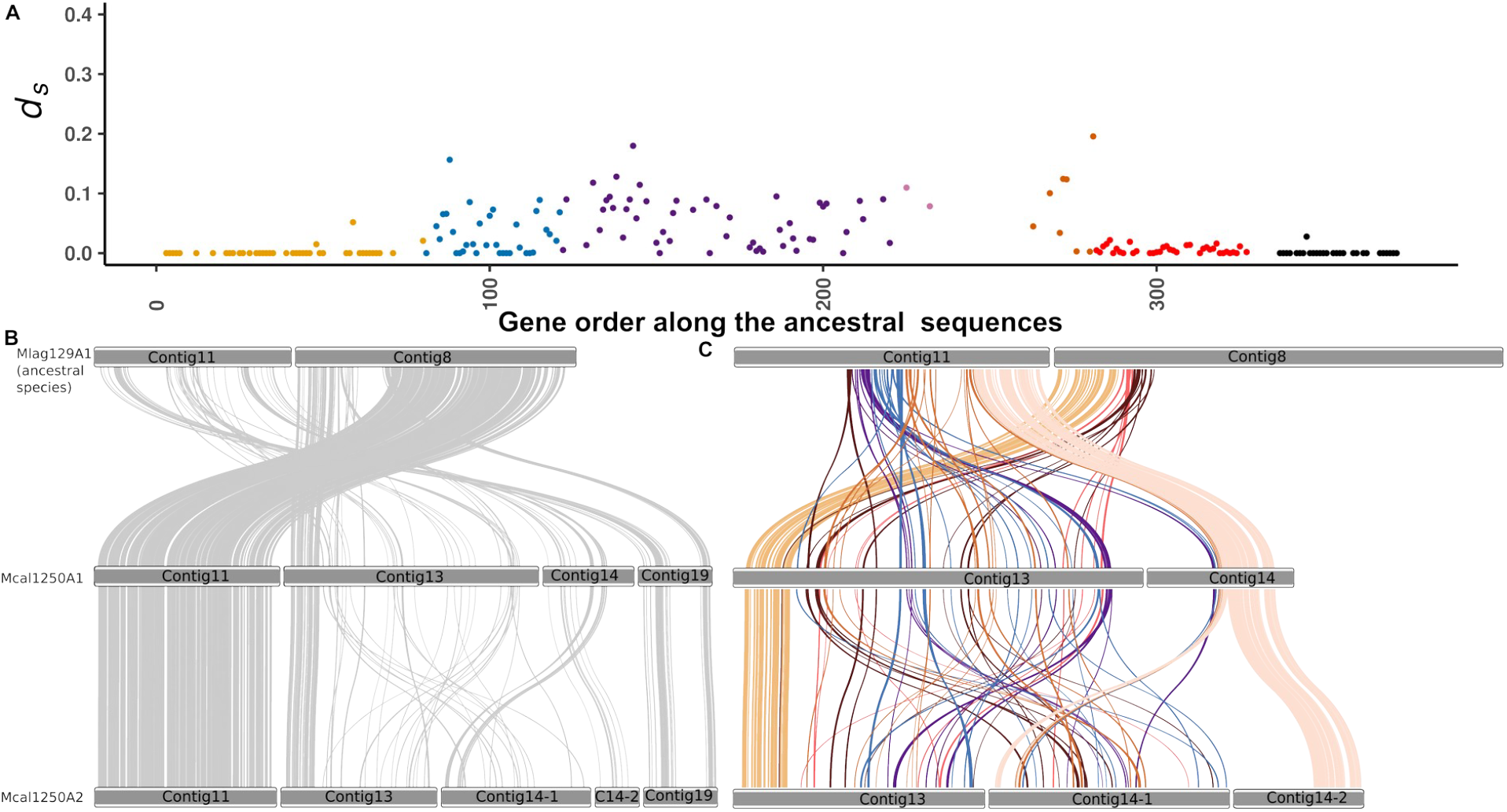
Evolution of mating-type chromosomes in *Microbotryum violaceum caroliniana*. A) Localisation of the evolutionary strata in *M. v. caroliniana* and their d_S_ values along the ancestral gene rank, using as proxy *M. lagerheimii (Mlag129a1)*. Each point represents the value for a gene (See **Fig S4** for the same plots with difflerent numbers of evolutionary strata). Panels B and C display ideograms plot showing links between single-copy orthologous genes of *M. v. caroliniana* a_1_ or a_2_ and the genome of *M. lagerheimii* used as a proxy for the ancestral gene order (*Mlag29a1*). **B)** The simplest version with grey links displays all links, including those on the current autosomal region in *M. v. caroliniana*. **C)** Single-copy orthologs (n = 201) used for the changepoint analysis colored according to their inferred evolutionary strata in the MCP analysis. As for the Circos plots, links are automatically colored according to quantiles of computed d_S_ values, which facilitate biological interpretation (see working examples on github).

## Discussion

The increasing number of available genome assemblies opens up new avenues for evolutionary research but can be at the same time a significant challenge, given the numerous softwares, dependencies and sometimes complex tools that might be needed to correctly annotate and analyse large-scale genomic data. In addition, the inference of evolutionary strata has often relied on simple visual inspection of d_S_ plots without any statistical tests or other lines of evidence, that can be presence/absence of strata in closely related species, rearrangements and transpecific polymorphisms [6]. Here, we provide a straightforward workflow that aims to simplify the various tasks required to infer evolutionary strata along non-recombining regions in genomes, for example in sex chromosomes. This workflow is flexible and should scale well with genome assemblies of any size. We provide examples of its application with the small genomes of *Microbotryum* fungi (30 to 45 Mb genomes with 1 to 10 Mb mating type chromosomes) and for the threespine stickleback (472 Mb genome with 20.5 and 15.8 Mb long X and Y chromosome, respectively), and it was also useful for identifying evolutionary strata in the giant Y chromosome of the plant *Silene latifolia* (2.9 Gb genome with a ∼ 350 and ∼500 Mb long X and Y chromosomes, respectively). It is also possible to use it to simply perform gene prediction, in which case only the TE discovery and Braker steps will be run. Similarly, if gene models and genome assemblies are already available, it is possible to skip the gene prediction step, so that only the assembly mapping, orthology, synteny and/or d _S_ analyses will be performed, providing a high level of flexibility to the end users. While we developed EASYstrata with the aim of comparing alternative haploid assemblies, it can also be seamlessly used to compare haploid-resolved assemblies of difflerent species, for example when exploring genome rearrangements and their correlation with sequence-level divergence. We have not systematically tested how divergent two haploid-resolved genomes can be for the approach to remain informative, but we have successfully used EASYstrata to compare species separated by several million years of evolution. This includes comparisons with an ancestral-state proxy in Microbotryum species, as well as identifying homologous blocks of sex chromosomes in the dioecious species *Silene latifolia* and the monoecious species *S. conica* and *S. vulgari*s

The quality of the analyses will depend on the availability of an accurate proxy for the ancestral gene order, especially for mating-type chromosomes and other supergenes for which both haplotypes are non-recombining, and therefore potentially rearranged. In the absence of a proxy for an ancestral arrangement, we recommend to use the order along the X or Z chromosome in the case of sex chromosomes, and an outgroup is still recommended to help identify single-copy orthologs [11]. Another critical factor is the quality of the genome assemblies. Users should aim at providing the most accurate and contiguous assembly, as assessed by the BUSCO completeness at the genome level, QV statistics, N50 and other standard measures. In particular, orientation errors and other kinds of misassemblies that mimic inversions may result in issues for the detection of evolutionary strata and false interpretations of degeneration signals associated with the loss of recombination. By default, the pipeline performs minimap alignments and constructs dot plots on the fly among the genomes to be studied, which provides an easy way to spot remaining assembly errors. While minimap is not designed to investigate full synteny, other minimizer-based approaches, such as nt-synt [44], could be used to complement the implemented analysis. Depending on the intended use, the contiguity of the genome assemblies is more or less important. If users are interested exclusively in obtaining gene and transposable elements, users can set the master script to option 6, even on a single genome, regardless of fragmentation status. For analysis including minimizer alignment and/or orthology based synteny between haplotypes (options 1 to 5) we have successfully treated cases in which one of the haploids is fragmented in up to 10 contigs [45]; in this case the most critical point is the ability to locate collinear regions in at least one end of the compared haplotigs, as this permits to orient the synteny plots. If a proxy for the ancestral rank is available, and thus evolutionary strata can be inferred (examples 1, 5-7) the fragmentation of haploid assemblies is not important as far as the genome annotation for both compared haploids is suffliciently complete. We issue a warning if BUSCO score drops below 85%.

Although our approach does not aim to identify sex chromosomes, it can indeed be used for this task in specific cases of anisogamic sex chromosomes, where autosomes are expected to be largely collinear and sex chromosomes to display re-arrangements that could be readily spot in the synteny plots produced by EASYstrata (see example 1 in the public repository). In such cases, we offler a mapping and kmer counting free approach complementing existing methods like SEX-DETector [46], DicoverY [47], or FIndzx [48].

Some bona fide pseudo-genes are included by default, although their precise identification should be performed afteerwards (by looking at predicted genes with nonsense mutations). In our tests with Microbotryum mating-type chromosomes, known strata could be readily retrieved with as few as ∼150 genes. Our pipeline crucially relies on the computation of the d_S_ values to detect evolutionary strata. We have never performed inferences with fewer than approximately one hundred and fiftey genes. Therefore, we recommend the users to carefully interpret their results when a smaller number of single-copy orthologs are available.

In addition, the accuracy of gene annotation may be affected by the availability of RNAseq data. This can be an issue if the studied group is poorly represented in existing databases, resulting in few external protein sequence datasets being available. At the end of the genome annotation steps, BUSCO scores are automatically computed. Users should aim at scores well above 90% of complete gene sets to confidently trust the rest of the analysis. In addition, some proteins might have a highly dynamic or particular evolution, especially considering the rate of degeneration of sex chromosomes, and may thus be hard to predict; this is particularly the case of the PR gene in the *Microbotryum* case, but is likely to be a more general issue. An easy approach to circumvent this problem, that we successfully implemented, is to transfer the annotation, using Miniprot [49], from a phylogenetically close species in which the protein was identified. We included an example script to do so in our workflow.

Our workflow should be appropriate for a wide number of taxa, provided that genome assembly quality is sufflicient. A limiting factor is the availability of external transcriptomic data, which can improve the quality of gene prediction in aiding the prediction of genuine genes. For instance, the gene annotation softeware used here (BRAKER) was successfully applied to annotate 200 insect genomes [50]. Yet, overly large genomes may be difflicult to annotate with BRAKER; for instance, we failed to accurately annotate the 10 gb *Bombina variegata* genome afteer several weeks of computation (BUSCO score ∼ 30%, same results being obtained with Helixer [51]). Yet, a simple alternative with Miniprot, using sets of closely related proteins, produced a relatively decent gene annotation (BUSCO score ∼ 85%). This approach can be a promising alternative, with the caveat that genuine genes will be missed if not present in the closely related species

Here, we provide a simple way to assess changes in mean d_S_ values along non-recombining regions in order to identify evolutionary strata. Our workflow uses functions implemented in the MCP package, which is one of the most comprehensive for this purpose to our knowledge. Its Bayesian basis allows an assessment of the credible intervals around mean changepoints, as well as an assessment of the support for difflerent changepoint sites and difflerent models. However, the results of the MCP analyses, especially the support for a given model, should be critically weighted by the user to make sure that the inferred strata make biological sense and that they are consistent with other types of evidence. Ideally, this analysis should be complemented by analyses of i) structural rearrangements (e.g., inversions, deletions, transpositions and duplications), as these likely affect recombination and accumulate with time in non-recombining regions [3], ii) the presence/absence of footprints of recombination suppression in closely related species, if available, as young evolutionary strata may be specific to some species [6], and iii) the segregation of alleles and trans-specific polymorphism to infer the age of recombination suppression relative to speciation events [6,52]. As these analyses heavily rely on the availability of outgroup data, they are not covered by our workflow, but can be conducted using phylogenetic approaches. Trans-specific polymorphism can be examined by inferring genealogical trees for the set of genes belonging to each inferred stratum. In case of evolutionary strata having evolved between different speciation events, the expectation is that gene trees in the difflerent strata display difflerent topologies and difflerent branch lengths.

Our workflow is rather modular and combines a set of state-of-the-art softewares. However, as bioinformatics is a fast-paced field, it is possible to modify our workflow to take advantage of newly developed tools. For instance, we obtained quick and high-quality gene prediction for *Microbotryum* fungi using the newly developed Helixer pipeline [51]. This could be easily combined with BRAKER to provide a high-quality set of genes [11]. The modularity of our workflow should allow an easy replacement of the implemented tools by other tools, especially regarding future versions of BRAKER or KAKScalculator instead of PAML. We expect that the flexibility of our workflow will facilitate the systematic discovery of evolutionary strata across the tree of life to help understand patterns and evolutionary processes.

## Data Availability

The code and data are available on GitHub: https://github.com/QueentinRougemont/EASYstrata/. All datasets used to illustrate the pipeline were previously published and have been deposited in a single repository (https://doi.org/10.5281/zenodo.14744965) to ease reproducibility. The *M. lagerheimii* 129.01 new genome assembly has been deposited at ENA under project PRJEB89 554.

## Acknowledgments

We would like to thank Jean-Philippe Vernadet for help with the server maintenance as well as Michel Bartoli for collection of the *M. lagerheimii* 129.01 strain.

## Author Contributions

Conceptualization: RCRdlV, QR, TG; Data curation: QR, RCRdlV; Formal analysis: QR; Funding acquisition: TG; Investigation: QR; Methodology: QR, RCRdlV; Project administration: TG; Resources: EL, LB, AS; Softeware: QR, EL, LB; Supervision: TG, RCRdlV; Validation: EL, LB, AJd, RCRdlV; Visualization: QR; Writing – original dra fte: QR; Writing – review & editing: all authors; Final drafte was prepared by QR, RCRdlV and TG.

## References

[1] Formenti G, Theissinger K, Fernandes C, Bista I, Bombarely A, Bleidorn C, et al. The era of reference genomes in conservation genomics. Trends in Ecology & Evolution 2022;37:197–202. 10.1016/j.tree.2021.11.008.

[2] Yue J, Krasovec M, Kazama Y, Zhang X, Xie W, Zhang S, et al. The origin and evolution of sex chromosomes, revealed by sequencing of the Silene latifolia female genome. Current Biology 2023;33:2504-2514.e3. 10.1016/j.cub.2023.05.046.

[3] Jay P, Jefflries D, Hartmann FE, Véber A, Giraud T. Why do sex chromosomes progressively lose recombination? Trends in Genetics 2024;40:564–79. 10.1016/j.tig.2024.03.005.

[4] Charlesworth D. The status of supergenes in the 21st century: recombination suppression in Batesian mimicry and sex chromosomes and other complex adaptations. Evolutionary Applications 2016;9:74–90. 10.1111/eva.12291.

[5] Lahn BT, Page DC. Four Evolutionary Strata on the Human X Chromosome. Science 1999;286:964–7. 10.1126/science.286.5441.964.

[6] Branco S, Badouin H, Rodríguez de la Vega RC, Gouzy J, Carpentier F, Aguileta G, et al. Evolutionary strata on young mating-type chromosomes despite the lack of sexual antagonism. Proceedings of the National Academy of Sciences 2017;114:7067–72. 10.1073/pnas.1701658114.

[7] Hartmann FE, Duhamel M, Carpentier F, Hood ME, Foulongne-Oriol M, Silar P, et al. Recombination suppression and evolutionary strata around mating-type loci in fungi: documenting patteerns and understanding evolutionary and mechanistic causes. New Phytologist 2021;229:2470–91. 10.1111/nph.17039.

[8] The Darwin Tree of Life Project Consortium. Sequence locally, think globally: The Darwin Tree of Life Project. Proceedings of the National Academy of Sciences 2022;119:e2.115 642 118. 10.1073/pnas.2115642118.

[9] Weisman CM, Murray AW, Eddy SR. Mixing genome annotation methods in a comparative analysis inflates the apparent number of lineage-specific genes. Curr Biol 2022;32:2632-2639.e2. 10.1016/j.cub.2022.04.085.

[10] Álvarez-Carretero S, Kapli P, Yang Z. Beginner’s Guide on the Use of PAML to Detect Positive Selection. Molecular Biology and Evolution 2023;40:msad041. 10.1093/molbev/msad041.

[11] Moraga C, Branco C, Rougemont Q, Jedlička P, Mendoza-Galindo E, Veltsos P, et al. The Silene latifolia genome and its giant Y chromosome. Science 2025;387:630–6. 10.1126/science.adj7430.

[12] Duhamel M, Hood ME, Rodríguez de la Vega RC, Giraud T. Dynamics of transposable element accumulation in the non-recombining regions of mating-type chromosomes in anther-smut fungi. Nat Commun 2023;14:5692. 10.1038/s41467-023-41413-4.

[13] Peichel CL, McCann SR, Ross JA, Naftealy AFS, Urton JR, Cech JN, et al. Assembly of the threespine stickleback Y chromosome reveals convergent signatures of sex chromosome evolution. Genome Biology 2020;21:177. 10.1186/s13059-020-02097-x.

[14] Mufflato M, Louis A, Nguyen NTT, Lucas J, Berthelot C, Roest Crollius H. Reconstruction of hundreds of reference ancestral genomes across the eukaryotic kingdom. Nat Ecol Evol 2023;7:355–66. 10.1038/s41559-022-01956-z.

[15] Gabriel L, Brůna T, Hoffl KJ, Ebel M, Lomsadze A, Borodovsky M, et al. BRAKER3: Fully automated genome annotation using RNA-seq and protein evidence with GeneMark-ETP, AUGUSTUS and TSEBRA. bioRxiv 2024:2023.06.10.544449. 10.1101/2023.06.10.544449.

[16] Tegenfeldt F, Kuznetsov D, Manni M, Berkeley M, Zdobnov EM, Kriventseva EV. OrthoDB and BUSCO update: annotation of orthologs with wider sampling of genomes. Nucleic Acids Research 2024:gkae987. 10.1093/nar/gkae987.

[17] Simão FA, Waterhouse RM, Ioannidis P, Kriventseva EV, Zdobnov EM. BUSCO: assessing genome assembly and annotation completeness with single-copy orthologs. Bioinformatics 2015;31:3210–2. 10.1093/bioinformatics/btv351.

[18] Flynn JM, Hubley R, Goubert C, Rosen J, Clark AG, Feschottee C, et al. RepeatModeler2 for automated genomic discovery of transposable element families. Proc Natl Acad Sci U S A 2020;117:9451–7. 10.1073/pnas.1921046117.

[19] Smit Hubley, R, Green, P. RepeatMasker Open-4.0. 2013.

[20] Brůna T, Lomsadze A, Borodovsky M. GeneMark-ETP significantly improves the accuracy of automatic annotation of large eukaryotic genomes. Genome Res 2024;34:757–68. 10.1101/gr.278373.123.

[21] Stanke M, Keller O, Gunduz I, Hayes A, Waack S, Morgenstern B. AUGUSTUS: ab initio prediction of alternative transcripts. Nucleic Acids Res 2006;34:W435–439. 10.1093/nar/gkl200.

[22] Bolger AM, Lohse M, Usadel B. Trimmomatic: a flexible trimmer for Illumina sequence data. Bioinformatics 2014;30:2114–20. 10.1093/bioinformatics/btu170.

[23] Wu TD, Reeder J, Lawrence M, Becker G, Brauer MJ. GMAP and GSNAP for Genomic Sequence Alignment: Enhancements to Speed, Accuracy, and Functionality. Methods Mol Biol 2016;1418:283–334. 10.1007/978-1-4939-3578-9_15.

[24] Li H, Handsaker B, Wysoker A, Fennell T, Ruan J, Homer N, et al. The Sequence Alignment/Map format and SAMtools. Bioinformatics 2009;25:2078–9. 10.1093/bioinformatics/btp352.

[25] Gabriel L, Hoffl KJ, Brůna T, Borodovsky M, Stanke M. TSEBRA: transcript selector for BRAKER. BMC Bioinformatics 2021;22:566. 10.1186/s12859-021-04482-0.

[26] Pertea G, Pertea M. GFF Utilities: GfflRead and GfflCompare 2020. 10.12688/f1000research.23297.1.

[27] Li H. New strategies to improve minimap2 alignment accuracy. Bioinformatics 2021;37:4572–4. 10.1093/bioinformatics/btab705.

[28] Winter Dd. pafr: Read, Manipulate and Visualize “Pairwise mApping Format” Data. R package 2025.

[29] Emms DM, Kelly S. OrthoFinder: phylogenetic orthology inference for comparative genomics. Genome Biology 2019;20:238. 10.1186/s13059-019-1832-y.

[30] Lovell JT, Sreedasyam A, Schranz ME, Wilson M, Carlson JW, Harkess A, et al. GENESPACE tracks regions of interest and gene copy number variation across multiple genomes. eLife 2022;11:e78.526. 10.7554/eLife.78526.

[31] Buchfink B, Reuter K, Drost H-G. Sensitive protein alignments at tree-of-life scale using DIAMOND. Nat Methods 2021;18:366–8. 10.1038/s41592-021-01101-x.

[32] Wang Y, Tang H, Debarry JD, Tan X, Li J, Wang X, et al. MCScanX: a toolkit for detection and evolutionary analysis of gene synteny and collinearity. Nucleic Acids Res 2012;40:e49. 10.1093/nar/gkr1293.

[33] Hao Z, Lv D, Ge Y, Shi J, Weijers D, Yu G, et al. RIdeogram: drawing SVG graphics to visualize and map genome-wide data on the idiograms. PeerJ Comput Sci 2020;6:e251.10.7717/peerj-cs.251.

[34] Gu Z, Gu L, Eils R, Schlesner M, Brors B. circlize Implements and enhances circular visualization in R. Bioinformatics 2014;30:2811–2. 10.1093/bioinformatics/btu393.

[35] Yang Z. PAML 4: Phylogenetic Analysis by Maximum Likelihood. Molecular Biology and Evolution 2007;24:1586–91. 10.1093/molbev/msm088.

[36] Ranwez V, Douzery EJP, Cambon C, Chantret N, Delsuc F. MACSE v2: Toolkit for the Alignment of Coding Sequences Accounting for Frameshiftes and Stop Codons. Molecular Biology and Evolution 2018;35:2582–4. 10.1093/molbev/msy159.

[37] Edgar RC. MUSCLE: multiple sequence alignment with high accuracy and high throughput. Nucleic Acids Res 2004;32:1792–7. 10.1093/nar/gkh340.

[38] Abascal F, Zardoya R, Telford MJ. TranslatorX: multiple alignment of nucleotide sequences guided by amino acid translations. Nucleic Acids Research 2010;38:W7–13. 10.1093/nar/gkq291.

[39] Lindeløv J. mcp: An R Package for Regression With Multiple Change Points 2020. 10.31219/osf.io/fzqxv.

[40] Vehtari A, Gelman A, Gabry J. Practical Bayesian model evaluation using leave-one-out cross-validation and WAIC. Stat Comput 2017;27:1413–32. 10.1007/s11222-016-9696-4.

[41] Patil I. Visualizations with statistical details: The “ggstatsplot” approach. Journal of Open Source Softeware 2021;6:3167. 10.21105/joss.03167.

[42] Devier B, Aguileta G, Hood ME, Giraud T. Ancient trans-specific polymorphism at pheromone receptor genes in basidiomycetes. Genetics 2009;181:209–23. 10.1534/genetics.108.093708.

[43] Branco S, Carpentier F, Rodríguez de la Vega RC, Badouin H, Snirc A, Le Prieur S, et al. Multiple convergent supergene evolution events in mating-type chromosomes. Nat Commun 2018;9:2000. 10.1038/s41467-018-04380-9.

[44] Coombe L, Kazemi P, Wong J, Birol I, Warren RL. Multi-genome synteny detection using minimizer graph mappings 2024:2024.02.07.579356. 10.1101/2024.02.07.579356.

[45] Lucottee EA, Jay P, Rougemont Q, Boyer L, Cornille A, Snirc A, et al. Repeated loss of function at HD mating-type genes and of recombination suppression without mating-type locus linkage in anther-smut fungi 2024:2024.03.03.583 181. 10.1101/2024.03.03.583181.

[46] Muyle A, Käfer J, Zemp N, Mousset S, Picard F, Marais GA. SEX-DETector: A Probabilistic Approach to Study Sex Chromosomes in Non-Model Organisms. Genome Biol Evol 2016;8:2530–43. 10.1093/gbe/evw172.

[47] Rangavitteal S, Stopa N, Tomaszkiewicz M, Sahlin K, Makova KD, Medvedev P. DiscoverY: a classifier for identifying Y chromosome sequences in male assemblies. BMC Genomics 2019;20:641. 10.1186/s12864-019-5996-3.

[48] Sigeman H, Sinclair B, Hansson B. Findzx: an automated pipeline for detecting and visualising sex chromosomes using whole-genome sequencing data. BMC Genomics 2022;23:328. 10.1186/s12864-022-08432-9.

[49] Li H. Protein-to-genome alignment with miniprot. Bioinformatics 2023;39:btad014. 10.1093/bioinformatics/btad014.

[50] Saenko S, Hoffl KJ, Stanke M. Annotation of 200 Insect Genomes with BRAKER for Consistent Comparisons across Species 2025:2025.04.17.649312. 10.1101/2025.04.17.649312.

[51] Holst F, Bolger A, Günther C, Maß J, Triesch S, Kindel F, et al. Helixer–de novo Prediction of Primary Eukaryotic Gene Models Combining Deep Learning and a Hidden Markov Model 2023:2023.02.06.527280. 10.1101/2023.02.06.527280.

[52] Duhamel M, Carpentier F, Begerow D, Hood ME, Rodríguez de la Vega RC, Giraud T. Onset and stepwise extensions of recombination suppression are common in mating-type chromosomes of Microbotryum anther-smut fungi. Journal of Evolutionary Biology 2022;35:1619–34. 10.1111/jeb.13991.

