## supplementary files for "EASYstrata: An All-in-One Workflow for Genome Annotation and Genomic Divergence Analysis"

### Supplementary Methods

#### Genome sequencing of *Microbotryum lagerheimii 1290a1*

We sequenced the genome of the strain *M. lagerheimii* 129.01 $a_1$ using the Oxford Nanopore Technology with an R9 flow cell on a MinION device and performed genome polishing by sequencing short reads on an illumina HiSeq2500 platform providing 2x150 bp long paired-end reads. We obtained 214,992 ONT reads and 9.7 millions paired-end Illumina reads. The strain 129.01 was collected on *Silene vulgaris* in the French Pyrenees Mountains in 2002, in Sazos (GPS coordinates 42°52'51.84"N and 0°1'16.18"W). Sporidia were spreaded and cultivated on potato dextrose agar (PDA) media (Sigma-Aldrich, Saint-Quentin-Fallavier, France) for 3 days at 25°C. We harvested 300 - 400 mg of fresh material in a 2 mL Eppendorf tube and stored it overnight at −80°C before a lyophilization for 24h. DNA was extracted with the NucleoBond High Molecular Weight DNA kit from Macherey-Nagel (Hoerdt, France), with a mechanical disruption of about 30 mg of lyophilized mycelium with 2 tungsten beads (diameter 3 mm) for 5 min at 30 Hz (TissueLyser II, Qiagen, Courtaboeuf, France). We resuspended and transferred the grinding material in a 50 mL centrifuge tube by adding 900 µl of lysis buffer H1 and 200 µl of proteinase K. We added 4mL of lysis buffer H1 and increased the lysis time incubation up to 3 hours. We followed the manufacturer's instructions and eluted the gDNA in 100 µl of buffer HE (5 mM Tris-Cl,PH8.5). We processed a size selection on the gDNA by removing fragments below 10 kb with the Short Read Eliminator kit (PacBio, Handbook v2.0, Menlo Park, USA). The selected gDNA was eluted in 65 µl in the elution buffer. Quality and quantity controls were checked with Qubit 2.0 fluorometer (Lifetechnologies SAS, Courtaboeuf, France) and Nanodrop™ ND-2000 (Thermofisher, Les Ulis, France). The Nanopore library was prepared using the native barcoding of amplicons from the EXP-NBD114 kit associated with the SQK-LSK109 reagents (protocol version NBA_9093_V109_revD_12nov2019).

### Genome Assembly of *Microbotryum lagerheimii 1290a1*

The workflow provided in Rougemont 2024 was used to assemble the genome. Briefly, following long read sequencing, the ONT reads werereads trimmed with Chopper (De Coster and Rademakers, 2023) requiring a minimum quality of 8 and minimal length of 900 bp. The reads were assembled with flye v2.9.1(Kolmogorov *et al.*, 2019) with the option –genome-size=30m and –nano-raw. The resulting assembly was polished using the long reads with MarginPolish with default settings except for the config where minChunkSize was set to 5000 and maxDepth to 128. To this end, long reads were aligned to the draft assembly using Minimap2 (Li, 2021). The assembly was finally polished once with Pilon v1.24 (Walker *et al.*, 2014) (multiple runs of Pilon did not improve the QV or BUSCO scores). To do so we trimmed the reads with fastp (Chen, 2023) to require a minimal length of 100 and quality of 10, and aligned the cleaned reads with BWA mem2 (Vasimuddin *et al.*, 2019). The completeness of the genome assembly was assessed with BUSCO v5.5.1 after the flye step and following each polishing step using the basidiolicota_odb10 database of orthologous groups (n = 1764). The quality was further assessed using quast v5.1 and mercury to assess QV score and k-mer completeness and produce graphs of k-mer distribution. Scripts to reproduce the genome assembly are provided in Rougemont, (2024).

**Table S1: List of all third-party software**

| non conda dependencies | purpose | source |
|---|---|---|
| *BRAKER* | genome annotation | https://github.com/Gaius-Augustus/BRAKER |
| *protHint* | genome annotation | https://github.com/gatech-genemark/ProtHint/releases/download/v2.6.0/ProtHint-2.6.0.tar.gz |
| *Diamond* | Faster blast | https://github.com/bbuchfink/diamond/releases/download/v2.1.1/diamond-linux64.tar.gz |
| *cdbfasta* | genome annotation | https://github.com/gpertea/cdbfasta.git |
| *bamtools* | genome annotation | https://github.com/pezmaster31/bamtools |
| *htslib* | generic tools | https://github.com/samtools/htslib/releases/download/1.18/htslib-1.18.tar.bz2 |
| *Augustus* | genome annotation | https://github.com/Gaius-Augustus/Augustus.git |
| *Genemark* | genome annotation | https://github.com/gatech-genemark/GeneMark-ETP/ |
| *TSEBRA* | genome annotation | https://github.com/Gaius-Augustus/TSEBRA |
| *Gmap/Gsnap* | RNAseq mamping | http://research-pub.gene.com/gmap/src/gmap-gsnap-2023-10-10.v2.tar.gz |
| *gffread* | CDS/protein extraction | https://github.com/gpertea/gffread |
| *orthofinder* | orthology inference | https://github.com/davidemms/OrthoFinder/releases/download/2.5.5/OrthoFinder.tar.gz |
| *MCScanX* | synteny inference | https://github.com/wyp1125/MCScanX |
| *Macse* | protein alignment | https://www.agap-ge2pop.org/wp-content/uploads/macse/releases/macse_v2.07.jar |
| *paml(yn00/codeml)* | dn/ds analysis | https://github.com/abacus-gene/paml/releases/download/4.10.7/paml-4.10.7-linux-X86_64.tgz |
| *translatorx_vLocal.pl* | protein translation | http://161.111.160.230/cgi-bin/translatorx_vLocal.pl |
| *GeneSpace* | R package for synteny analysis | https://github.com/jtlovell/GENESPACE |

| Conda Dependencies | purpose | Conda Dependencies | purpose |
|---|---|---|---|
| python=3.12.8 | generic tools | perl-yaml-xs | braker dependencies |
| wget=1.21.4 | web download | matplotlib | python generic plotting tools |
| busco=5.7.1 | genome quality evaluation | cdbtools=0.99 | braker dependencies |
| repeatmasker=4.1.5 | TE masking | samtools=1.21 | indexing |
| repeatmodeler=2.0.5 | TE prediction | minimap2=2.28 | whole genome alignment |
| Curl=8.9.0 | web download | emboss=6.6.0 | various genomic tools |
| numpy=1.26.4 | generic tools | json5 | extension to json format |
| biopython=1.83 | generic python tools | jags=4.3.2 | Just Another Gibbs Sampler |
| make=4.3 | compiler | r-rjags=4.14 | Just Another Gibbs Sampler |
| cmake=3.28.3 | compiler | r-rsvg=2.6.0 | R package for svg viewing |
| pandas | generic formating python tools | r-igraph=2.0.2 | network analysis and visualisation |
| perl-app-cpanminus | braker dependencies | bioconductor-biostrings=2.70.1 | manipulation of biological strings |
| perl-hash-merge | braker dependencies | r-devtools=2.4.5 | tools to develop packages |
| perl-parallel-forkmanager | braker dependencies | r-curl=5.2.1 | R package (curl web interface) |
| perl-scalar-util-numeric | braker dependencies | r-tidyverse=2.0.0 | Data manipulation |
| perl-yaml | braker dependencies | r-cowplot=1.1.3 | R package (organising plot) |
| perl-class-data-inheritable | braker dependencies | r-viridis=0.6.5 | R package for colors |
| perl-exception-class | braker dependencies | r-ggrepel=0.9.6 | R package |
| perl-test-pod | braker dependencies | r-patchwork=1.3.0 | R package (organising plot) |
| perl-file-which | braker dependencies | r-wesanderson=0.3.7 | R package for colors |
| perl-mce | braker dependencies | r-ggbreak=0.1.2 | R package Set Axis Break for 'ggplot2' |
| perl-threaded | braker dependencies | r-circlize=0.4.16 | R package for circos plot |
| perl-list-util | braker dependencies | r-optparse=1.7.5 | R package to make options |
| perl-math-utils | braker dependencies | r-ggstatsplot=0.13.0 | R package for plots and stats |
| perl-list-moreutils | braker dependencies | r-pafr=0.0.2 | R package to view alignments |
| perl-file-homedir | braker dependencies | r-ggpubr=0.6.0 | ggplot2' Based Publication Ready Plots |
| perl-devel-size | braker dependencies | r-knitr=1.49 | R package to create nice report |
| perl-posix | braker dependencies | r-ggforce=0.4.2 | R package boosting ggplot2 |
| perl-file-spec | braker dependencies | r-rideogram=0.2.2 | R package for synteny analysis |
| perl-module-load-conditional | braker dependencies | r-mcp=0.3.4 | R package for changepoint analysis |
| perl-data-dumper | braker dependencies | | |

Table S2: characteristics of the new reference for *Microbotryum lagerheimii from* the strain *129.01.a1*

| Characteristics | 129.01.A1 |
|---|---|
| **Ncontigs** | 17 |
| **Length (bp)** | 25,762,778 |
| **Largest contigs (bp)** | 3,244,683 |
| **N50 (bp)** | 1,753,139 |
| **N90 (bp)** | 1,380,884 |
| **L50** | 6 |
| **L90** | 11 |
| **GC%** | 56.05 |
| **QV** | 42.45 |
| **Error rate** | 5.6e-05 |
| **Mean depth** | 116 |
| **K-mer completeness (%)** | 99.73 |
| **BUSCO from genome assembly** | S:98.9% (1745), D:0.23%(4),F:0.17%(3),I(0), M: 0.68% (15) |
| **BUSCO** ***from gene annotations*** | C:97.6%[S:97.0%,D:0.6%],F:0.8%,M:1.6%,n:1764 |

Table S3: Assembly characteristics of the genomes of the three *Microbotryum* species analysed for evolutionary strata as case examples with two haploid genomes per species, from the same strain for each species and of alternative mating types (a1 and a2).

| | *M. lychnidis-dioicae* 1064 $a_1$ | *M. lychnidis-dioicae* 1064 $a_2$ | *M. v. caroliniana* 1250 $a_1$ | *M. v. caroliniana* 1250 $a_2$ | *M. v. tatarinowii* 1400 $a_1$ | *M. v. tatarinowii* 1400 $a_2$ |
|---|---|---|---|---|---|---|
| **Ncontigs** | 48 | 37 | 131 | 137 | 24 | 26 |
| **Length** | 29 901 156 | 30 318 316 | 28 837 994 | 28 906 851 | 24 278 522 | 24 145 075 |
| **Largest contigs** | 3 412 169 | 4 051 571 | 2 207 875 | 3 481 606 | 3 362 224 | 3 353 295 |
| **N50** | 1 736 850 | 1 730 088 | 1 517 213 | 1 311 234 | 1 639 285 | 1 470 213 |
| **N90** | 648 002 | 1 040 457 | 503 608 | 296 686 | 1 048 688 | 917 274 |
| **L50** | 6 | 6 | 8 | 8 | 6 | 6 |
| **L90** | 16 | 14 | 20 | 21 | 13 | 14 |
| **GC%** | 55.6 | 55.6 | 56.11 | 56.04 | 55.97 | 55.94 |
| **BUSCO** | C:98.4% [S:97.7%,D:0.7%],F:0.2%,M:1.4% | C:98.4% [S:97.4%,D:1.0%],F:0.2%,M:1.4% | C:98.5% [S:98.0%,D:0.5%],F:0.3%,M:1.2% | C:98.5% [S:98.2%,D:0.3%],F:0.3%,M:1.2% | C:98.0% [S:97.7%,D:0.3%],F:0.3%,M:1.7% | C:98.0% [S:97.4%,D:0.6%],F:0.4%,M:1.6% |
| ***Complete BUSCOs (C )*** | 1736 | 1736 | 1737 | 1738 | 1730 | 1730 |
| ***Complete and single-copy BUSCOs (S)*** | 1724 | 1724 | 1229 | 1732 | 1724 | 1719 |
| ***Complete and duplicated BUSCOs (D)*** | 12 | 18 | 8 | 6 | 6 | 11 |
| ***Fragmented BUSCOs (F)*** | 3 | 8 | 6 | 6 | 6 | 7 |
| ***Missing BUSCOs (M)*** | 25 | 23 | 21 | 20 | 28 | 27 |
| ***Total BUSCO groups searched*** | 1764 | 1764 | 1764 | 1764 | 1764 | 1764 |

**Table S4: Full details of gene annotation for the different kinds of data used for training, on the *Microbotryum lychnidis-dioicae* $a_2$ genome.**

| | RNAseq only | OrthoDB11 | CustomDB | ALL |
|---|---|---|---|---|
| *Number of genes* | 9997 | 9998 | 9562 | 10 767 |
| *Nb transcripts* | 12 061 | 11 690 | 12 499 | 14 378 |
| *Mean CDS length* | 1602 | 1618.19 | 1689.97 | 1516.66 |
| *Number of introns* | 4.65 | 4.28 | 4.42 | 4.50 |
| *Complete genes* | 99.89 | 99.77 | 99.85 | NA |
| *BUSCO* | C:98.4% [S:97.5%,D:0.9%] F:0.6%,M:1% | C:98.4% [S:97.4%,D:1.0%] F:0.3%,M:1.3% | C:97.6% [S:96.7%,D:0.9%] F:0.6%,M:1.8% | C:98.1% [S:97.1%,D:1.0%] F:0.4%,M:1.5% |
| *Complete BUSCOs (C )* | 1735 | 1736 | 1720 | 1729 |
| *Complete and single-copy BUSCOs (S)* | 1720 | 1719 | 1705 | 1712 |
| *Complete and duplicated BUSCOs (D)* | 15 | 17 | 15 | 17 |
| *Fragmented BUSCOs (F)* | 11 | 6 | 10 | 7 |
| *Missing BUSCOs (M)* | 18 | 22 | 34 | 28 |
| *Total BUSCO groups searched* | 1764 | 1764 | 1764 | 1764 |
| *Prop. interproScan annotation* | 90.93 | 91.8 | 93.3 | 89.1 |
| *Uniprot match (%)* | 62.03 | 63.36 | 60.93 | 58.88 |

**Table S5: BUSCOs scores for all *Microbotryum* species:** scores for all 42 published *Microbotryum* genomes, reannotated with the present workflow. The conventional BUSCO score annotation is used with complete BUSCOs (C), complete and single-copy BUSCOs (S), complete and duplicated BUSCOs (D), fragmented BUSCOs (F) and missing BUSCOs (M). The number of genes was n = 1764 from the basidiomycota_odb10 dataset and scores were computed with BUSCO 5.5.1 and hmmsearch 3.1. See Table S4 for *M. lychnidis-dioicae* 1064 $a_1$ and $a_2$, respectively.

| species_id | accession n° | Busco_score |
|---|---|---|
| Mv-cal-1250-A1 | GCA_900014965.1 | C:98.0%[S:97.6%,D:0.4%],F:0.3%,M:1.7% |
| Mv-cal-1250-A2 | GCA_900014955.1 | C:97.9%[S:97.6%,D:0.3%],F:0.5%,M:1.6% |
| Mv-cor-1247-A1 | GCA_900015435.1 | C:97.2%[S:96.3%,D:0.9%],F:0.6%,M:2.2% |
| Mv-cor-1247-A2 | GCA_900009495.1 | C:94.9%[S:94.3%,D:0.6%],F:1.4%,M:3.7% |
| Mv-dio-1303-A1 | GCA_900120095.1 | C:94.9%[S:93.7%,D:1.2%],F:0.5%,M:4.6% |
| Mv-dio-1303-A2 | GCA_014805725.1 | C:94.8%[S:93.7%,D:1.1%],F:0.5%,M:4.7% |
| Mv-gra-01299-A1 | GCA_022506005.1 | C:97.9%[S:97.6%,D:0.3%],F:0.6%,M:1.5% |
| Mv-gra-01299-A2 | GCA_022505985.1 | C:97.3%[S:96.8%,D:0.5%],F:0.7%,M:2.0% |
| Mv-int-01389-A1 | GCA_022702645.1 | C:97.6%[S:97.1%,D:0.5%],F:0.2%,M:2.2% |
| Mv-int-1389-T | GCA_900096595.1 | C:97.5%[S:97.2%,D:0.3%],F:0.4%,M:2.1% |
| Mv-lag-1253-A1 | GCA_900015505.1 | C:98.0%[S:97.3%,D:0.7%],F:0.3%,M:1.7% |
| Mv-lag-1253-A2 | GCA_900013405.1 | C:98.1%[S:97.6%,D:0.5%],F:0.4%,M:1.5% |
| Mv-lat-01509-A1 | GCA_022506095.1 | C:98.0%[S:97.4%,D:0.6%],F:0.4%,M:1.6% |
| Mv-lat-01509-A2 | GCA_022506105.1 | C:97.3%[S:96.8%,D:0.5%],F:0.6%,M:2.1% |
| Mv-lyc-1064-A1 | GCA_900015465.1 | C:98.4%[S:97.1%,D:0.5%],F:0.2%,M:2.2% |
| Mv-lyc-1064-A2 | GCA_900015445.1 | C:96.9%[S:96.2%,D:0.7%],F:0.3%,M:2.8% |
| Mv-lyc-1318-A1 | GCA_003121365.1 | C:95.3%[S:94.8%,D:0.5%],F:0.3%,M:4.4% |
| Mv-lyc-1318-A2 | GCA_003121355.1 | C:91.6%[S:91.0%,D:0.6%],F:0.6%,M:7.8% |
| Mv-mel-1296-A1 | GCA_900015965.1 | C:98.0%[S:97.4%,D:0.6%],F:0.2%,M:1.8% |
| Mv-mel-1296-A2 | GCA_900011735.1 | C:98.0%[S:97.4%,D:0.6%],F:0.2%,M:1.8% |
| Mv-par-01510-A1 | GCA_022505985.1 | C:97.9%[S:97.3%,D:0.6%],F:0.3%,M:1.8% |
| Mv-par-01510-A2 | GCA_022506025.1 | C:97.8%[S:97.3%,D:0.5%],F:0.4%,M:1.8% |
| Mv-pax-1252-A1 | GCA_900015495.1 | C:97.9%[S:97.3%,D:0.6%],F:0.5%,M:1.6% |
| Mv-pax-1252-A2 | GCA_900015485.1 | C:98.0%[S:97.5%,D:0.5%],F:0.5%,M:1.5% |
| Mv-sac-1248-A1 | GCA_003665825.1 | C:98.0%[S:97.2%,D:0.8%],F:0.6%,M:1.4% |
| Mv-sac-1248-A2 | GCA_003665835.1 | C:98.1%[S:96.7%,D:1.4%],F:0.2%,M:1.7% |
| Mv-sap-1268-A1 | GCA_900015975.1 | C:98.2%[S:97.2%,D:1.0%],F:0.4%,M:1.4% |
| Mv-sap-1268-A2 | GCA_900015475.1 | C:98.3%[S:97.4%,D:0.9%],F:0.4%,M:1.3% |
| Mv-sca-1118-A1 | GCA_900008855.1 | C:97.9%[S:97.3%,D:0.6%],F:0.5%,M:1.6% |
| Mv-sca-1118-A2 | GCA_900015415.1 | C:97.9%[S:97.3%,D:0.6%],F:0.5%,M:1.6% |
| Mv-tat-1400-A1 | GCA_022702585.1 | C:98.1%[S:97.7%,D:0.4%],F:0.6%,M:1.46% |
| Mv-tat-1400-A2 | GCA_022702565.1 | C:97.8%[S:97.2%,D:0.6%],F:0.6%,M:1.6% |
| Mv-vis-01506-A1 | GCA_022505995.1 | C:97.7%[S:97.2%,D:0.5%],F:0.3%,M:2.0% |
| Mv-vis-01506-A2 | GCA_022506015.1 | C:97.7%[S:97.2%,D:0.5%],F:0.3%,M:2.0% |
| Mv-cat-1212-A1 | SAMN44081838 | C:85.0%[S:83.2%,D:1.8%],F:0.6%,M:14.4%,n:1764 |
| Mv-cat-1212-A2 | SAMN44081838 | C:81.5%[S:80.4%,D:1.1%],F:0.7%,M:17.8%,n:1764 |
| Mv-sup-1065-A1 | GCA_049639085.1 | C:97.6%[S:95.6%,D:2.0%],F:0.2%,M:2.2%,n:1764 |
| Mv-sup-1065-A2 | GCA_049639065.1 | C:97.8%[S:96.8%,D:1.0%],F:0.5%,M:1.7%,n:1764 |
| M-scorzo-A1 | GCA_049638985.1 | C:97.8%[S:97.3%,D:0.5%],F:0.6%,M:1.6%,n:1764 |
| M-scorzo-A2 | GCA_049638945.1 | C:97.5%[S:97.0%,D:0.5%],F:0.6%,M:1.9%,n:1764 |
| MvSn-1249-A1-R1 | GCA_900015425.1 | C:95.5%[S:94.8%,D:0.7%],F:0.2%,M:4.3%,n:1764 |
| MvSn-1249-A2-R1 | GCA_900015455 | C:95.0%[S:94.5%,D:0.5%],F:0.3%,M:4.7%,n:1764 |

**Table S6 :** mean depth of RNAseq mappings on the A1 genome of *Microbotryum lychnidis-dioicae* across the different RNAseq conditions. Sequence data taken from PRJNA246 470

**A)**

| contig | Replicate1 | Replicate2 | Replicate3 |
|---|---|---|---|
| Mlyc1064a1_A1 | 137.8 | 154.5 | 147.6 |
| Mlyc1064a1_MC01-1 | 201.6 | 204.0 | 192.3 |
| Mlyc1064a1_MC01-2 | 190.6 | 180.5 | 169.2 |
| Mlyc1064a1_MC02 | 205.1 | 207.3 | 196.7 |
| Mlyc1064a1_MC03 | 183.9 | 189.3 | 177.2 |
| Mlyc1064a1_MC04-1 | 165.0 | 164.6 | 155.2 |
| Mlyc1064a1_MC04-2 | 88.2 | 88.3 | 79.8 |
| Mlyc1064a1_MC05-1 | 139.7 | 138.6 | 126.7 |
| Mlyc1064a1_MC05-2 | 190.6 | 199.1 | 180.3 |
| Mlyc1064a1_MC05-3 | 227.3 | 234.9 | 224.2 |
| Mlyc1064a1_MC06 | 208.9 | 214.9 | 202.8 |
| Mlyc1064a1_MC07 | 222.1 | 228.9 | 214.8 |
| Mlyc1064a1_MC08 | 225.9 | 225.1 | 217.3 |
| Mlyc1064a1_MC09-1 | 55.7 | 50.0 | 50.4 |
| Mlyc1064a1_MC09-2 | 210.7 | 216.6 | 203.5 |
| Mlyc1064a1_MC10-1 | 191.1 | 188.6 | 175.3 |
| Mlyc1064a1_MC10-2 | 264.5 | 249.1 | 230.2 |
| Mlyc1064a1_MC11 | 185.8 | 183.2 | 173.9 |
| Mlyc1064a1_MC12 | 242.6 | 242.9 | 231.2 |
| Mlyc1064a1_MC13 | 218.8 | 225.5 | 211.2 |
| Mlyc1064a1_MC14 | 150.2 | 160.2 | 151.3 |
| Mlyc1064a1_MC15 | 185.0 | 191.6 | 190.7 |
| Mlyc1064a1_MC17 | 175.9 | 162.8 | 188.0 |

**B)** mean depth of RNAseq mappings on the A2 genome of *Microbotryum lychnidis-dioicae* across the different RNAseq conditions.

| contig | Replicate1 | Replicate2 | Replicate3 |
|---|---|---|---|
| Mlyc1064a2_A2 | 114.7 | 115.0 | 106.9 |
| Mlyc1064a2_MC01 | 230.0 | 230.9 | 217.9 |
| Mlyc1064a2_MC02 | 238.6 | 241.5 | 229.3 |
| Mlyc1064a2_MC03 | 212.9 | 219.6 | 205.6 |
| Mlyc1064a2_MC04-1 | 189.7 | 190.7 | 179.3 |
| Mlyc1064a2_MC04-2 | 187.3 | 183.7 | 172.8 |
| Mlyc1064a2_MC05 | 226.1 | 234.7 | 221.3 |
| Mlyc1064a2_MC06-1 | 212.8 | 223.5 | 206.9 |
| Mlyc1064a2_MC06-2 | 248.7 | 254.0 | 238.5 |
| Mlyc1064a2_MC07 | 256.9 | 264.8 | 248.5 |
| Mlyc1064a2_MC08-1 | 270.5 | 269.7 | 257.6 |
| Mlyc1064a2_MC08-2 | 217.3 | 216.2 | 208.2 |
| Mlyc1064a2_MC09 | 246.1 | 249.8 | 233.7 |
| Mlyc1064a2_MC10 | 256.2 | 252.1 | 237.1 |
| Mlyc1064a2_MC11 | 210.2 | 211.2 | 202.6 |
| Mlyc1064a2_MC12 | 283.3 | 281.7 | 269.2 |
| Mlyc1064a2_MC13 | 254.5 | 262.7 | 246.4 |
| Mlyc1064a2_MC14 | 177.0 | 190.8 | 175.8 |
| Mlyc1064a2_MC15 | 203.5 | 210.6 | 197.1 |
| Mlyc1064a2_MC16 | 54.4 | 60.7 | 51.2 |
| Mlyc1064a2_MC17 | 179.4 | 166.2 | 159.2 |
| Mlyc1064a2_MC18 | 2.4 | 2.9 | 2.7 |

**Table S7: Model weight obtained in *Microbotryum v. caroliniana*, *M. v. tatarinowii* and *M. lychnidis dioicae.*** Models with the highest weight are displayed in bold.

| | *M. v. caroliniana* | | | *M. v. tatarinowii* | | | *M. lychnidis dioicae* | | |
|---|---|---|---|---|---|---|---|---|---|
| | **elpd.diff** | **se.diff** | **weight** | **elpd.diff** | **se.diff** | **weight** | **elpd.diff** | **se.diff** | **weight** |
| ***Model1 (3 cp)*** | −17.1 | 7.8 | 0 | −21.63 | 14.98 | 0.003 | −10.38 | 7.59 | 0.05 |
| ***Model2 (4cp)*** | −7.8 | 8.5 | 0.082 | −10.64 | 12.45 | 0.16 | −7.45 | 5.80 | 0.046 |
| ***Model3 (5cp)*** | −1.8 | 7.1 | **0.339** | −18.25 | 2.76 | 0 | −4.05 | 2.94 | 0.030 |
| ***Model4 (6cp)*** | −12.9 | 8.7 | 0.002 | 0 | 0 | **0.52** | −2 | 2.8 | **0.165** |
| ***model5 (7cp)*** | **0** | **0** | **0.577** | −19.74 | 11.24 | 0.01 | −3.02 | 1.05 | 0.03 |
| ***Model6 (8cp)*** | Not tested | Not tested | Not tested | −2.77 | 4.12 | **0.27** | **0** | **0** | **0.55** |
| ***Moldel7 (9 cp)*** | Not tested | Not tested | Not tested | −22.90 | 5.06 | 1.6e-8 | −2.65 | 1.90 | 0.12 |

**Table S8: Example table from the summary function for the 'most likely' model with 7 changepoint in *Microbotryum violaceum caroliniana***

cp_1 to cp7 = changepoint location (along the gene order) with the mean location of the changepoint and lower and upper value. Rhat = n.eff = proportion of the MCMC contributing to the changepoint.

int_1 to int_8 = dS value along each interval in-between the respective changepoint.

Sigma_1 = variance parameter of the model.

Each of the tables is generated automatically for each changepoint.

| *name* | *mean* | *lower* | *upper* | *Rhat* | *n.eff* |
|---|---|---|---|---|---|
| *cp_1* | 8.00E+01 | 71.5275 | 85.998 | 1 | 13 374 |
| *cp_2* | 1.30E+02 | 120.7824 | 134.001 | 1 | 1252 |
| *cp_3* | 1.50E+02 | 145.0044 | 170.697 | 1.1 | 128 |
| *cp_4* | 2.10E+02 | 195.6807 | 224.99 | 1 | 1513 |
| *cp_5* | 2.50E+02 | 239.8215 | 259.917 | 1 | 24 238 |
| *cp_6* | 2.80E+02 | 281.0026 | 282 | 1 | 11 745 |
| *cp_7* | 3.30E+02 | 325.1766 | 344.867 | 1 | 24 039 |
| *int_1* | 2.10E-03 | −0.0065 | 0.01 | 1 | 23 598 |
| *int_2* | 3.30E-02 | 0.0235 | 0.042 | 1 | 18 922 |
| *int_3* | 8.50E-02 | 0.0645 | 0.103 | 1 | 474 |
| *int_4* | 3.80E-02 | 0.0253 | 0.051 | 1 | 2075 |
| *int_5* | 7.20E-02 | 0.0443 | 0.108 | 1.1 | 3422 |
| *int_6* | 7.80E-02 | 0.0578 | 0.097 | 1 | 22 074 |
| *int_7* | 5.70E-03 | −0.003 | 0.014 | 1 | 24 962 |
| *int_8* | 9.90E-04 | −0.0095 | 0.012 | 1 | 24 749 |
| *sigma_1* | 2.80E-02 | 0.0247 | 0.031 | 1 | 11 464 |

**Table S9: Bayes Factor testing for the evidence of different evolutionary strata in *Microbotryum violaceum caroliniana***

Column "Hypothesis" displays the directionality of the difference in $d_S$ values being tested. Mean, lower and upper are the differences in $d_S$ values. P = posterior probability of the difference being significant, BF = Bayes Factor.
Interval 1 and Interval 8 correspond to the pseudoautosomal region freely recombining.
Interval 5 and 6 are similar, supporting results previously reported (Branco et al. 2017).
Interval 7 is only modestly different from the recombining region.

Such tables are automatically computed for each direction of difference and each interval, but we encourage users to investigate more subtle hypotheses manually.

| *hypothesis* | *mean* | *lower* | *upper* | *p* | *BF* |
|---|---|---|---|---|---|
| *Int_1 – int_2 <0* | −0.0307578 | −0.043094 | −0.0178699 | 1 | Inf |
| *Int_2 – int_3 < 0* | −0.0519023 | −0.071797 | −0.0300964 | 1 | Inf |
| *Int_3 – int_4 > 0* | 0.0470306 | 0.026117 | 0.0673731 | 1 | Inf |
| *Int_5 – int_4 > 0* | 0.0340599 | 0.003354 | 0.0679488 | 0.988925 | 89.2934 |
| *Int_6 – int_5 > 0* | 0.0057295 | −0.033658 | 0.0412362 | 0.652475 | 1.87749 |
| *Int_6 – int_7 > 0* | 0.0718367 | 0.050656 | 0.0931820 | 1 | Inf |
| *Int_7 – int_8 > 0* | 0.0046758 | −0.009170 | 0.0181581 | 0.749625 | 2.99401 |

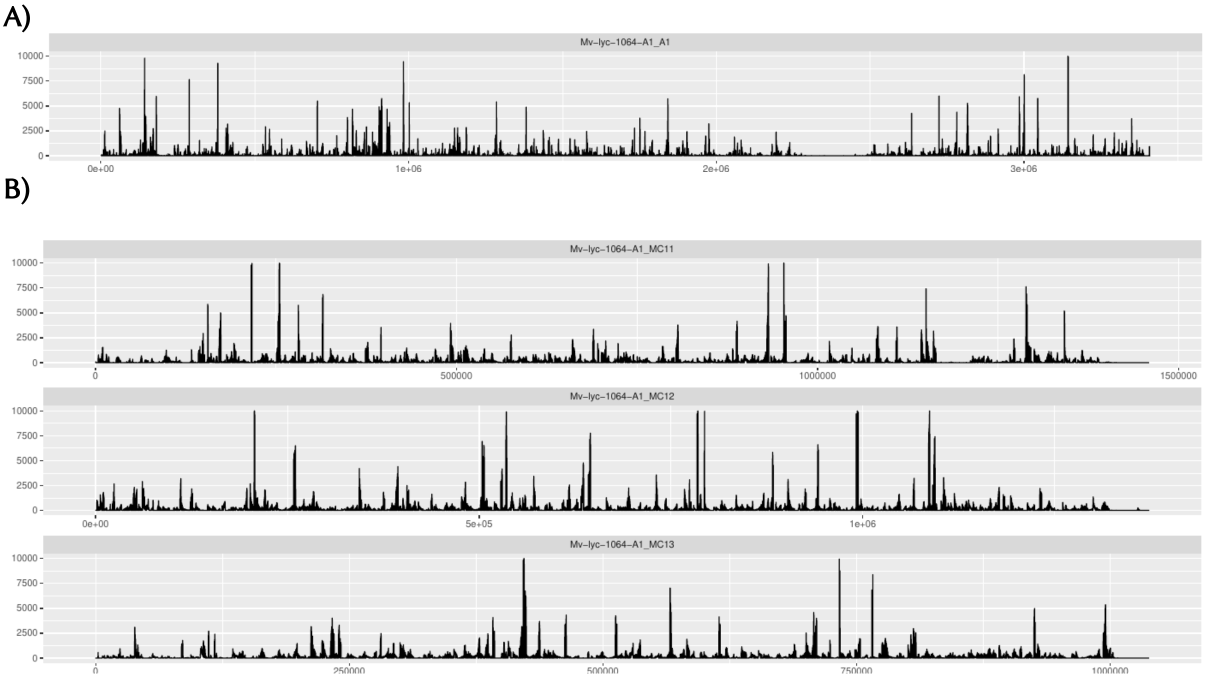


**Figure S1:** Read-depth variation for a single condition along (A) the $a_1$ mating-type chromosome and (B) along three autosomes. Each panel represents a chromosome, read depth is shown along the Y axis while the X axis represents genomic positions in base pairs. The workflow automatically produces these plots for each condition/replicates and each chromosome as well as tables of mean read depth per chromosome.

A)

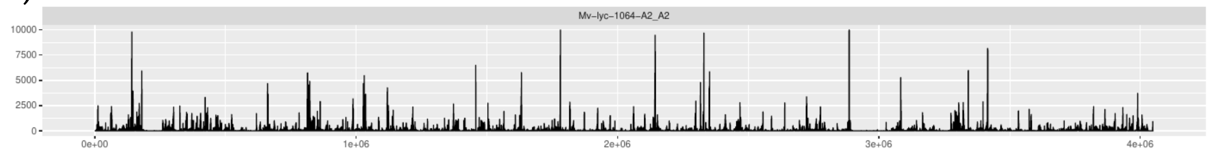


B)

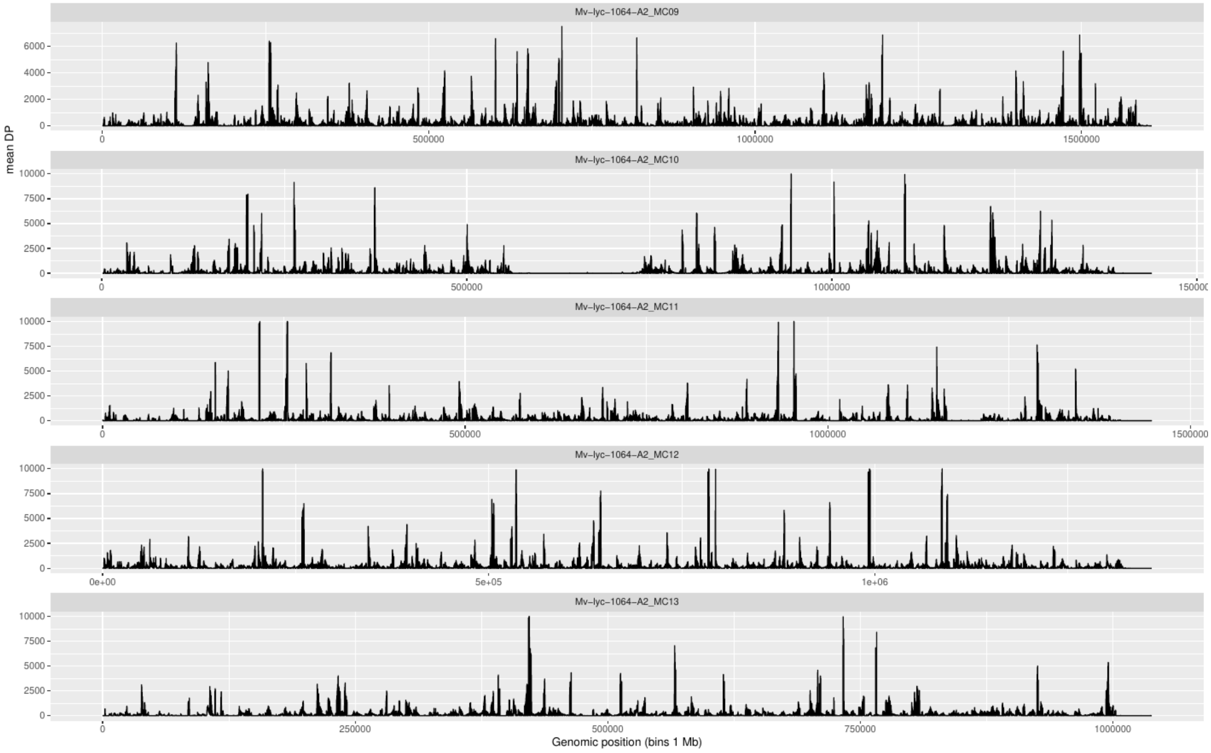


**Figure S2 :** Read-depth variation for a single condition along A) the $a_2$ mating-type chromosome and (B) along four autosomes. Each panel represents a chromosome, read depth is shown along the Y axis while the X axis represents genomic positions in base pairs. The workflow automatically produces these plots for each condition/replicates and each chromosome as well as tables of mean read depth per chromosome.

A)

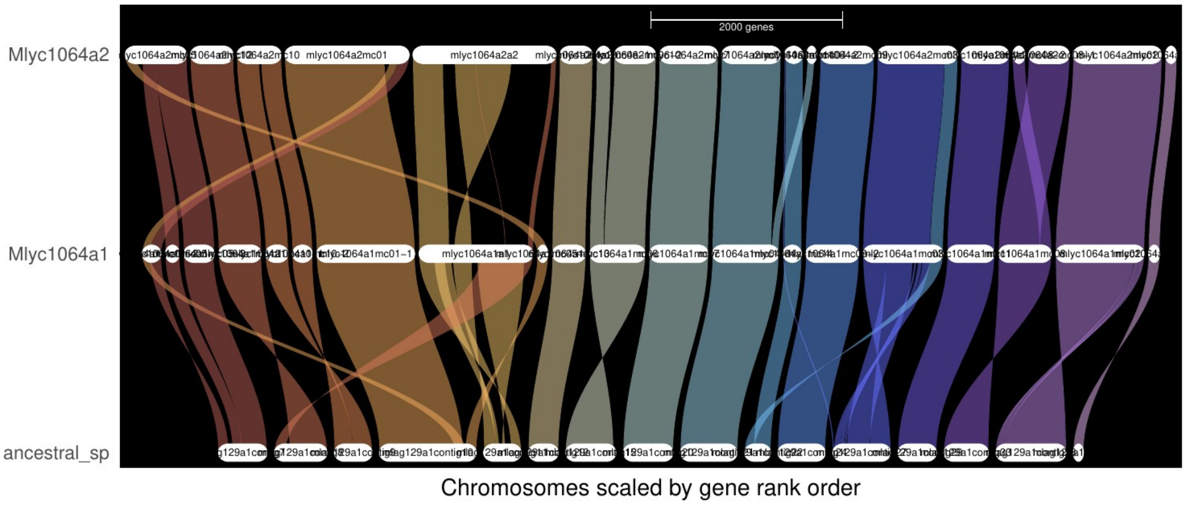


B)

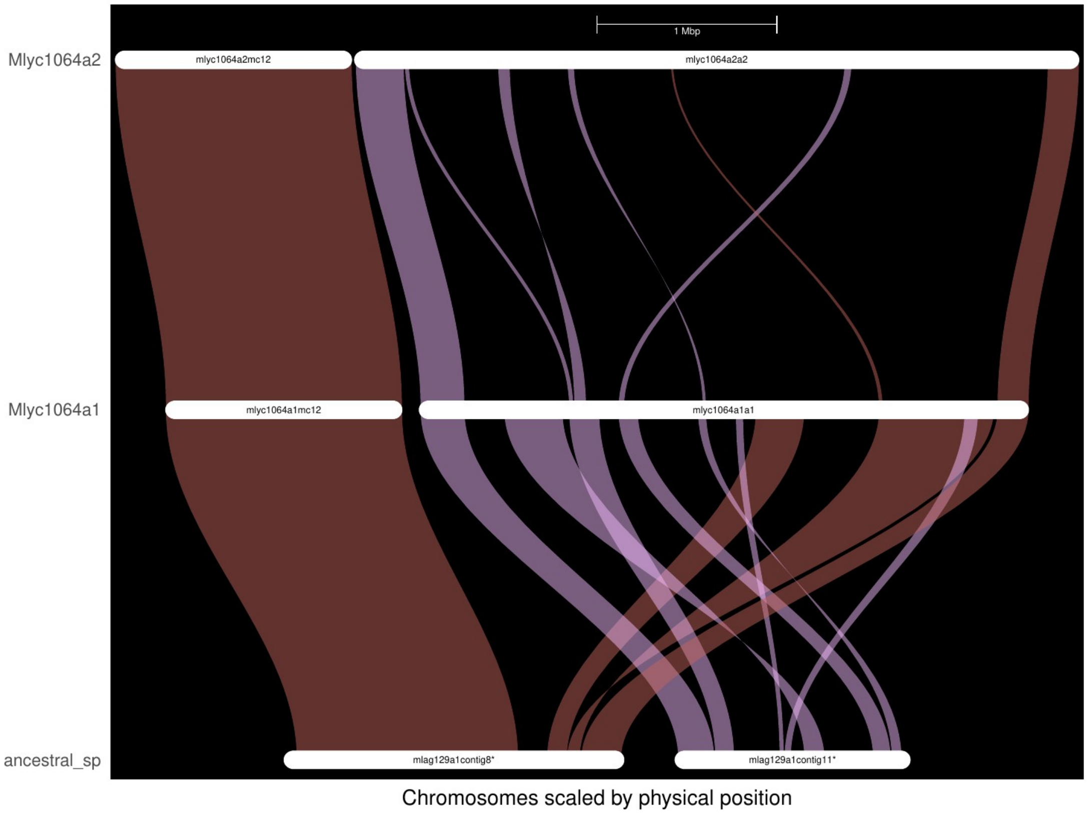


**Figure S3: GeneSpace Plots**

**A) Synteny comparison between the whole genomes of the two mating types and the species with an ancestral-like genome arrangement.** As expected, the colinearity is preserved between the two genomes of opposite mating types, except along the sex chromosomes. Genes are arranged according to their order along the sequence. Colours represent synteny blocks from different chromosomes. “Ancestral_sp” = species used as a proxy for the ancestral gene order. Mlyc1064a1 and Mlyc1064a2 = short name for *Microbotryum lychnidis-dioicae* strain 1064 mating-type $a_1$ or $a_2$

**B) Plot focussing on the mating-type chromosomes ($a_1$ and $a_2$) obtained with the plot_riparian GENESPACE function.**

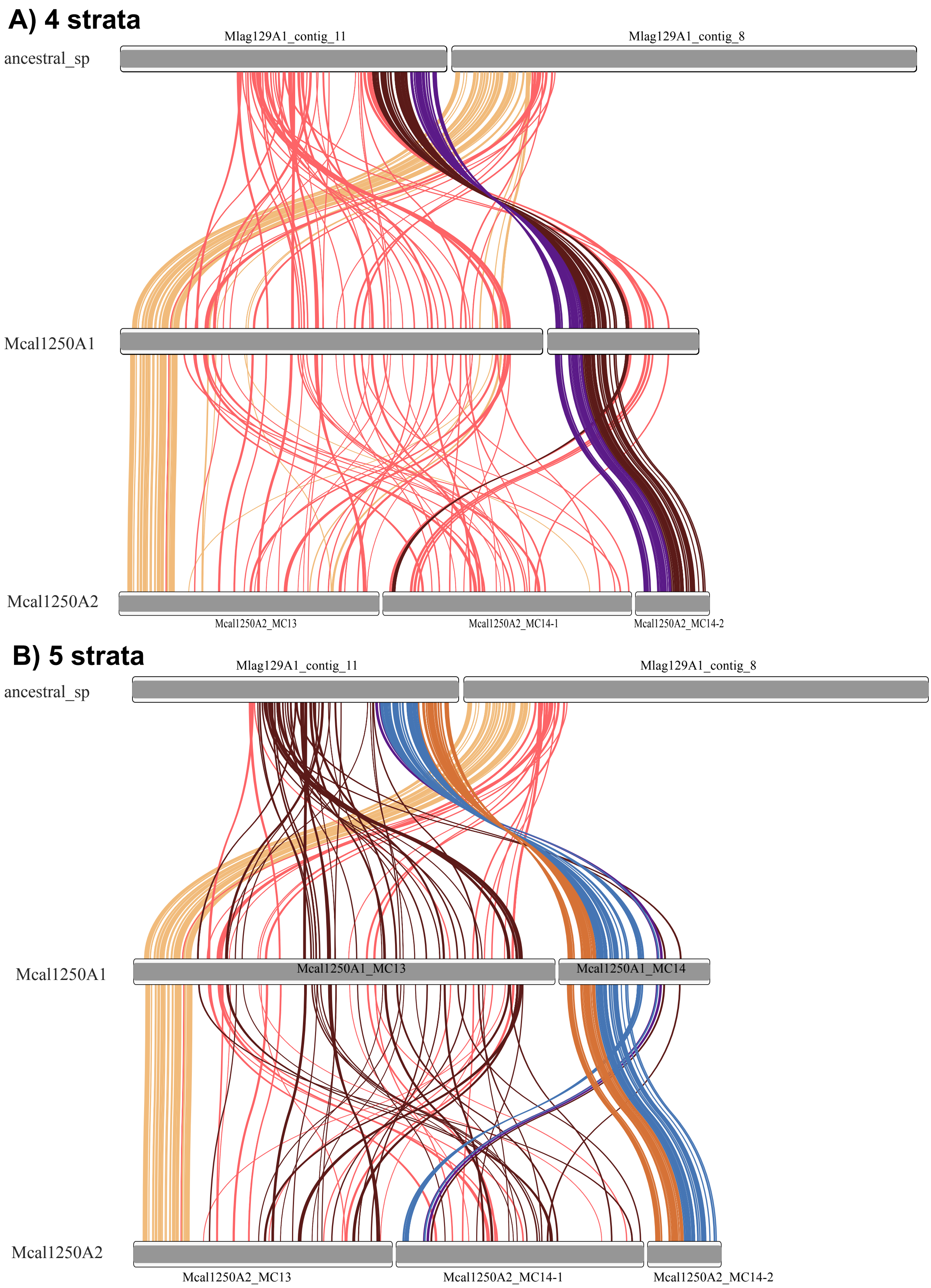


**Figure S4:** Ideogram colored by strata in *Microbotryum violaceum caroliniana (Mcal1250a1 and Mcal1250a2)*, with synteny between the $a_1$ and $a_2$ mating-type chromosomes, and compared to the ancestral-like mating-type chromosome a*1* in *M. lagerheimii*

*(ancestral_sp).* Such plots are generated automatically for a range of changepoint values from 4 to 9 changepoints in the workflow.

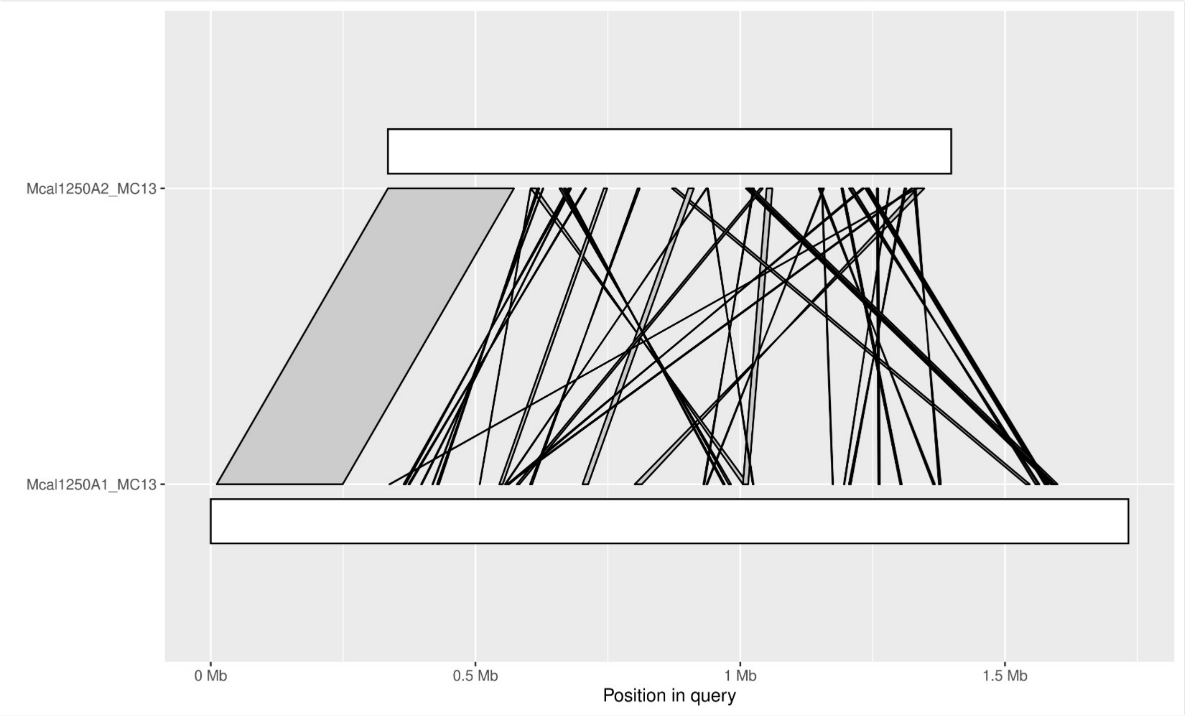


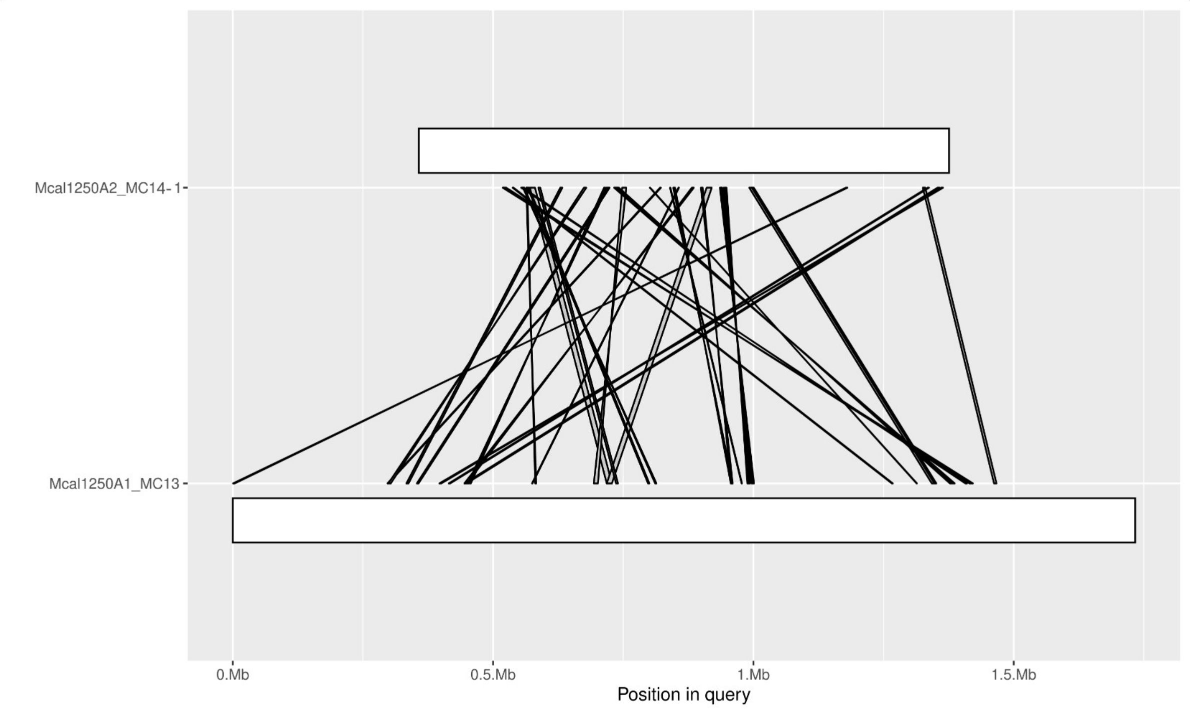


**Figure S5** Example of synteny plots between two contigs from the mating type chromosomes in *Microbotryum violaceum caroliniana (Mcal1250a1 and Mcal1250a2)*,. All such contigs are plotted automatically and can be exploited to identify sex chromosomes, pseudo-autosomal regions (PARs) and extremely rearranged regions such as the one displayed here. The suffix "_MCXX" refers to the different contigs.

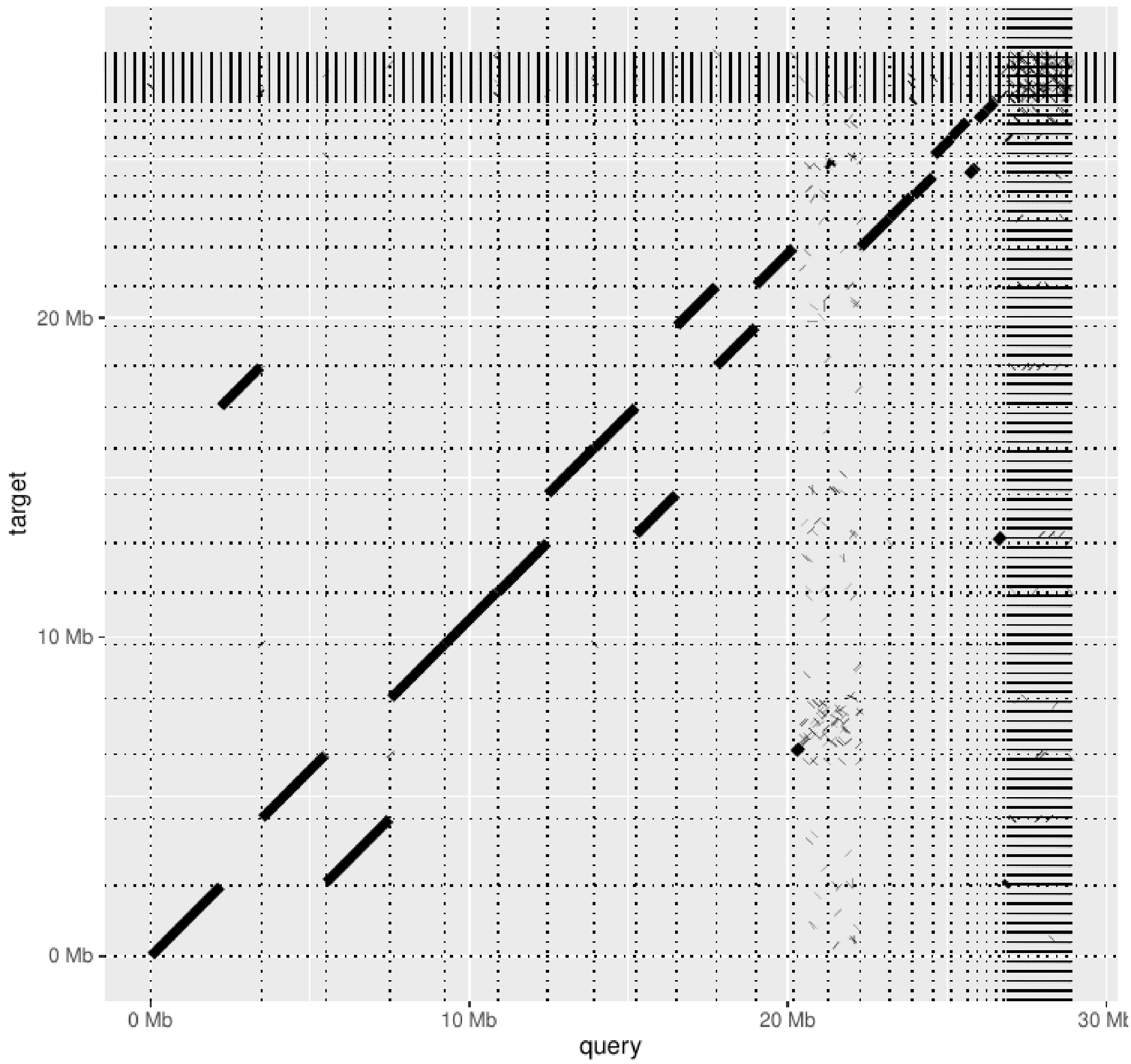


**Figure S6:** Example dotplot between *Microbotryum violaceum caroliniana,* $a_1$ and $a_2$ whole genome assemblies. The completely rearranged region corresponds to mating type chromosomes.

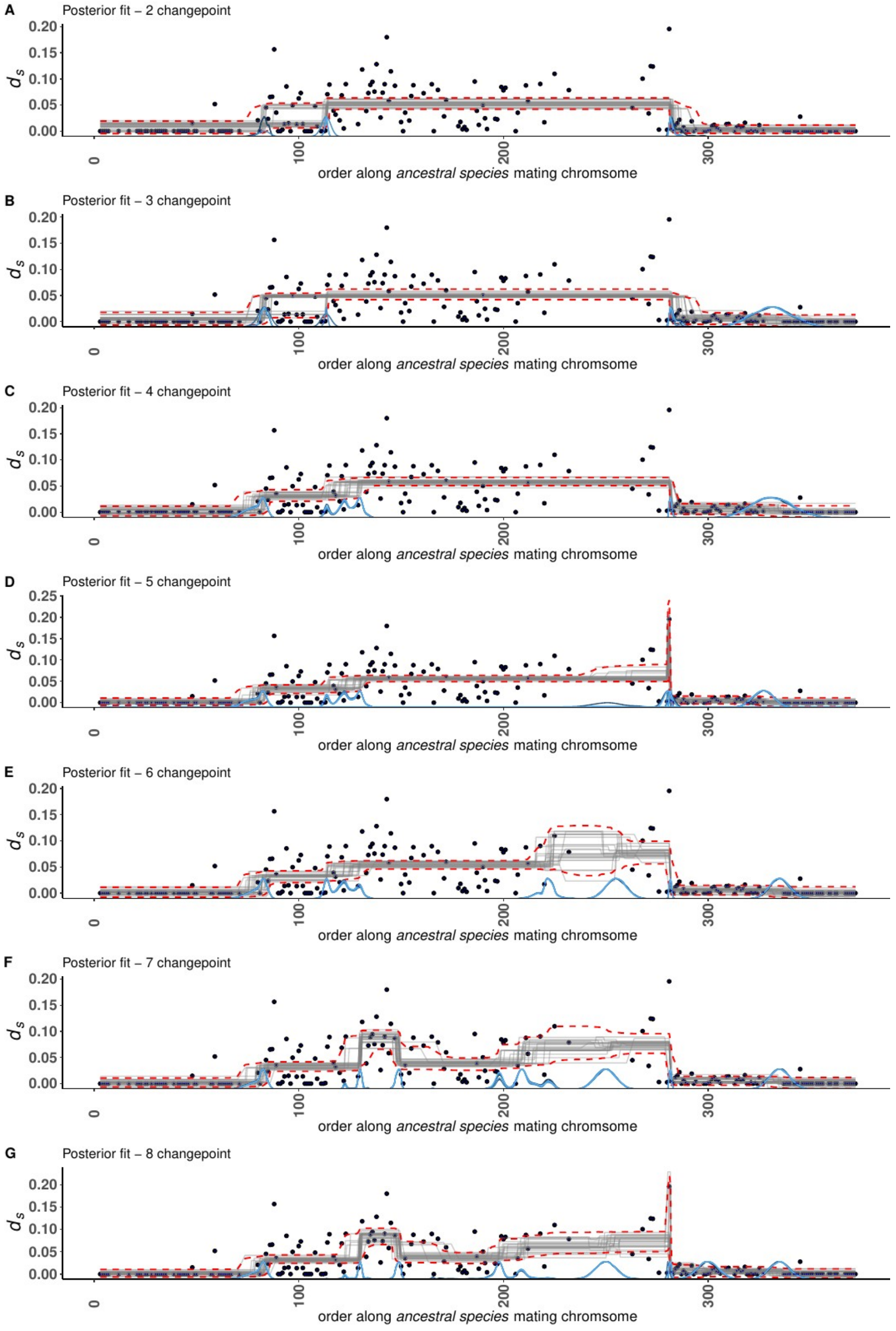


**Figure S7: Changepoint details in *Microbotryum violaceum caroliniana* (MCP results for 3 to 8 evolutionary strata).** Each changepoint panel displays the distribution of raw data (i.e., $d_S$ values as black dots) along with 25 draws from the joint posterior distribution (grey lines) and 95% highest density interval (red lines). Posterior distributions of the changepoints are shown in blue with one line for each chain. Posterior fits are displayed for models with 5, 6 and 7 changepoints, corresponding to 3, 4 and 5 evolutionary strata, respectively. Such plots are generated automatically for a range of changepoint values from 3 to 9 changepoints in the workflow.

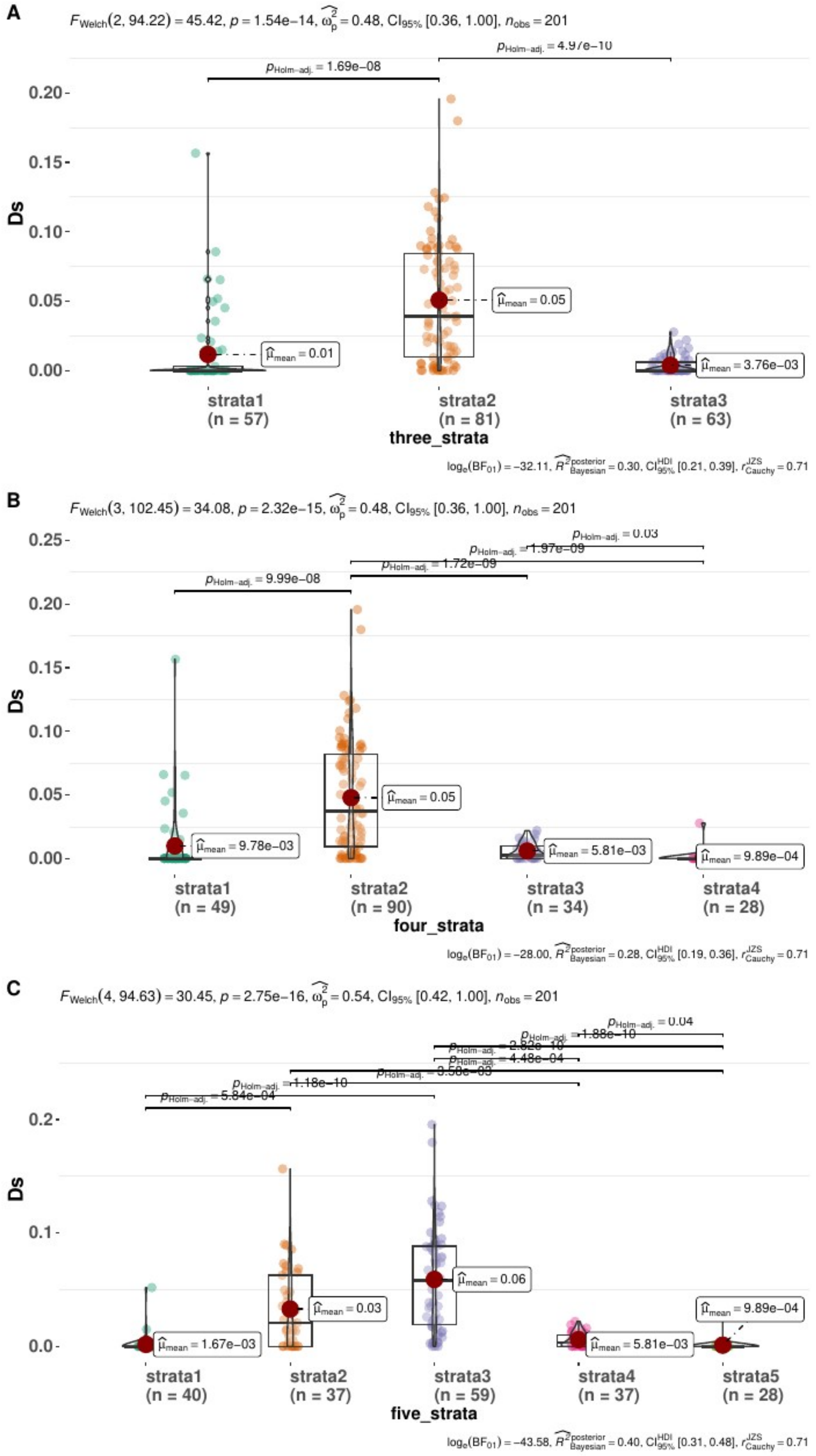


**Figure S8: Violin plots of $d_S$ values for three to five evolutionary strata in *Microbotryum violaceum caroliniana.*** Plots obtained with default parameters from the ggbetweenstats function in the ggstatsplot package. Such plots are generated automatically for a range of changepoint values from 3 to 9 changepoints in the workflow.

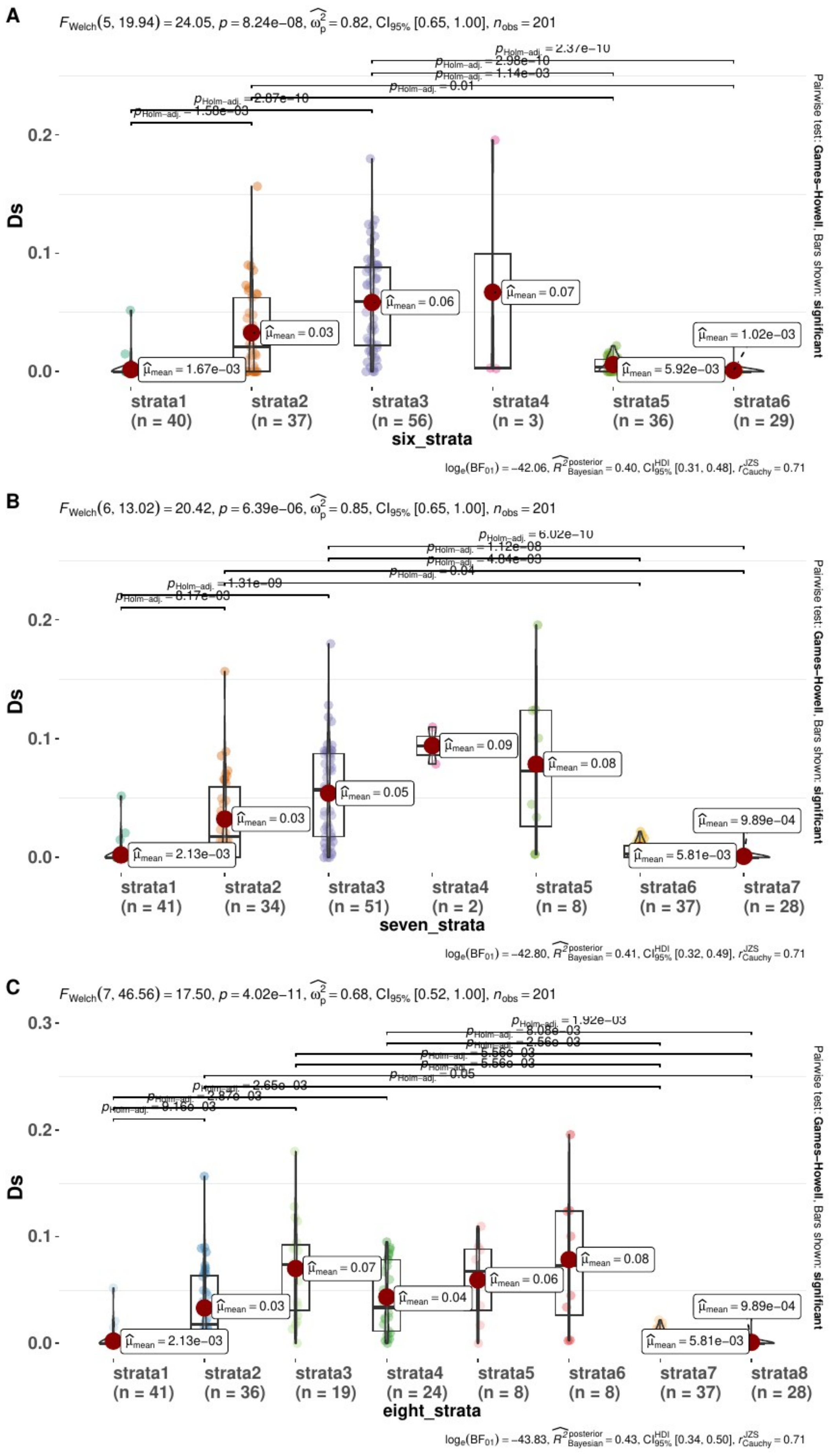
A
$F_{Welch}(5, 19.94) = 24.05$, $p = 8.24e-08$, $\widehat{\omega_p^2} = 0.82$, $CI_{95\%}$ [0.65, 1.00], $n_{obs} = 201$
Ds
strata1 (n = 40)
strata2 (n = 37)
strata3 (n = 56)
strata4 (n = 3)
strata5 (n = 36)
strata6 (n = 29)
six_strata
Pairwise test: Games-Howell, Bars shown: significant
$\log_e(BF_{01}) = -42.06$, $\widehat{R^2}^{posterior}_{Bayesian} = 0.40$, $CI^{HDI}_{95\%}$ [0.31, 0.48], $r^{JZS}_{Cauchy} = 0.71$
B
$F_{Welch}(6, 13.02) = 20.42$, $p = 6.39e-06$, $\widehat{\omega_p^2} = 0.85$, $CI_{95\%}$ [0.65, 1.00], $n_{obs} = 201$
Ds
strata1 (n = 41)
strata2 (n = 34)
strata3 (n = 51)
strata4 (n = 2)
strata5 (n = 8)
strata6 (n = 37)
strata7 (n = 28)
seven_strata
Pairwise test: Games-Howell, Bars shown: significant
$\log_e(BF_{01}) = -42.80$, $\widehat{R^2}^{posterior}_{Bayesian} = 0.41$, $CI^{HDI}_{95\%}$ [0.32, 0.49], $r^{JZS}_{Cauchy} = 0.71$
C
$F_{Welch}(7, 46.56) = 17.50$, $p = 4.02e-11$, $\widehat{\omega_p^2} = 0.68$, $CI_{95\%}$ [0.52, 1.00], $n_{obs} = 201$
Ds
strata1 (n = 41)
strata2 (n = 36)
strata3 (n = 19)
strata4 (n = 24)
strata5 (n = 8)
strata6 (n = 8)
strata7 (n = 37)
strata8 (n = 28)
eight_strata
Pairwise test: Games-Howell, Bars shown: significant
$\log_e(BF_{01}) = -43.83$, $\widehat{R^2}^{posterior}_{Bayesian} = 0.43$, $CI^{HDI}_{95\%}$ [0.34, 0.50], $r^{JZS}_{Cauchy} = 0.71$

**Figure S9: Violin plots for numbers of evolutionary strata from 6 to 8.** Plots obtained with default parameters from the ggbetweenstats function in ggstatsplot package. Such plots are generated automatically for a range of changepoint values from 3 to 9 changepoints in the workflow.

**Figure S10:** _Per-gene dS values plotted along gene order, colored according to two evolutionary strata (i.e. four changepoints) to five evolutionary strata (i.e. seven changepoints) in *Microbotryum violaceum caroliniana.* The dS values at zero at the

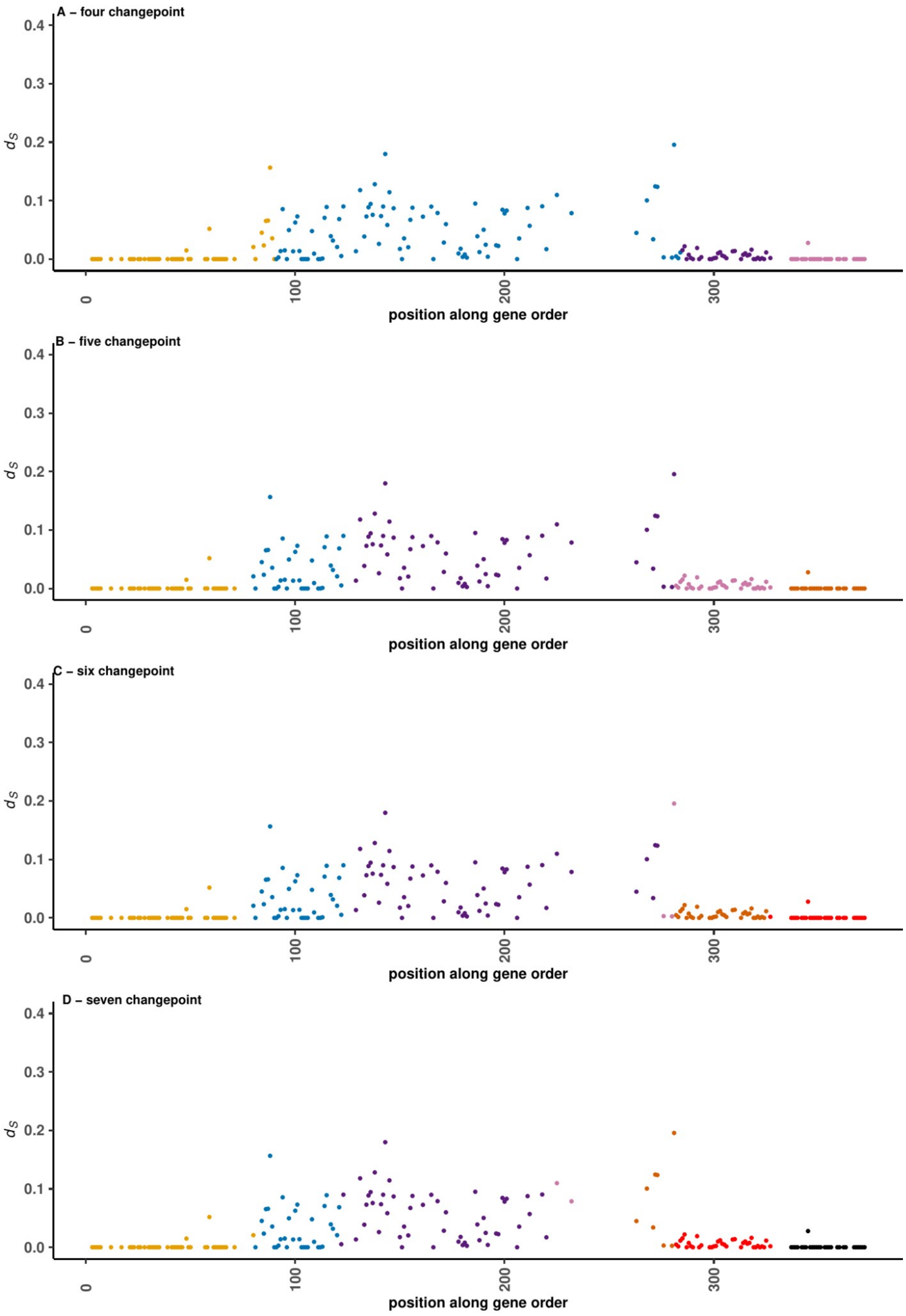


beginning and end of the plot correspond to the pseudo-autosomal regions (PARs). Such plots are generated automatically for a range of changepoint values from 3 to 9 changepoints

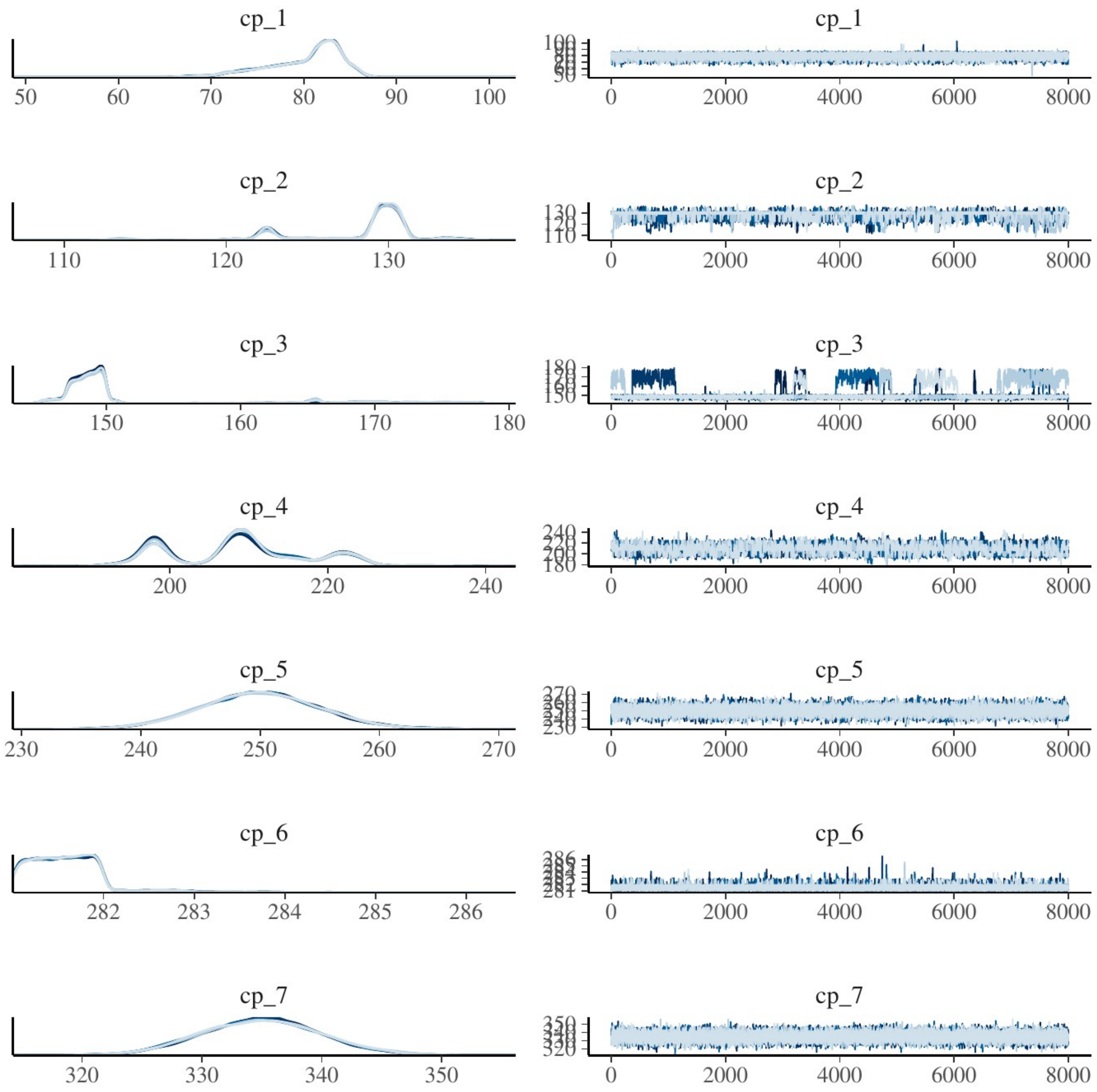


**Figure S11: MCP convergence of $d_S$ values for inferring evolutionary strata in *Microbotryum violaceum caroliniana* for the best models.** Such plots are generated automatically for a range of changepoint values from 3 to 9 changepoints in our workflow.

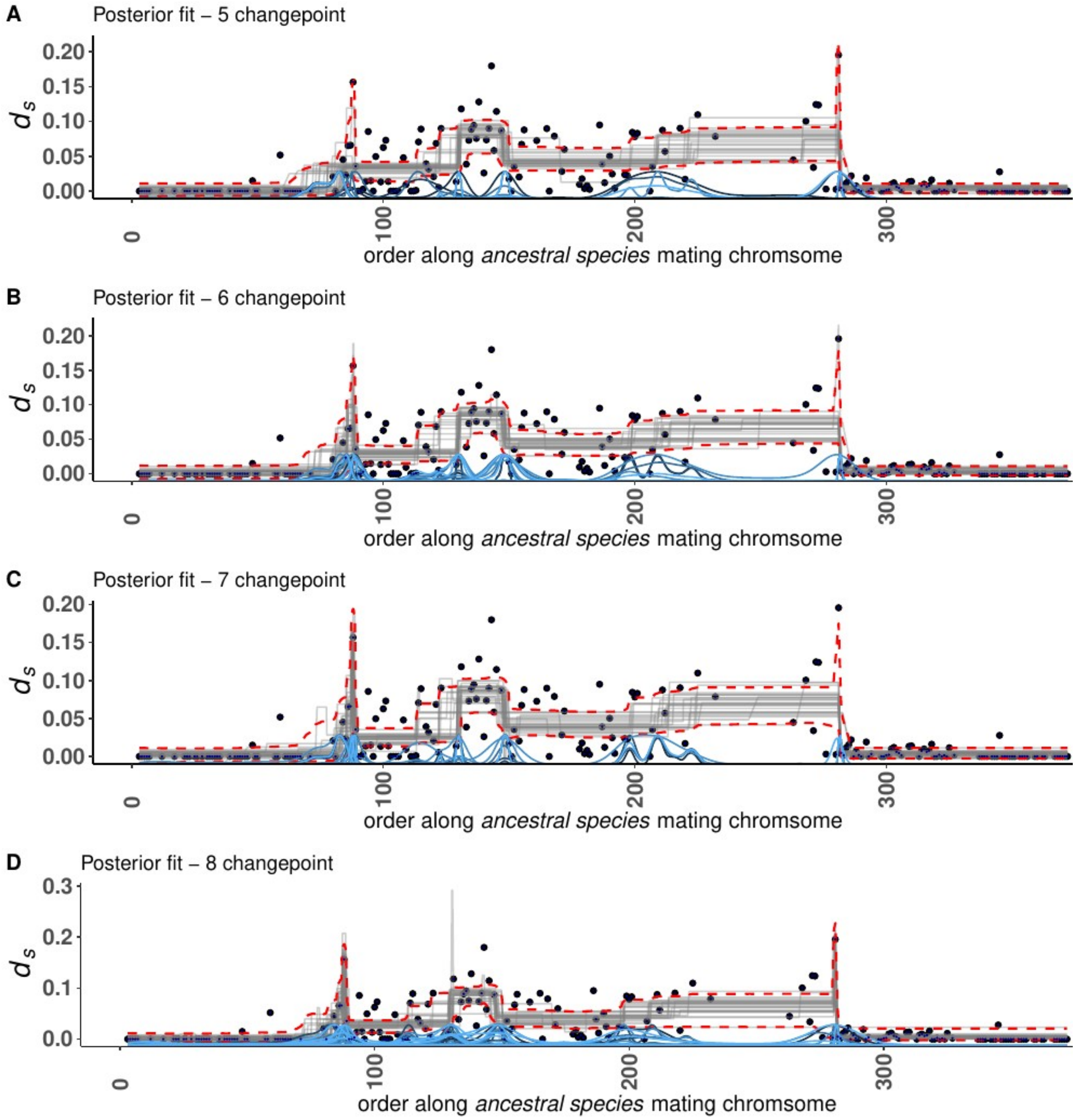


**Figure S12 : MCP figure without priors for inferring evolutionary strata in *Microbotryum violaceum caroliniana.*** Such plots are generated automatically for a range of changepoint values, from 3 to 9 changepoints in our workflow.

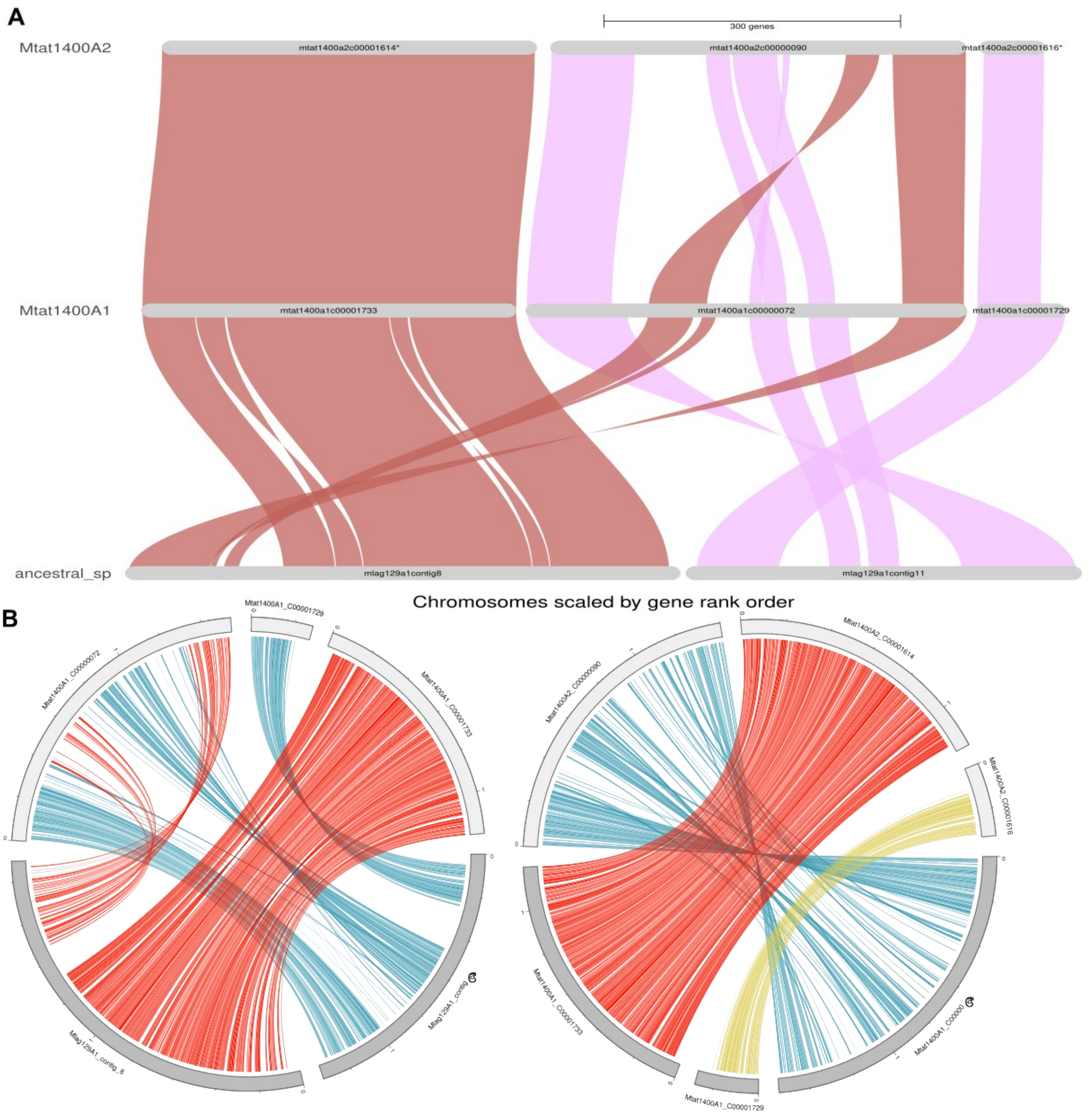


**Figure S13: GeneSpace (A) and Circos plots (B) showing synteny and orthology relationship between among species and mating-type chromosomes of *Microbotryum lagerheimii (ancestral_sp on the plots)* and *M. v. tatarinowii 1400* $a_1$ (Mtat1400a1) and between the two mating-type chromosomes of *M. v. tatarinowii 1400* ($a_1$ and $a_2$, Mtat1400a2).** Mlag129A1_contig 8 corresponds to the HD chromosome and Malg129A1contig11 to the PR chromosome.

**A)** Plots showing major synteny blocks as inferred by combining orthofinder, blasts and MCScan results. Synteny groups must include at least five consecutive genes.

**B)** Circos plots of the *M. v. tatarinowii 1400* $a_1$ mating-type chromosomes compared to the *M. lagerheimii* HD and PR mating-type chromosomes and *M. v. tatarinowii 1400* $a_1$ versus *M. v. tatarinowii 1400* $a_2$ mating-type chromosomes. Red link = single-copy orthologous genes between PR contigs. Blue links = single-copy orthologs genes between HD contigs. Colors in the inner track display the PR/HD and pheromone genes and their links between their positions in the two genomes. Inner circle displays gene density in light blue and TE density in spring green respectively.

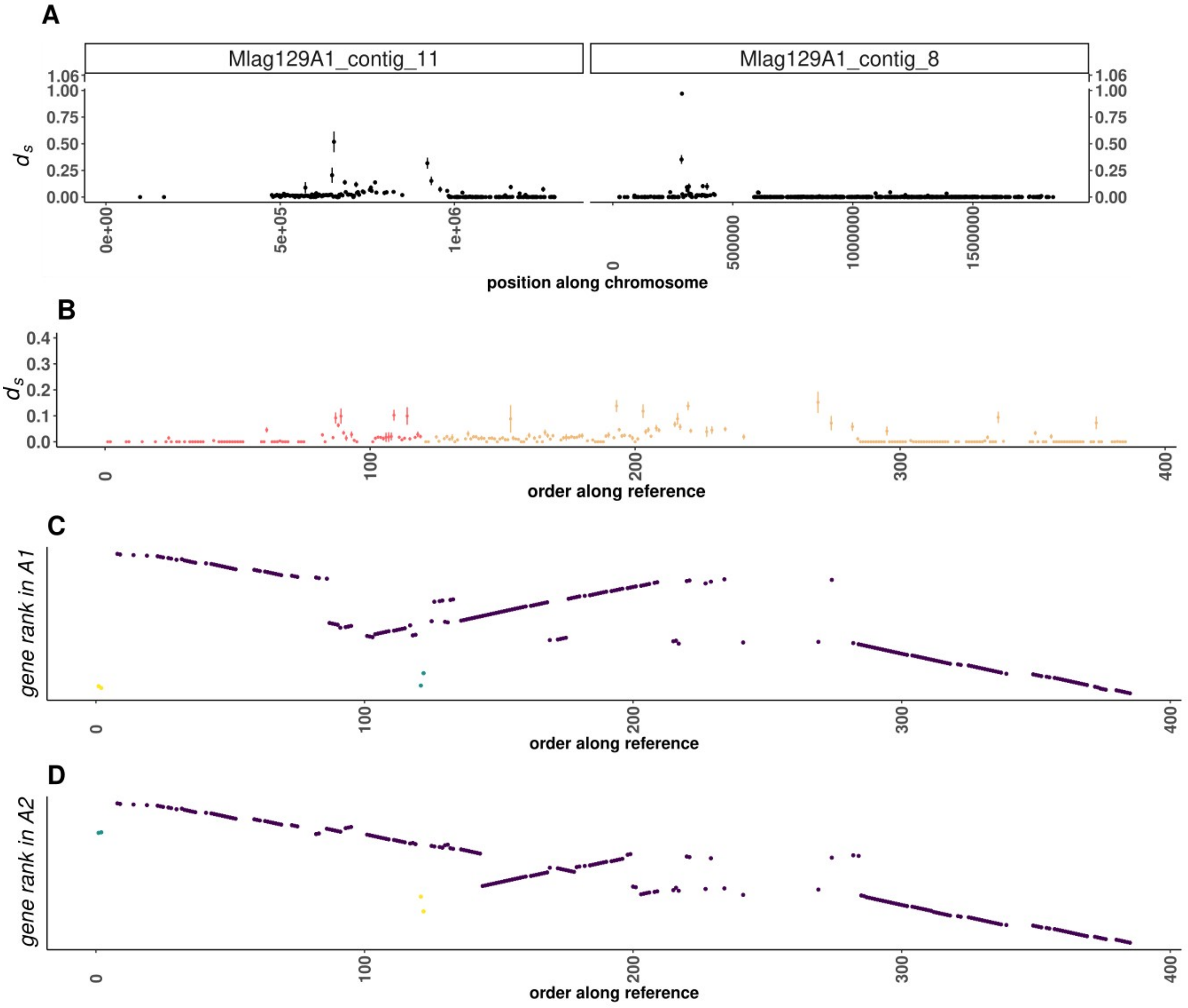


**Figure S14: Observed $d_S$ values and rearrangements in the *Microbotryum v. tatarinowii* case. A)** observed $d_S$ values between the *M. v. tatarinowii* $a_1$ and $a_2$ mating-type chromosomes along the mating-type chromosomes of *M. lagerheimii*, with the contig 11 corresponding to the PR chromosome and the contig 8 corresponding to the HD chromosome. B) Distribution of the $d_S$ values considering the ancestral gene order instead of the physical position along the *M. lagerheimii* mating-type chromosomes. Contigs are reversed to match the expected order of the chromosomal rearrangement and fusion in *M. v. tatarinowii*. The large region in contig 8 that became an autosome upstream of the centromere in panel A (>500 kbp) has been removed. C) and D) Current distribution of the rank of the genes in the $a_1$ and $a_2$ mating-type chromosomes along the ancestral gene order, showing the rearrangements.

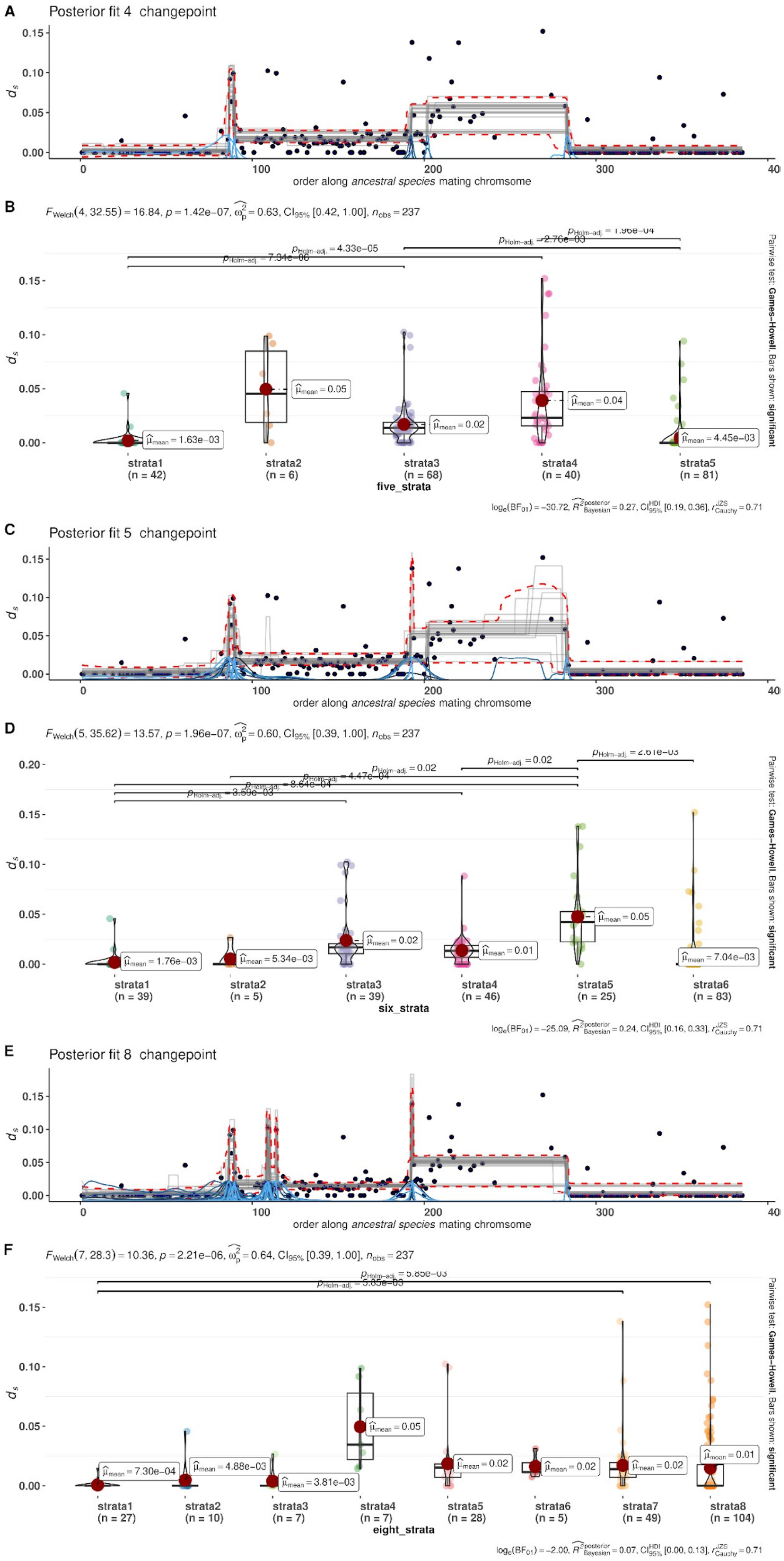

A
Posterior fit 4 changepoint
0.15
0.10
0.05
0.00
ds
0
100
200
300
400
order along ancestral species mating chromsome
B
FWelch(4, 32.55) = 16.84, p = 1.42e-07, ωp2 = 0.63, CI95% [0.42, 1.00], nobs = 237
pHolm-adj. = 1.96e-04
pHolm-adj. = 2.76e-03
pHolm-adj. = 4.33e-05
pHolm-adj. = 7.34e-06
μmean = 1.63e-03
μmean = 0.05
μmean = 0.02
μmean = 0.04
μmean = 4.45e-03
strata1 (n = 42)
strata2 (n = 6)
strata3 (n = 68)
strata4 (n = 40)
strata5 (n = 81)
five_strata
Pairwise test: Games-Howell; Bars shown: significant
loge(BF01) = −30.72, R2posterior Bayesian = 0.27, CIHDI95% [0.19, 0.36], rJZS Cauchy = 0.71
C
Posterior fit 5 changepoint
order along ancestral species mating chromsome
D
FWelch(5, 35.62) = 13.57, p = 1.96e-07, ωp2 = 0.60, CI95% [0.39, 1.00], nobs = 237
pHolm-adj. = 2.61e-03
pHolm-adj. = 0.02
pHolm-adj. = 0.02
pHolm-adj. = 4.47e-04
pHolm-adj. = 8.64e-04
pHolm-adj. = 3.59e-03
μmean = 1.76e-03
μmean = 5.34e-03
μmean = 0.02
μmean = 0.01
μmean = 0.05
μmean = 7.04e-03
strata1 (n = 39)
strata2 (n = 5)
strata3 (n = 39)
strata4 (n = 46)
strata5 (n = 25)
strata6 (n = 83)
six_strata
Pairwise test: Games-Howell; Bars shown: significant
loge(BF01) = −25.09, R2posterior Bayesian = 0.24, CIHDI95% [0.16, 0.33], rJZS Cauchy = 0.71
E
Posterior fit 8 changepoint
order along ancestral species mating chromsome
F
FWelch(7, 28.3) = 10.36, p = 2.21e-06, ωp2 = 0.64, CI95% [0.39, 1.00], nobs = 237
pHolm-adj. = 5.85e-03
pHolm-adj. = 3.85e-03
μmean = 7.30e-04
μmean = 4.88e-03
μmean = 3.81e-03
μmean = 0.05
μmean = 0.02
μmean = 0.02
μmean = 0.02
μmean = 0.01
strata1 (n = 27)
strata2 (n = 10)
strata3 (n = 7)
strata4 (n = 7)
strata5 (n = 28)
strata6 (n = 5)
strata7 (n = 49)
strata8 (n = 104)
eight_strata
Pairwise test: Games-Howell; Bars shown: significant
loge(BF01) = −2.00, R2posterior Bayesian = 0.07, CIHDI95% [0.00, 0.13], rJZS Cauchy = 0.71

**Figure S15: Changepoint inference without priors (A-C-E) and corresponding violin plots (B-D-F) displaying the distribution of $d_S$ values and significance of mean differences between evolutionary strata in *Microbotryum violaceum tatarinowii.*** Each changepoint panel displays the distribution of raw data (i.e., $d_S$ values as black dots) along with 25 draws from the joint posterior distribution (grey lines) and 95% highest density interval (red lines). Posterior distributions of the change points are shown in blue with one line for each chain. Posterior fits are displayed for models with 5, 6 and 7 changepoints, corresponding to 3, 4 and 5 evolutionary strata, respectively. The analysis was run without prior regarding the location of changepoints, based on observed discrete changes in the distribution of mean $d_S$ values. The violin-boxplots show the observed distribution of $d_S$ values for each evolutionary stratum, as colored points, with a box plot superimposed and a violin plot superimposed . Each read dot displays the mean of $d_s$ values along with its inferred value.

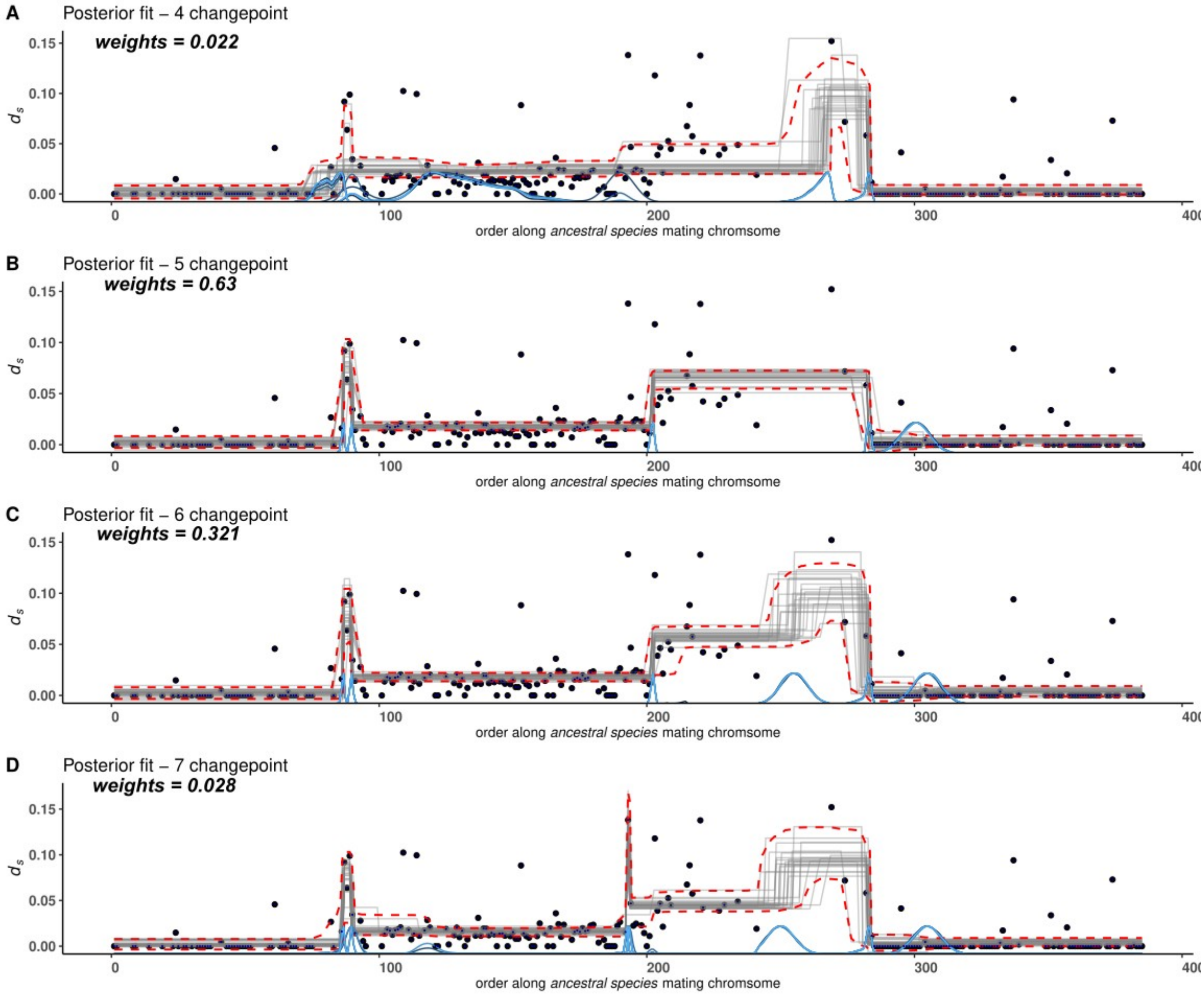


**Figure S16: Changepoint inference with priors in *Microbotryum violaceum tatarinowii*.** Each changepoint panel displays the distribution of raw data (i.e., $d_S$ values as black dots) along with 25 draws from the joint posterior distribution (grey lines) and 95% highest density interval (red lines). Posterior distributions of the change points are shown in blue with one line for each chain. Posterior fits are displayed for models with 5, 6 and 7 changepoints, corresponding to 3, 4 and 5 evolutionary strata, respectively.

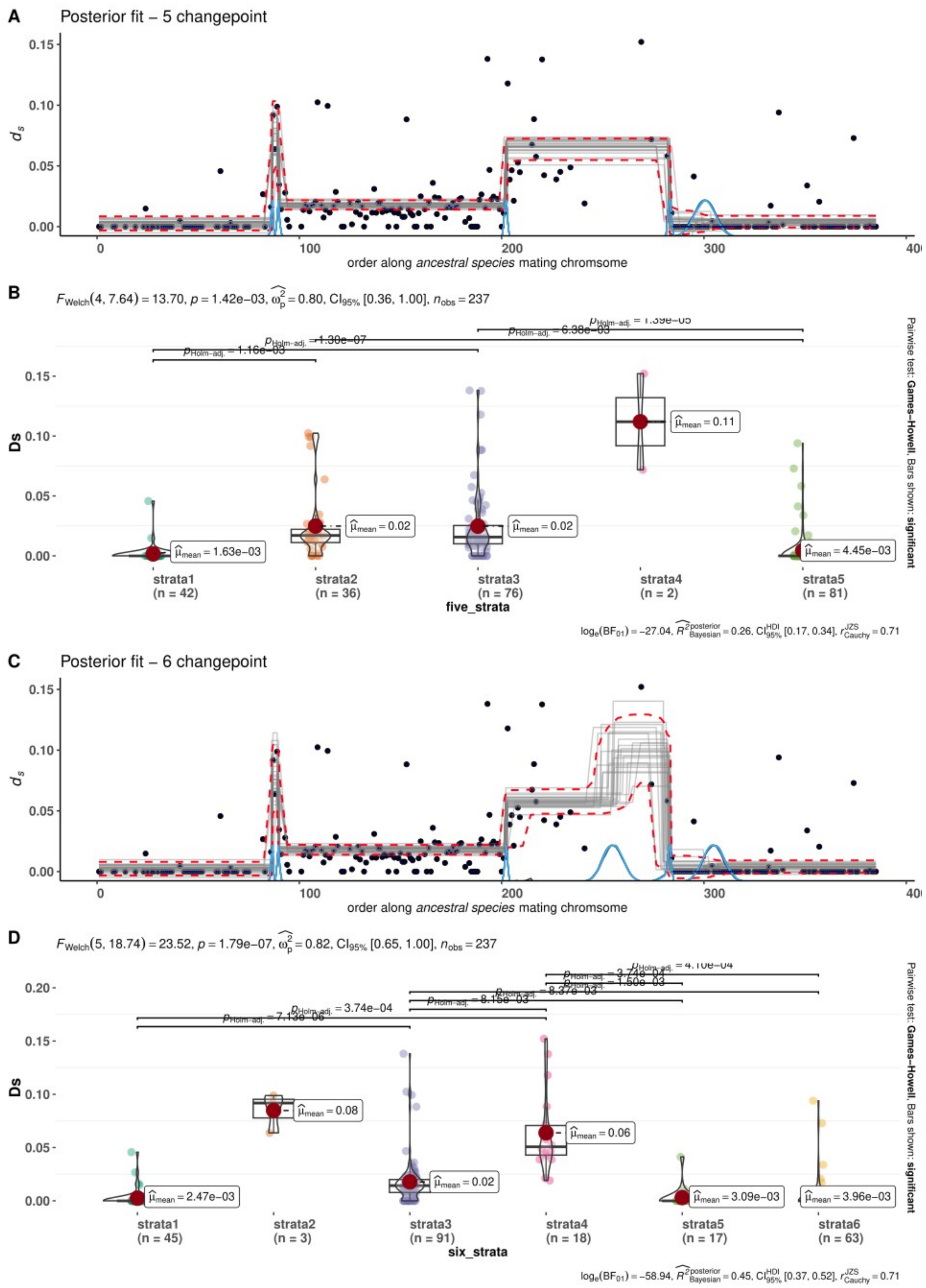


**Figure S17: Changepoint inference without priors (A-C) and corresponding violin plots (B-D) displaying the distribution of $d_S$ values and significance of mean differences between evolutionary strata in *Microbotryum violaceum tatarinowii*.** Each changepoint panel displays the distribution of raw data (i.e., $d_S$ values as black dots) along with 25 draws from the joint posterior distribution (grey lines) and 95% highest density interval (red lines).

Posterior distributions of the change points are shown in blue with one line for each chain. Posterior fits are displayed for models with 5 and 6 changepoints, corresponding to 4 and 5 evolutionary strata, respectively. The violin-boxplots show the observed distribution of $d_S$ values for each evolutionary stratum, as colored points, with a box plot superimposed and a violin plot superimposed. Each read dot displays the mean of $d_s$ values along with its inferred value.

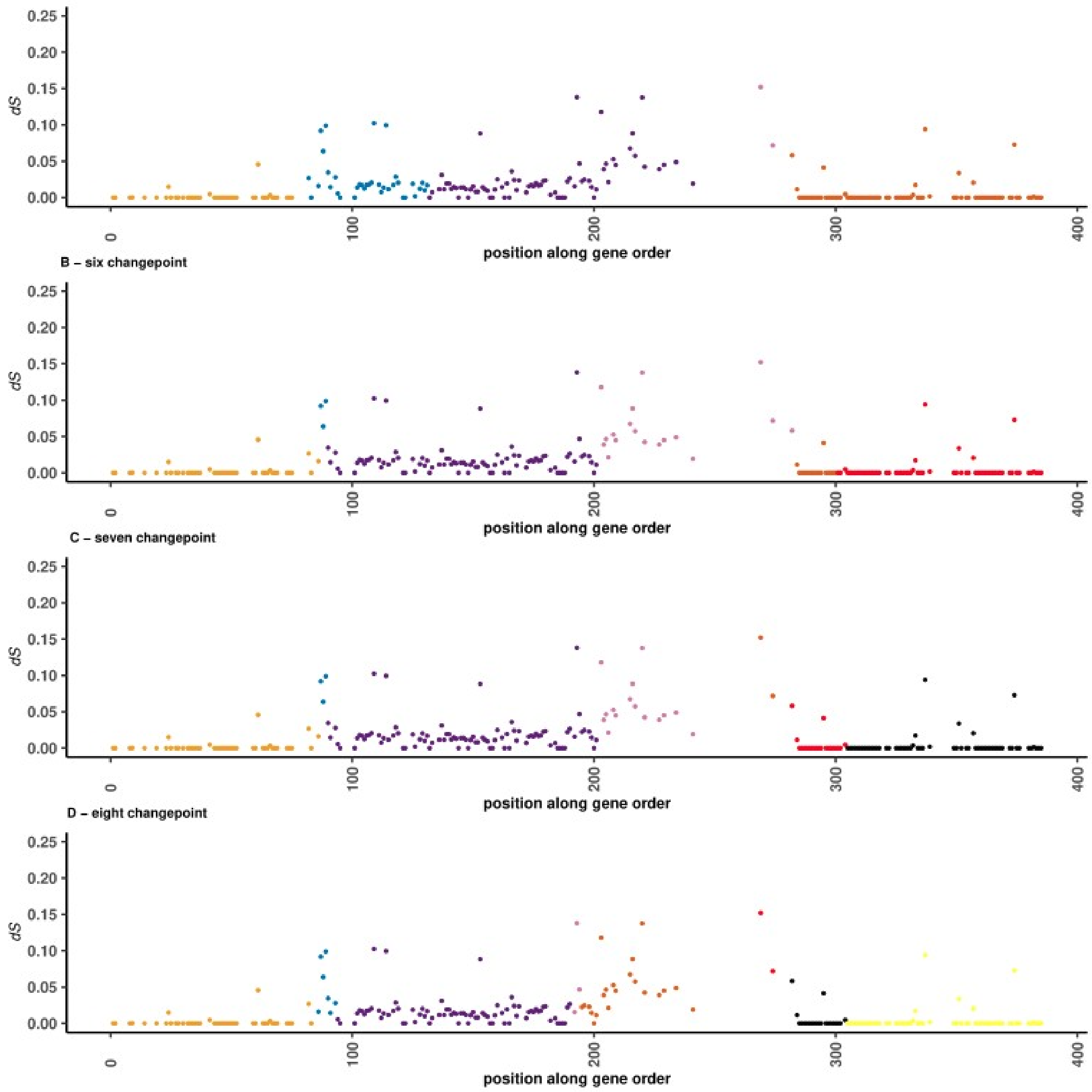


**Figure S18: Evolution of mating-type chromosomes in *Microbotryum violaceum tatarinowii.*** Localisation of the evolutionary strata in *M. v. tatarinowii* and their $d_S$ values along the ancestral gene order, using as proxy *M. lagerheimii.* Each point represents the value for a gene, assuming three (panel A, five changepoint) to five (panel D, eight changepoint) evolutionary strata color with $d_S$ values. The $d_S$ values at zero at the beginning and end of the plot correspond to the pseudo-autosomal regions (PARs) and do not correspond to evolutionary strata.

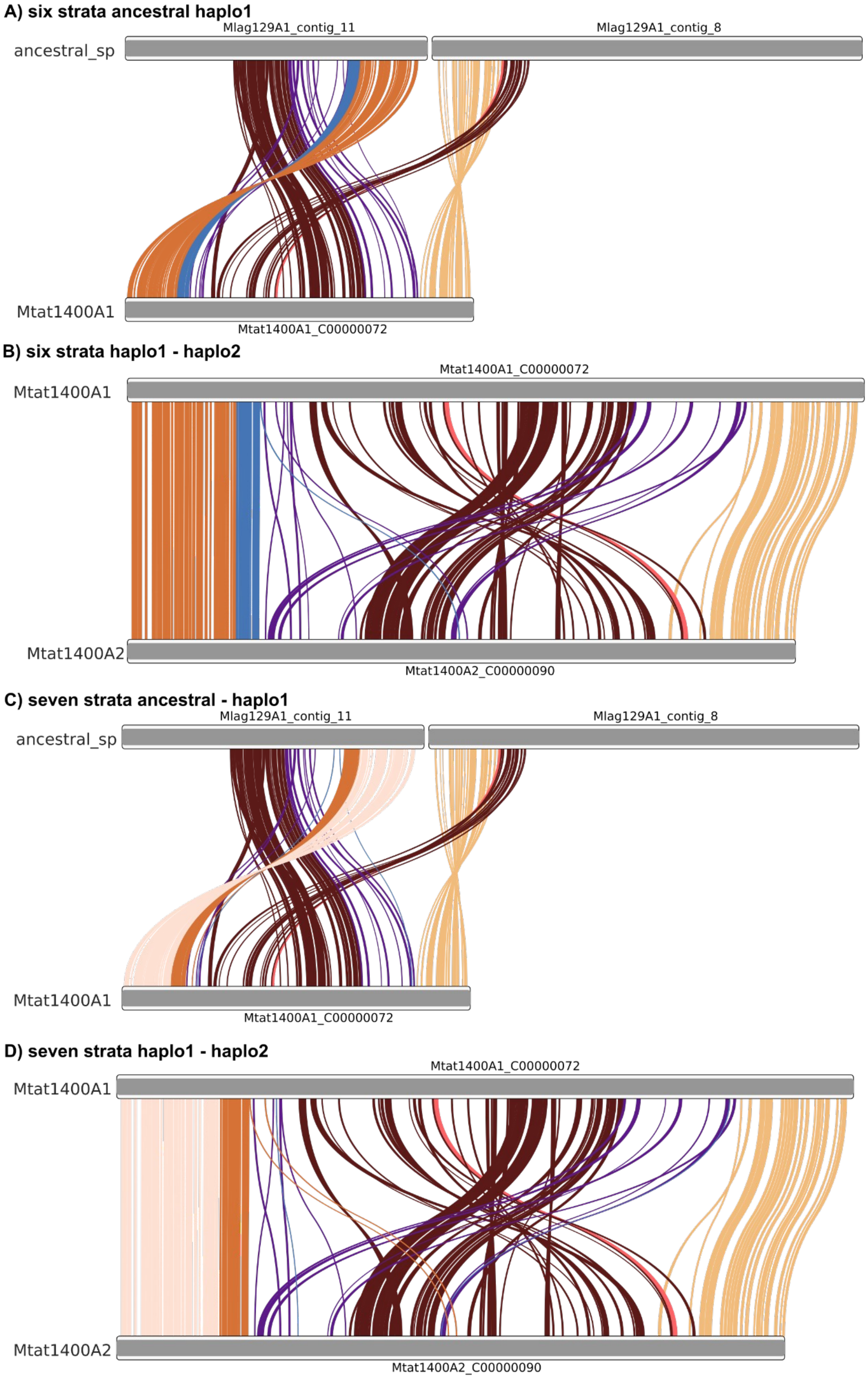


**Figure S19: Evolution of mating-type chromosomes in *Microbotryum violaceum tatarinowii.*** A) Ideograms showing links between single-copy orthologs genes of *Microbotryum violaceum tatarinowii* $a_1$ (Mtat1400A1) and either *M. v. caroliniana tatarinowii* $a_2$ *(Mtat1400A2)* or the genome used as a proxy for the ancestral gene order (*M. lagerheimii, labeled as "ancestral_sp"*). Evolutionary strata are colored, assuming four (panel A) to five strata (panel D) inferred a posteriori from the MCP analysis.

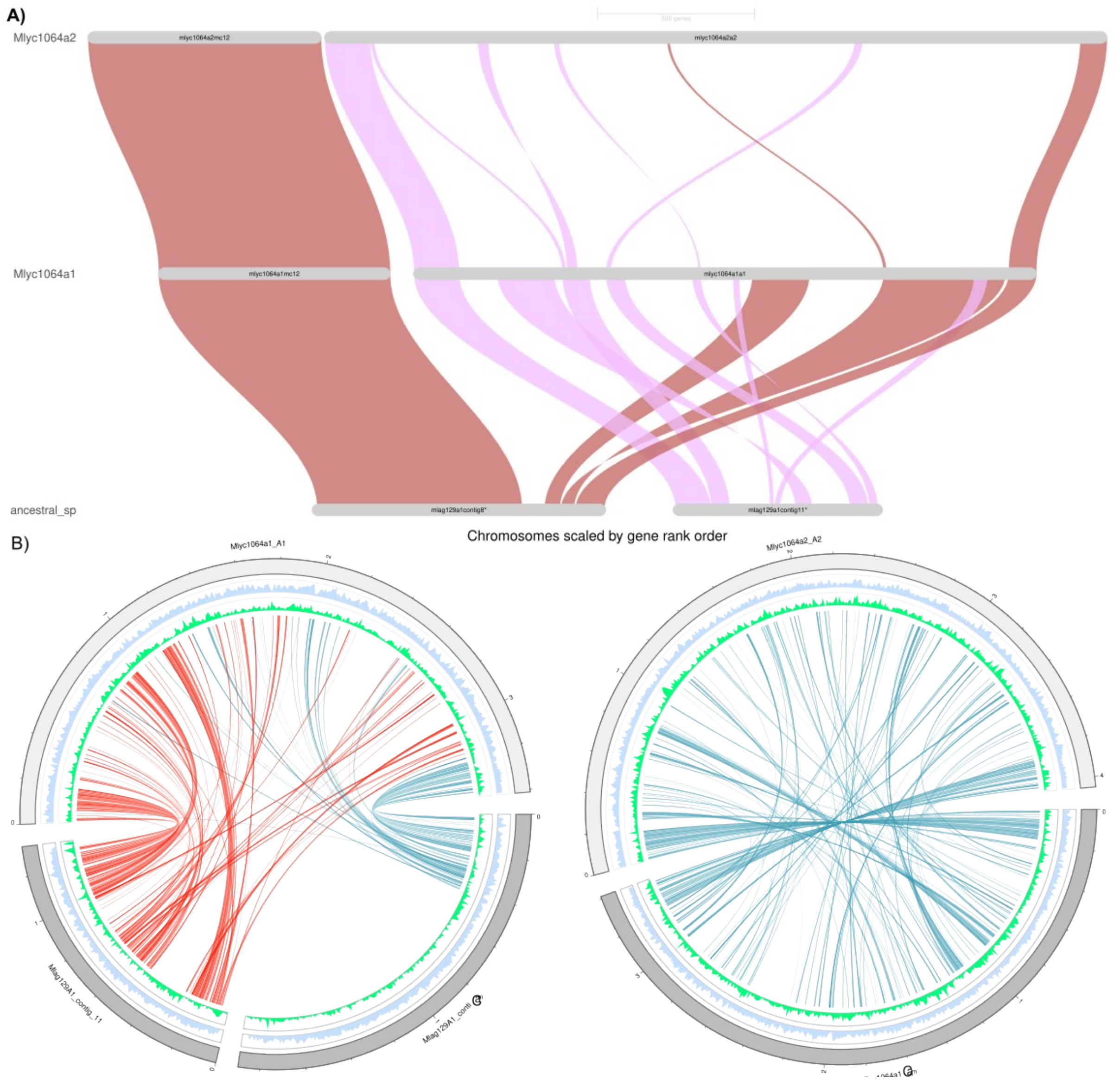


**Figure S20: GeneSpace (A) and Circos plots (B) showing synteny and orthology relationship between species and mating-type chromosomes of *Microbotryum lagerheimii* and *Microbotryum lychnidis-dioicae 1064* $a_1$ and between the two mating-type chromosomes of *M. lychnidis-dioicae* ($a_1$ and $a_2$). Mlag129A1_contig 8 corresponds to the HD chromosome and Malg129A1contig11 to the PR chromosome.**

**A)** Plots showing major synteny blocks as inferred by combining orthofinder, blasts and MCScan results. Synteny groups must include at least five consecutive genes.

**B)** Circos plots of the *M. lychnidis-dioicae $a_1$* mating-type chromosomes compared to the *M. lagerheimii* HD and PR mating-type chromosomes and *M. lychnidis-dioicae $a_1$* versus *M. lychnidis-dioicae $a_2$* mating-type chromosomes. Red link = single-copy orthologous genes between PR contigs. Blue links = single-copy orthologs genes between HD contigs.

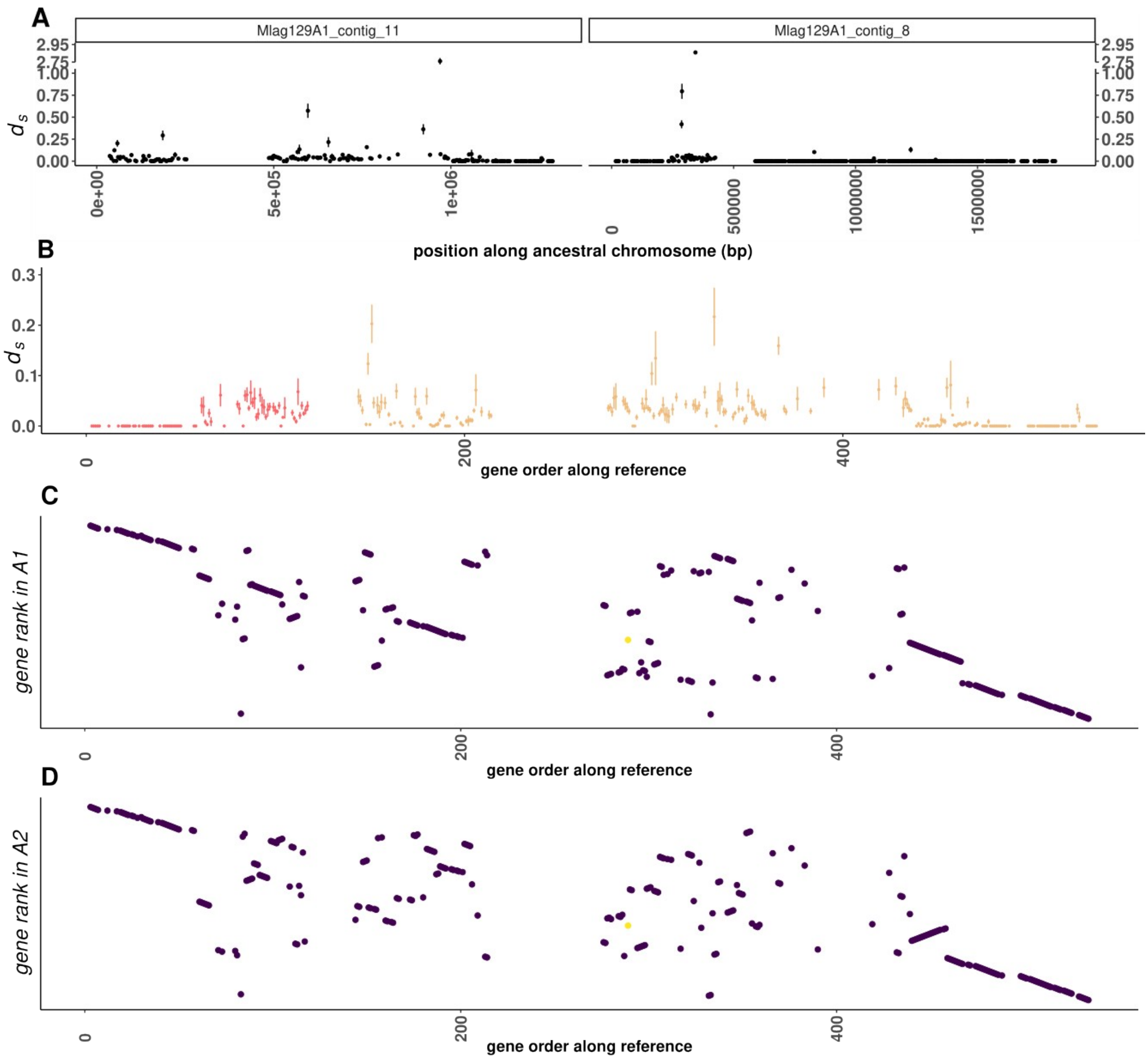


**Figure S21: Observed $d_S$ values and rearrangements in the *Microbotryum lychnidis-dioicae 1064* case. A)** observed $d_S$ values between the *Microbotryum lychnidis-dioicae 1064* $a_1$ and $a_2$ mating-type chromosomes along the mating-type chromosomes of *M. lagerheimii*, with the contig 11 corresponding to the PR chromosome and the contig 8 corresponding to the HD chromosome. B) Distribution of the $d_S$ values considering the ancestral gene order instead of the physical position along the *M. lagerheimii* mating-type chromosomes. The large region in contig 8 (HD, red) that became an autosome upstream of the centromere in panel A (>500 kbp) has been removed. Contig 11 (PR, yellow) is placed upstream of HD. C) and D) Current distribution of the rank of the genes in the $a_1$ and $a_2$ mating-type chromosomes along the ancestral gene order, showing the rearrangements.

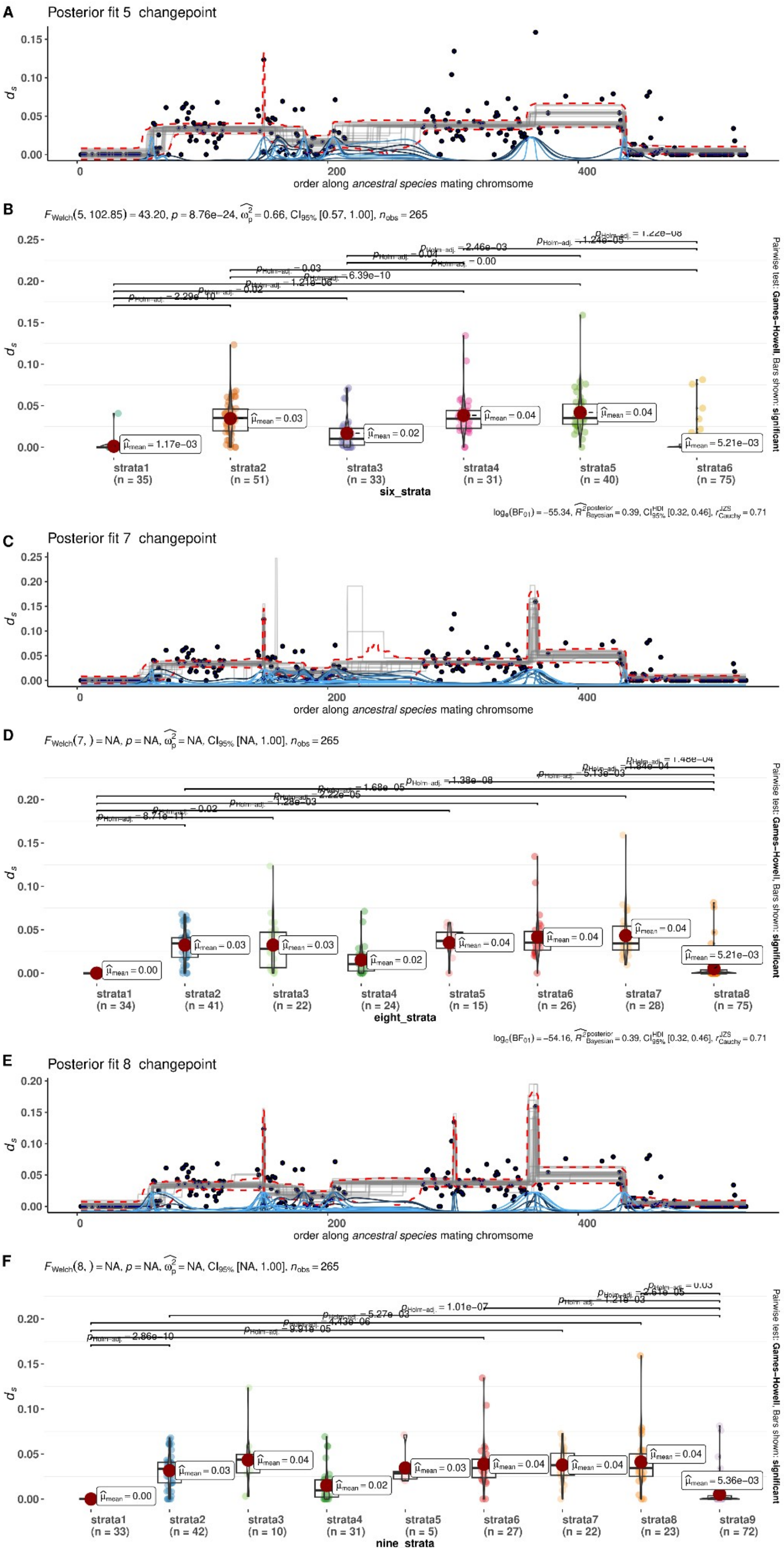
A
Posterior fit 5 changepoint
order along ancestral species mating chromsome
B
FWelch(5, 102.85) = 43.20, p = 8.76e-24, ω̂p2 = 0.66, CI95% [0.57, 1.00], nobs = 265
strata1 (n = 35)
strata2 (n = 51)
strata3 (n = 33)
strata4 (n = 31)
strata5 (n = 40)
strata6 (n = 75)
six_strata
log(BF01) = −55.34, R̂2posterior Bayesian = 0.39, CIHDI 95% [0.32, 0.46], rJZS Cauchy = 0.71
Pairwise test: Games-Howell; Bars shown: significant
C
Posterior fit 7 changepoint
order along ancestral species mating chromsome
D
FWelch(7, ) = NA, p = NA, ω̂p2 = NA, CI95% [NA, 1.00], nobs = 265
strata1 (n = 34)
strata2 (n = 41)
strata3 (n = 22)
strata4 (n = 24)
strata5 (n = 15)
strata6 (n = 26)
strata7 (n = 28)
strata8 (n = 75)
eight_strata
log(BF01) = −54.16, R̂2posterior Bayesian = 0.39, CIHDI 95% [0.32, 0.46], rJZS Cauchy = 0.71
Pairwise test: Games-Howell; Bars shown: significant
E
Posterior fit 8 changepoint
order along ancestral species mating chromsome
F
FWelch(8, ) = NA, p = NA, ω̂p2 = NA, CI95% [NA, 1.00], nobs = 265
strata1 (n = 33)
strata2 (n = 42)
strata3 (n = 10)
strata4 (n = 31)
strata5 (n = 5)
strata6 (n = 27)
strata7 (n = 22)
strata8 (n = 23)
strata9 (n = 72)
nine_strata
Pairwise test: Games-Howell; Bars shown: significant

**Figure S22: Changepoint inference without priors (A,C,E) and corresponding violin plots (B,D,F) displaying the distribution of $d_S$ values and significance of mean differences between evolutionary strata in *Microbotryum lychnidis-dioicae 1064*.** Each changepoint panel displays the distribution of raw data (i.e., $d_S$ values as black dots) along with 25 draws from the joint posterior distribution (grey lines) and 95% highest density interval (red lines). Posterior distributions of the change points are shown in blue with one line for each chain. Posterior fits are displayed for models with 5 and 6 changepoints, corresponding to 4 and 5 evolutionary strata, respectively. The violin-boxplots show the observed distribution of $d_S$ values for each evolutionary stratum, as colored points, with a box plot superimposed and a violin plot superimposed. Each read dot displays the mean of $d_s$ values along with its inferred value.

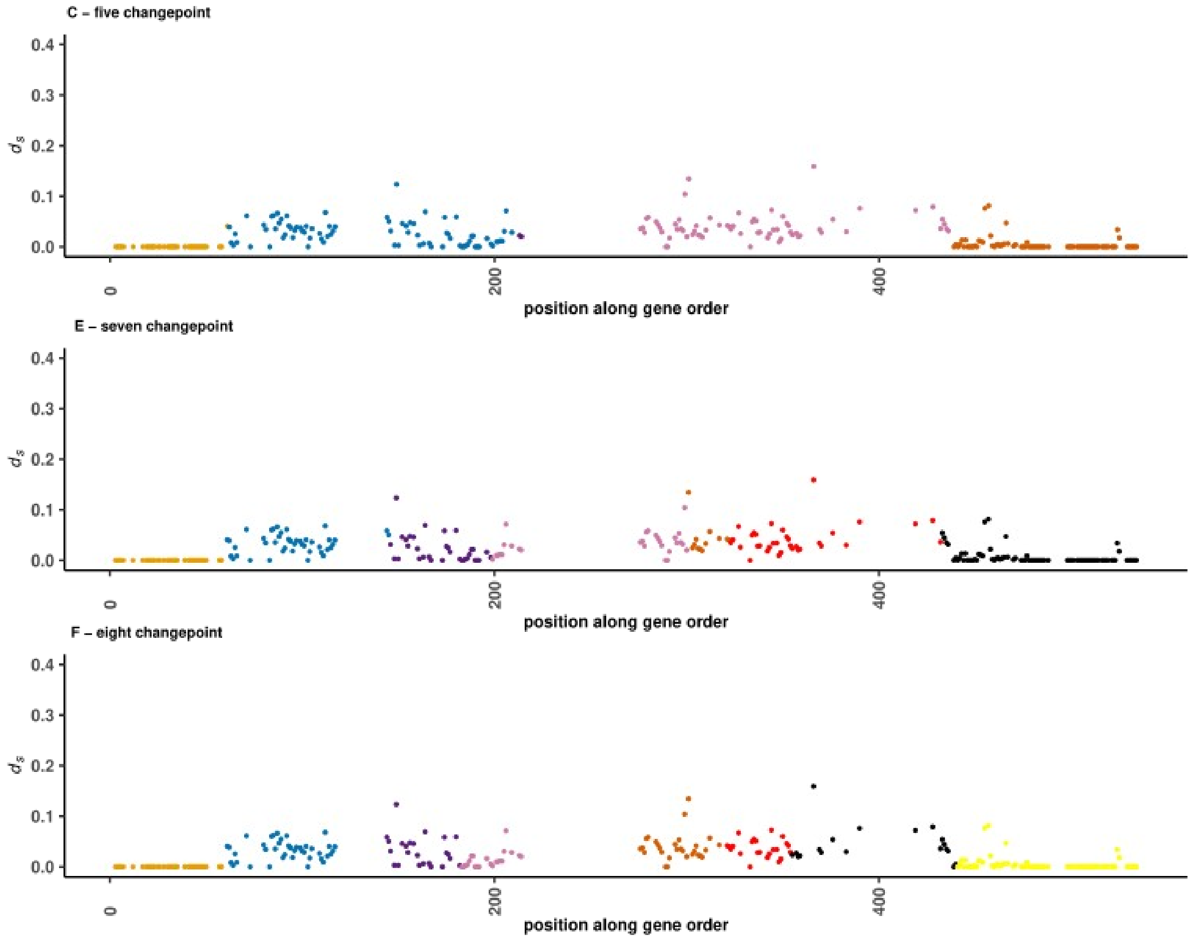


**Figure S23: Evolution of mating-type chromosomes in *Microbotryum lychnidis-dioicae 1064*.** Localisation of the evolutionary strata in *Microbotryum lychnidis-dioicae* and their $d_S$ values along the ancestral gene order, using as proxy *M. lagerheimii.* Each point represents the value for a gene, assuming three (panel A, five changepoint) to five (panel D, eight changepoint) evolutionary strata color with $d_S$ values. The $d_S$ values at zero at the beginning and end of the plot correspond to the pseudo-autosomal region (PARs) and not to evolutionary strata.

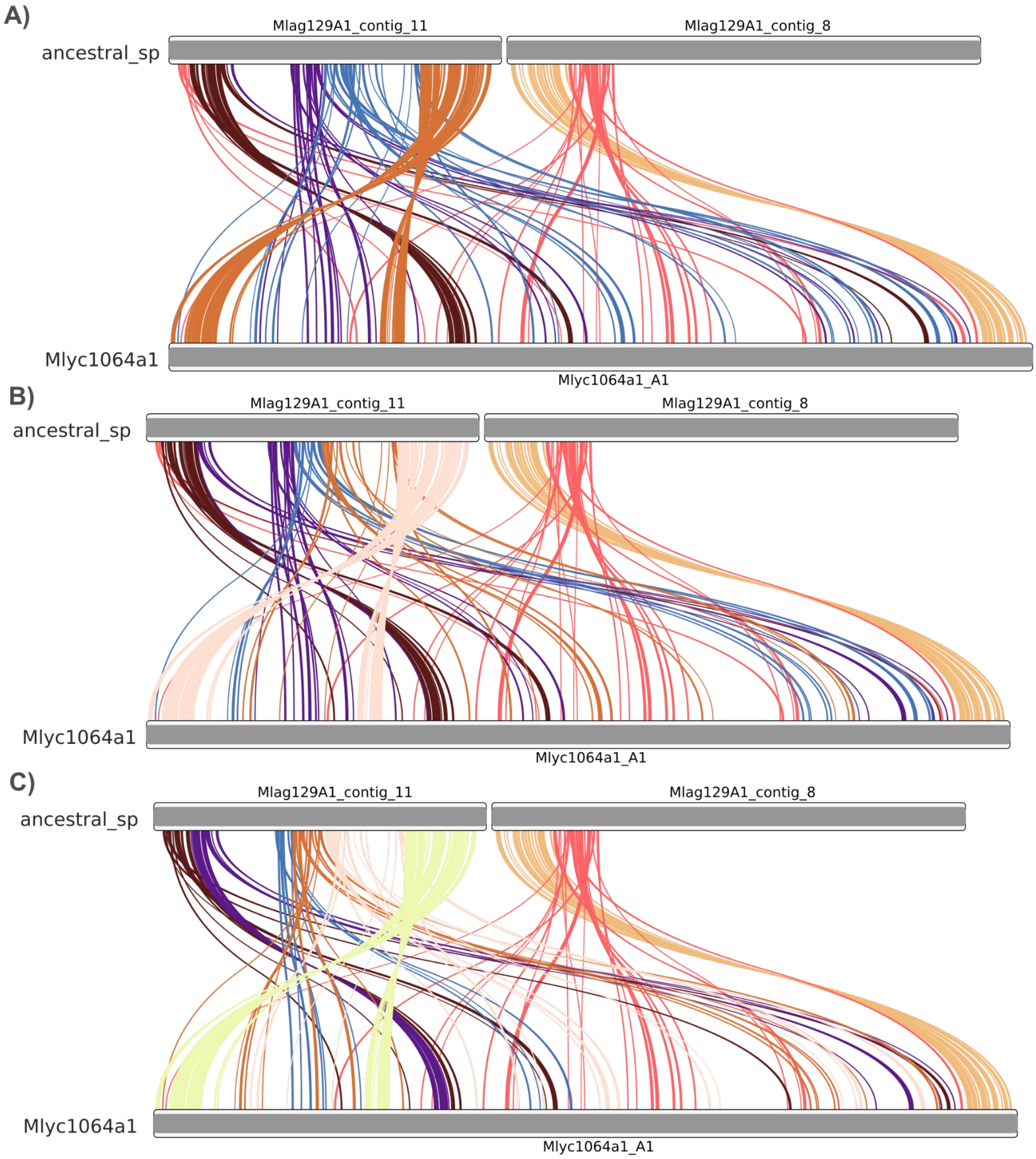

A)
ancestral_sp
Mlag129A1_contig_11
Mlag129A1_contig_8
Mlyc1064a1
Mlyc1064a1_A1
B)
ancestral_sp
Mlag129A1_contig_11
Mlag129A1_contig_8
Mlyc1064a1
Mlyc1064a1_A1
C)
ancestral_sp
Mlag129A1_contig_11
Mlag129A1_contig_8
Mlyc1064a1
Mlyc1064a1_A1

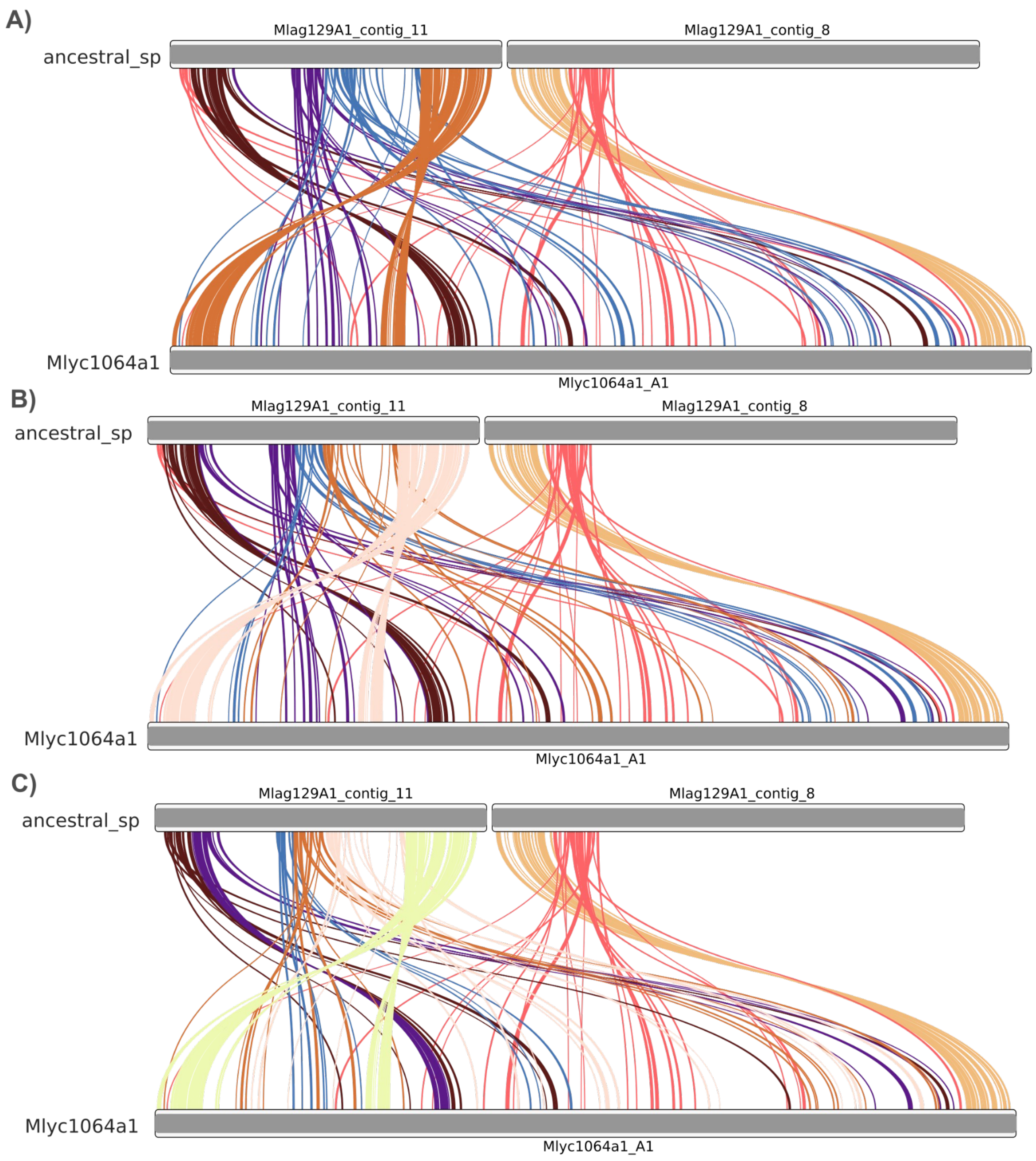


**Figure S24: Evolution of mating-type chromosomes in *Microbotryum lychnidis-dioicae 1064.*** A) Ideograms showing links between single-copy orthologs genes of *M. lychnidis-dioicae 1064* $a_1$ and the genome used as a proxy for the ancestral gene order (*M. lagerheimii*). Evolutionary strata are colored, assuming four (panel A) five (panel B) or six strata (panel C) inferred a posteriori from the MCP analysis.

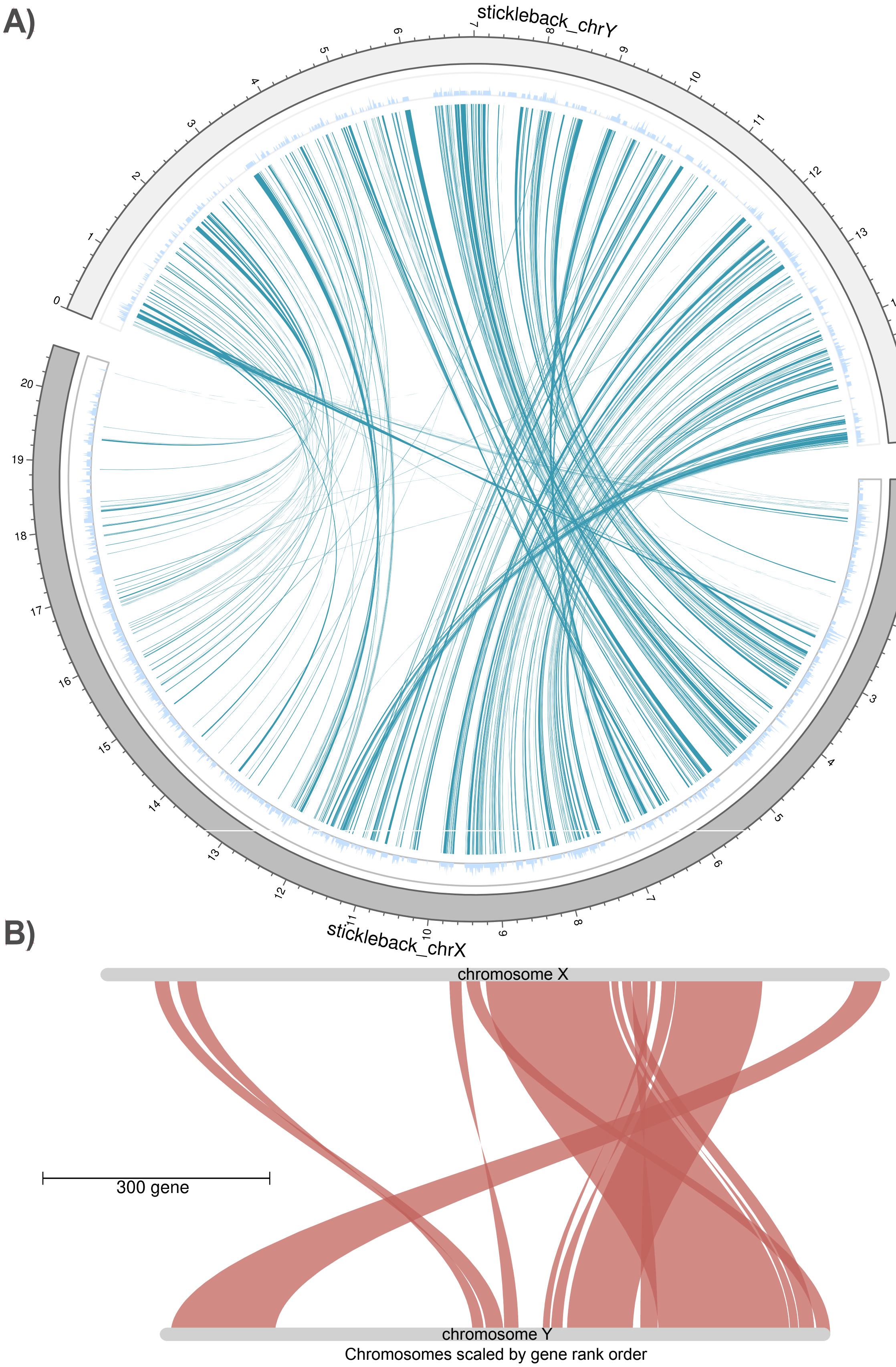


**Figure S25:** Circos plots (A) and GeneSpace (B) showing synteny and orthology relationships between X and Y chromosome of the threespine stickleback A) Plots showing major synteny blocks as inferred by combining orthofinder, blasts and MCScanX results. B)

Circos plots of the X and Y chromosomes

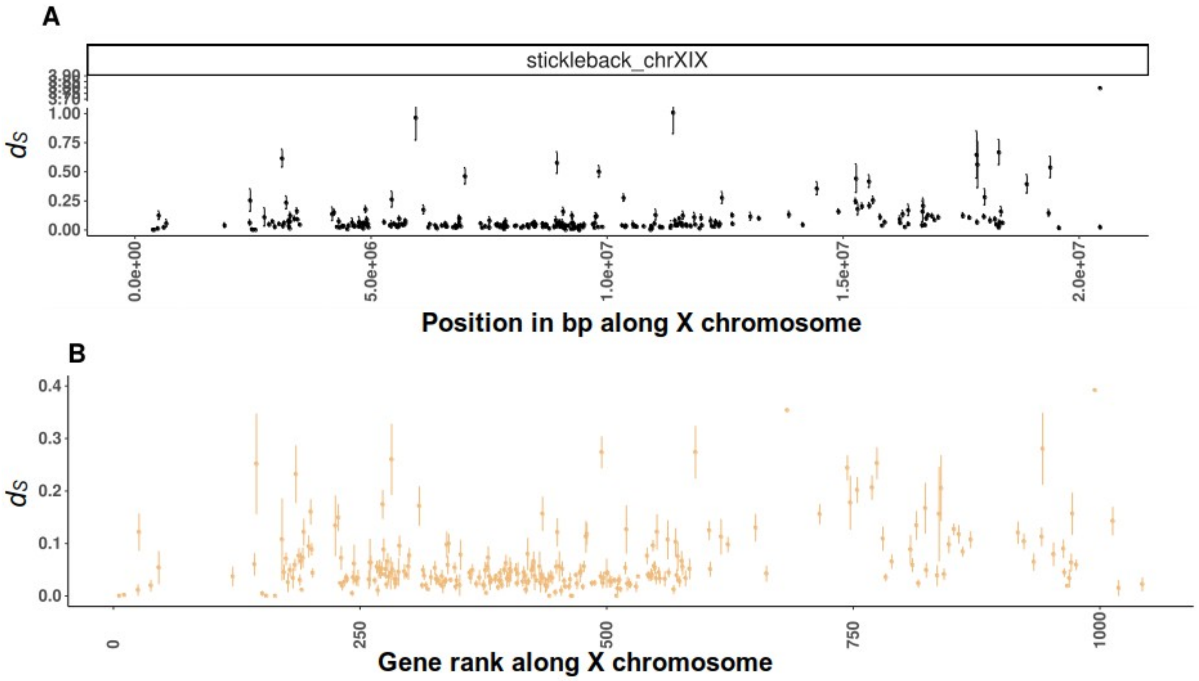


**Figure S26:** visualization of $d_S$ values between single copy orthologs of X and Y chromosomes plotted along the X chromosome of the stickleback. A) $d_S$ values plotted along the genome in base pair. B) $d_S$ values plotted along the gene rank

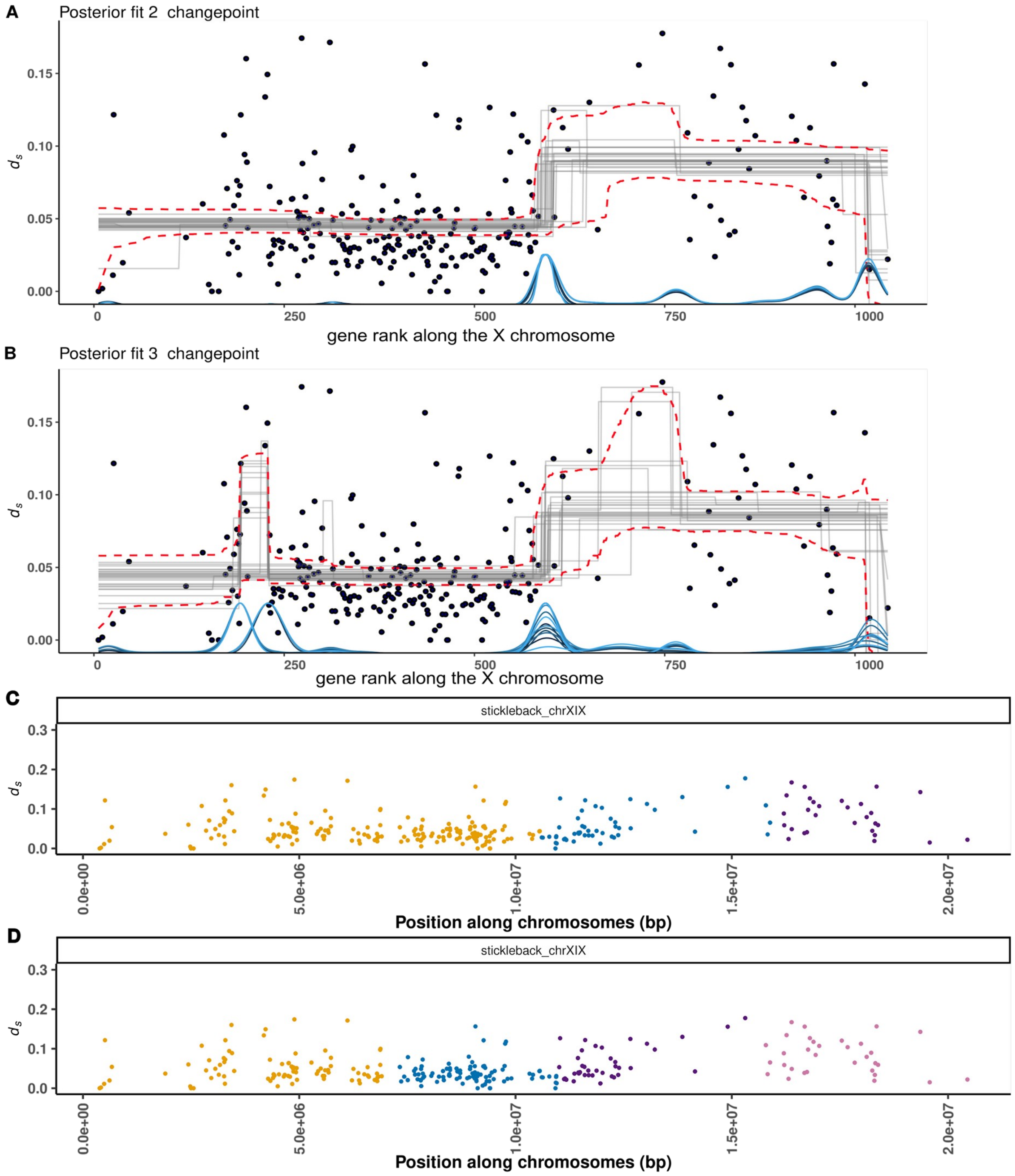


**Figure S27:** Changepoint analysis recovers the three major evolutionary strata identified previously in Peichel et al. 2020 (see Figure 4 in Peichel). Each changepoint panel (A and B) displays the distribution of raw data (i.e., $d_S$ values as black dots) along with 25 draws from the joint posterior distribution (grey lines) and 95% highest density interval (red lines). Posterior distributions of the changepoints are shown in blue with one line for each chain.
C) and D) displays the corresponding location of the strata where each point represents a gene colored according to the inferred strata by the changepoint analysis with 2 and 3 changepoint respectively.

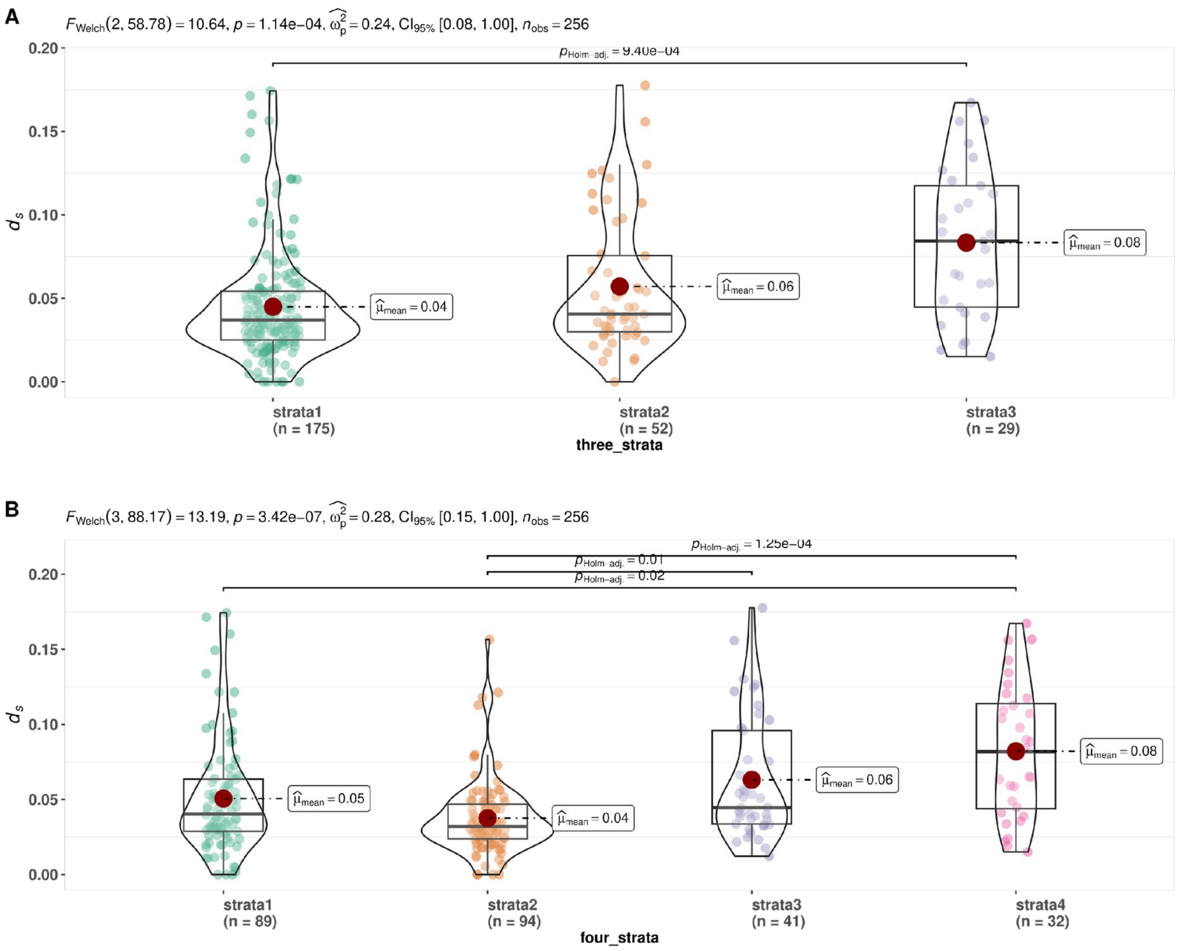


**Figure S28:** violin-boxplot displaying the distribution of $d_S$ values for each of the strata according to the 2 changepoint model (A) and 3 changepoint model (B). Significance of the difference are displayed. Each red point represents the mean dS value per strata

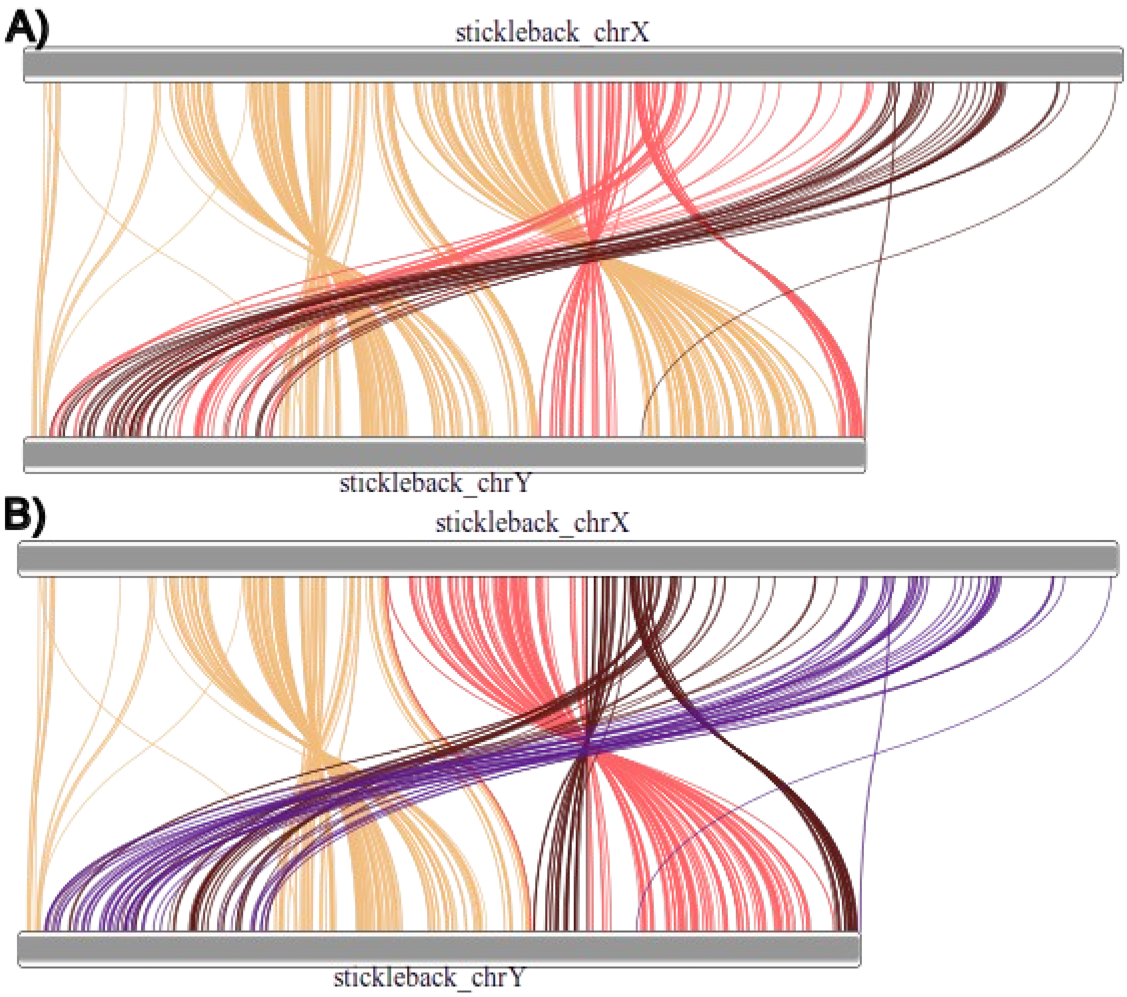


**Figure 29:** Evolution of X and Y chromosomes strata in threespine stickleback A) Ideograms showing links between single-copy orthologs genes of X and Y chromosomes. Evolutionary strata are colored, assuming three (panel A) or four strata (panel B) inferred a posteriori from the MCP analysis.

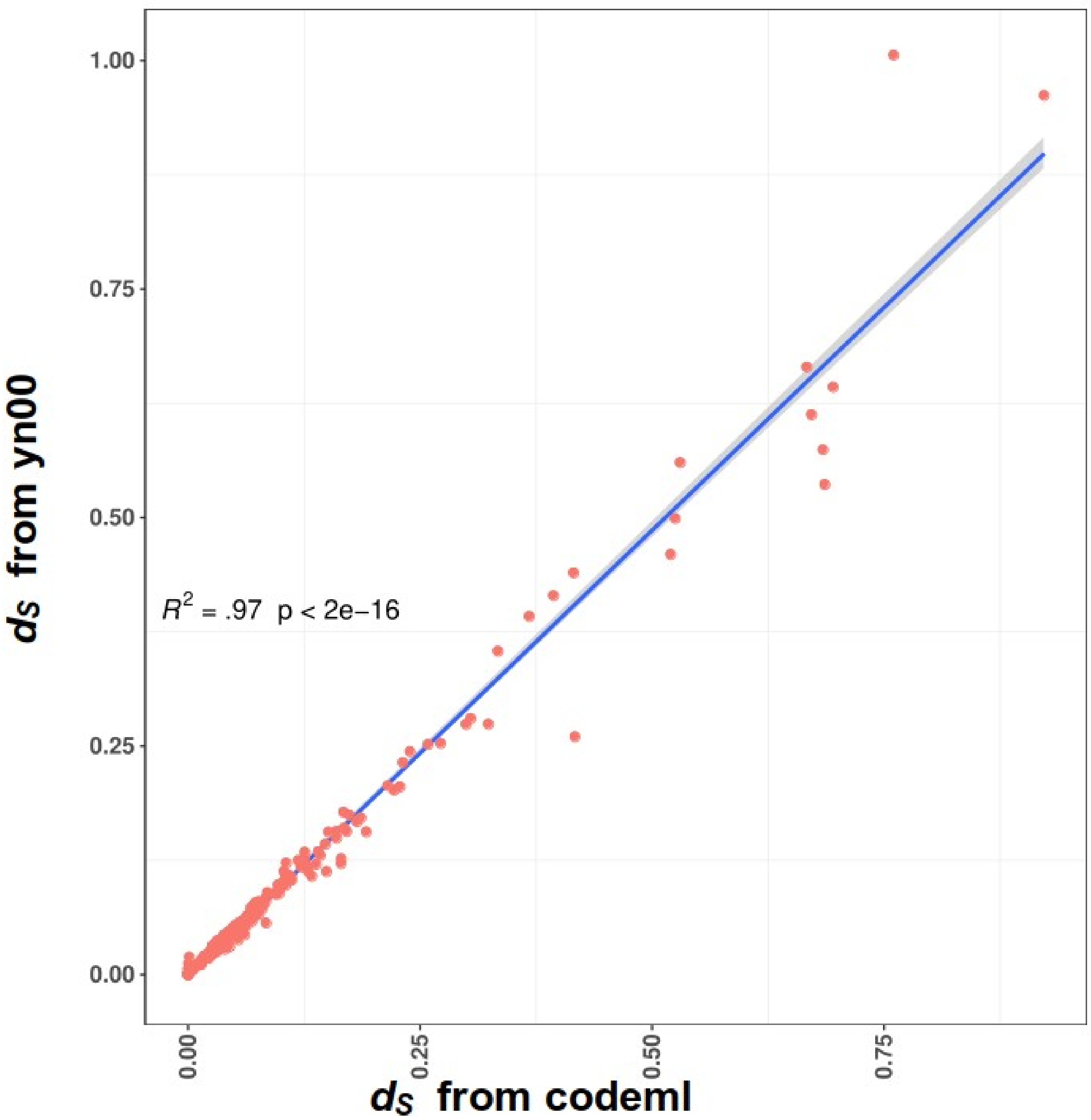


**Figure S30:** Correlation between $d_S$ values obtained from codeml (x-axis) and yn00 (y-axis). Each red dot represents a $d_S$ value computed for the same pair of gene by the two methods. The blue line represents the regression line from the lm() approach implemented in the R package ggplot2. The grey area represents the 95% confidence intervals computed around the regression mean.
